# Expression of nano-engineered RNA organelles in bacteria

**DOI:** 10.1101/2025.07.08.663582

**Authors:** Brian Ng, Catherine Fan, Milan Dordevic, Adam Knirsch, Layla Malouf, Giacomo Fabrini, Sabrina Pia Nuccio, Roger Rubio-Sánchez, Graham Christie, Masahiro Takinoue, Pietro Cicuta, Lorenzo Di Michele

**Affiliations:** Department of Chemical Engineering and Biotechnology, University of Cambridge, Cambridge, United Kingdom; Cavendish Laboratory, Department of Physics, University of Cambridge, Cambridge, United Kingdom; Department of Chemistry, Imperial College London, London, United Kingdom; The Francis Crick Institute, London, United Kingdom; Department of Computer Science, Institute of Science Tokyo, Tokyo, Japan; Research Center for Autonomous Systems Materialogy (ASMat), Institute of Integrated Research (IIR), Institute of Science Tokyo, Tokyo, Japan; fabriCELL, Imperial College London, London, United Kingdom

**Author notes:** These authors contributed equally to this work (Brian Ng, Catherine Fan, Milan Dordevic, Adam Knirsch).

**Keywords:** RNA Nanotechnology, Biomolecular Condensates, Membrane-less Organelles, Synthetic Biology, E. coli

## Abstract

Designing synthetic biomolecular condensates, or membrane-less organelles, offers insights on the functions of their natural counterparts, and is equally valuable for cellular and metabolic engineering. Choosing E. coli for its biotechnological relevance, we deploy RNA nanotechnology to design and express non-natural membrane-less organelles in vivo. The designer condensates assemble co-transcriptionally from branched RNA motifs interacting via base-pairing. Exploiting binding selectivity we express orthogonal, non-mixing condensates, and by embedding a protein-binding aptamer we achieve selective protein recruitment. Condensates can be made to dissolve and re-assemble upon thermal cycling, thereby reversibly releasing and re-capturing protein clients. The synthetic organelles are expressed robustly across the cell population and remain stable despite enzymatic RNA processing. Compared with existing solutions based on peptide building blocks or repetitive RNA sequences, these nanostructured RNA motifs enable algorithmic control over interactions, affinity for clients, and condensate microstructure, opening new directions in synthetic biology and biotechnology.

## Introduction

Membraneless organelles (MLOs), emerging from condensation of proteins and nucleic acids, are found in most living cells, where they orchestrate biochemical reactions by sequestering and co-localising enzymes, substrates, and metabolites, and by maintaining homeostasis [1–6]. Historically thought not to contain intracellular compartments, prokaryotes are now recognised to leverage membraneless organelles, such as BR-bodies, carboxysomes, polyP granules, ParB condensates, and RNA polymerase clusters, to modulate processes including division, polarisation, nucleoid replication, transcription, and protein homeostasis [5–8].

Engineering non-natural, *designer* MLOs both in vitro and in vivo is key to unravelling the biophysics and functions of natural condensates [9–11], and equally valuable as a tool to program metabolism and cellular functions in synthetic biology and biomanufacturing [12, 13]. To this end, establishing robust strategies to engineer biomolecular condensation in widely adopted prokaryotic chassis is particularly urgent.

Synthetic MLOs have been constructed from peptides and proteins, demonstrating remarkable functionalities [14–28]. Condensate-forming proteins, however, often assemble through non-specific interactions between Intrinsically Disordered Regions (IDRs), commonly combining electrostatic, *π* − *π* and hydrophobic interactions between amino acid residues [29, 30]. The complexity of these interactions makes them difficult to prescribe by design [31–33], with modelling often requiring advanced numerical simulations [34–36].

Alternative to proteins, naturally occurring RNA repeats can be deployed for MLO assembly [37–39], but their simple structure limits the scope for more advanced engineering.

Different from proteins, however, nucleic acids benefit from highly predictable base-pairing interactions, which have led to the emergence of robust, algorithmic strategies to design nanostructures and materials [40–44]. DNA nanotechnology has been successfully deployed to construct synthetic condensates with precisely controlled size and microstructure [45–50], and displaying advanced functions ranging from reversible assembly and disassembly [51–53], to large-scale morphological restructuring [54–60], capture and release of functional payloads [51, 61, 62], and spatial organization of biocatalytic pathways [63–66]. Despite this versatility, the use of synthetic DNA building blocks is limited to in vitro applications, as DNA nanostructures cannot be expressed by living cells [67].

To overcome this roadblock, here we deploy RNA nanostructures, which retain much of the programmability of DNA constructs [68–71], but can be synthesised in vivo [72] and in crowded environments that mimic cellular conditions [73]. We engineer *E. coli* to express branched RNA motifs that assemble into synthetic MLOs when featuring complementary base-pairing domains, while remaining largely soluble when designed to be non-interacting. Base-pairing selectivity enables the expression of orthogonal nanostructures, producing non-mixing, co-existing MLOs, while embedding aptamers facilitates specific protein capture. MLOs can be made to assemble and disassemble upon thermal cycling, reversibly releasing and re-capturing payloads in the process. Our results demonstrate the potential of co-transcriptional RNA nanotechnology for engineering non-natural MLOs, combining a robust and programmatic design paradigm with genetic encodability.

## Results and discussion

### Design and expression of condensate-forming RNA motifs in *E. coli*

The condensate-forming motifs consist of branched RNA *nanostars* (NS), with four double stranded (ds) ‘arms’ emanating from a central junction (Fig. 1A) [69]. Each nanostar arm terminates with a 9-nucleotide (nt) kissing loop (KL), consisting of a 6 nt recognition domain asymmetrically flanked by adenines (Fig. 1A). NS designs A and B feature palindromic, self-complementary domains in the KLs, producing affinity between same-type NSs and leading to condensation [69]. A and B KLs are mutually orthogonal resulting in negligible cross-binding [69]. NS designs Ā and B̄ are identical to A and B, respectively, but have KL sequences scrambled to eliminate self-complementarity and prevent condensation [69]. Fluorescent Light-Up Aptamers (FLAPs) are embedded in one of the NS arms to facilitate detection and visualisation. Designs A and Ā include a ‘Pepper aptamer’ binding the HBC620 dye [74] (Fig. 1A(i) and Supplementary Fig. 1), while B and B̄ feature a ‘Broccoli aptamer’ [77] binding the TBI dye (Fig. 1A(ii) and Supplementary Fig. 1) [76]. All NS sequences are reported in Supplementary Table 1.

**Fig. 1:**
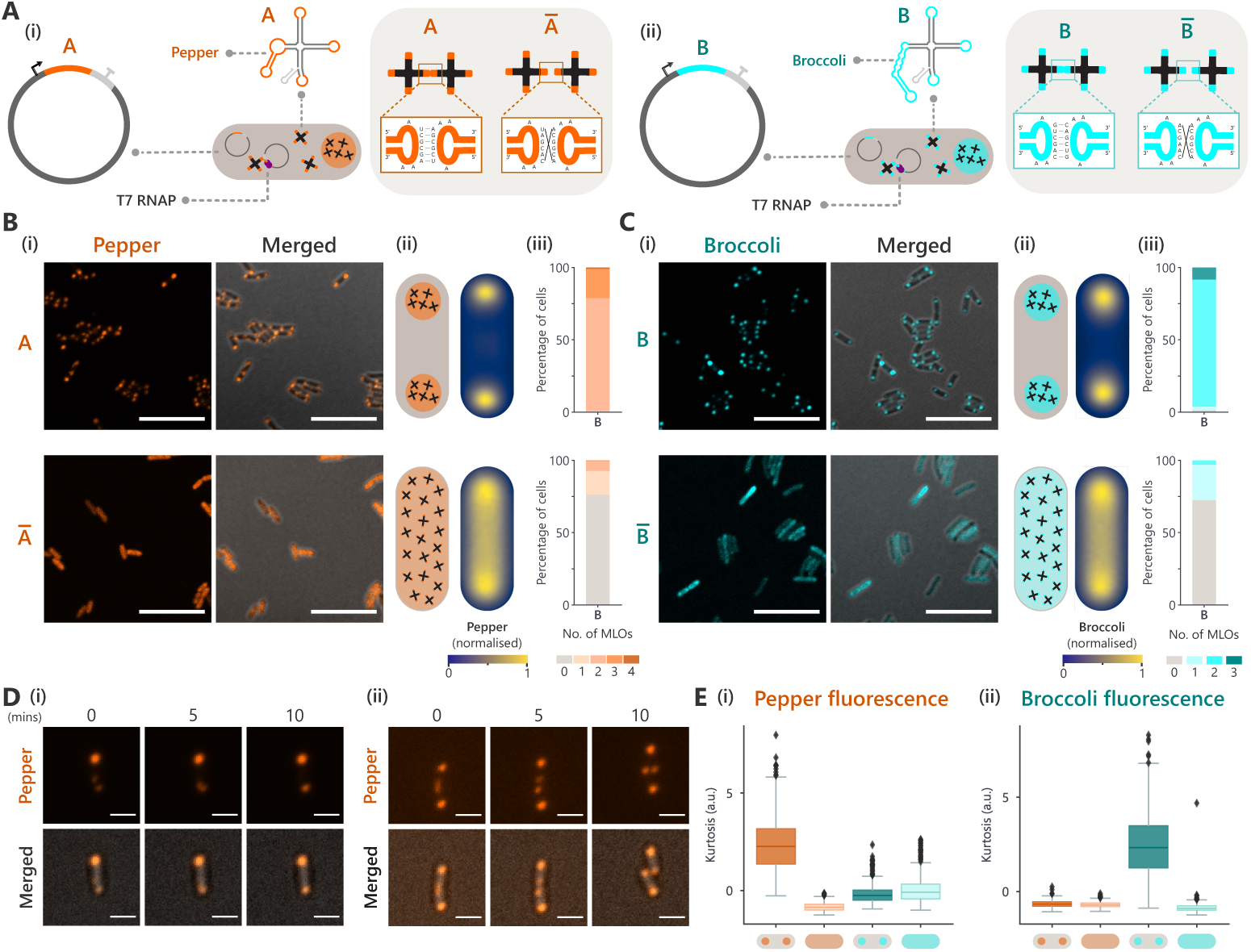
Expression of nanostructured RNA MLOs in *E. coli*. (**A**) Schematic of nanostar (NS) expression and membrane-less organelle (MLO) formation in *E. coli*. (i) NS A contains four self-complementary kissing loops (KLs), driving condensation. NS Ā has non-complementary KLs with scrambled sequences, preventing assembly. Both A and Ā contain a Pepper aptamer, which binds to HBC620 activating its fluorescence [74]. (ii) NS B contains four self-complementary KLs, with a different sequence from that in A. B̄ has non-complementary loops and does not form MLOs. Both B and B̄ include a Broccoli aptamer [75], which binds TBI and activates its fluorescence [76]. Each NS sequence is cloned into a plasmid for expression with T7 RNA polymerase (T7 RNAP) upon transformation into *E. coli*. A terminator sequence (gray loop) is included. (**B**) Expression of A (top) and Ā (bottom) in *E. coli*. (i) Confocal micrographs. Left: Pepper fluorescence. Right: Merged Pepper fluorescence with bright-field. Scale bars, 10 µm. (ii) Schematic representation (left) and population-averaged maps (right) of Pepper fluorescence distribution in *E. coli* (Methods). (iii) Percentage of *E. coli* containing 0, 1, 2, 3 or 4 MLOs per cell. (**C**) Data as for panel **B**, but for the expression of B (top) and B̄ (bottom) NSs in *E. coli*. (**D**) Time-lapse confocal micrographs of *E. coli* expressing A MLOs. Top: Pepper fluorescence. Bottom: Merged Pepper and bright-field fluorescence. Time is shown relative to the first frame (0 min). (i) MLOs fusion events within cells. Scale bars, 2 µm. (ii) Cell division in the presence of MLOs. Scale bars, 2 µm. (**E**) Box plots of the values of kurtosis extracted from the pixel-value distribution of fluorescence intensity in single cells. Analysis is performed on (i) Pepper and (ii) Broccoli fluorescence for *E. coli* expressing A, Ā, B and B̄ NSs (from left to right). Outcomes of statistical test are provided in Tables 4 and 5. In **B**, **C** and **E**, data were obtained from 3 separate cultures, each starting from a single colony.

For all NS designs, correct expression, condensate formation (or lack thereof) and interaction orthogonality were confirmed with in vitro transcription experiments (Extended Data Fig. 1 and Supplementary Figs. 2, 3, 4, 5, 6, 7, Methods). Encouragingly for in vivo expression, robust condensation was observed in buffers mimicking the intracellular ionic conditions encountered in *E. coli* (200 mM K^+^ and 1 mM Mg^2+^ [78, 79]) (Supplementary Fig. 8).

NS sequences, followed by transcription termination domains [80, 81], were then cloned in plasmids for expression in the *E. coli* BL21(DE3) strain with the IPTG-inducible T7 transcription system, as outlined in the Methods. Plasmid maps and sequences are reported in Supplementary Figs. 9, 10 and Supplementary Table 2. *E. coli* were grown from a single colony in supplemented M9 medium to mid-log phase (OD 0.7) before inducing NS expression. Cells were characterised by confocal microscopy one hour post-induction (Methods).

We first tested *E. coli* expressing individual NSs: A, B, Ā or B̄. For both “sticky” motifs, A and B, visual inspection confirms MLO formation (Fig. 1B(i)-C(i) and Supplementary Figs. 11, 12). Condensates appear to be overwhelmingly located at the poles of the cell, a common pattern in protein MLOs [8]. This observation is confirmed by averaging the fluorescence distributions of individual cells across the populations, producing the 2D maps in Fig. 1B(ii)-C(ii). Pole localisation is likely to result from steric hindrance by the nucleoid located in the central section of the cell [82]. To test this hypothesis, in Supplementary Note 1, we use simple polymer physics arguments to estimate the energy cost of forming a small condensate (50 nm in diameter) within the nucleoid [83]. We find this penalty to be as high as ∼130 *k*_B_*T*, equivalent to ∼ 500× the KL hybridisation free energy, which justifies the exclusion of the condensates from the central segment of the cells. In addition to the effect of the nucleoid, anisotropic confinement has also been hypothesised as contributing to pole accumulation of aggregates [8]. Pole accumulation consistently emerges from the distribution of MLOs positions that, however, also reveals a sparse third cluster in the central section of the cell (Extended Data Fig. 2). Image analysis confirms that most cells, 78% for A and 88% for B, contain two MLOs, while some feature three (20% for A and 8% for B, Fig. 1B(iii)-C(iii)). Rarely, cells have a single MLO, and a negligible fraction is condensate-free. Detailed statistics on condensate numbers are reported in Table 1. RT-qPCR was used to quantify NS expression levels, yielding an intra-cellular concentration of 16 ± 5 µM for NS B 1 hour post-induction (Supplementary Figs. 13, 14, Methods). By assuming that most NSs accumulate in the condensates, and approximating the per-NS volume in the condensed phase as that of a sphere with radius equal to the NS arm length (∼ 8 nm [69]), we can roughly estimate the total condensate volume from the NS concentration. By further assuming that this volume is equally divided between two condensates, we estimate a mean condensate diameter of ∼ 400 nm, compatible with microscopy observations (Fig. 1B(i)-C(i)). Intra-cellular NS concentration is comparable with in vitro expression yields, measured as 13 ± 4 µM 3 hours from starting the reaction (Supplementary Fig. 14).

**Table 1:**
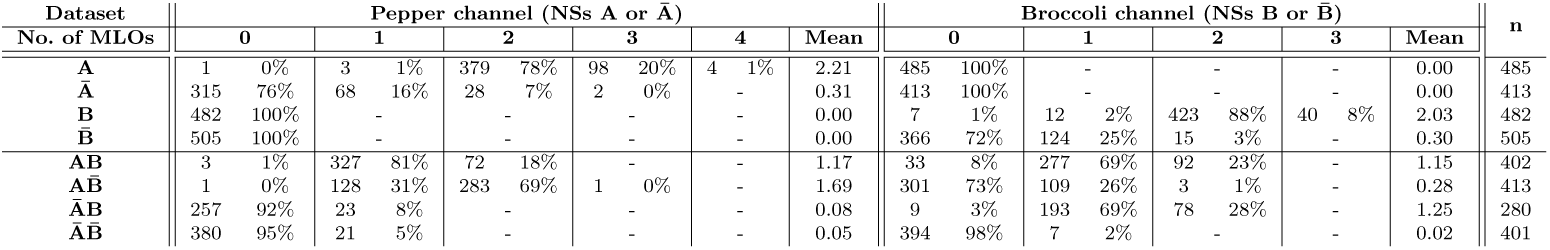
Statistical snapshot of the number and percentage of cells expressing a given number of MLOs of each type. . “Dataset” indicates experiment types with *E. coli* expressing one or a combination of NSs, as indicated. “Mean” indicates the mean number of MLOs per cell in each dataset. *n* is the number of cells analysed for each dataset. For each dataset, data are collated from 3 separate cultures, each starting from a single colony.

| **Dataset** | **Pepper channel (NSs A or Ā)** |  |  |  |  |  |  |  |  |  |  | **Broccoli channel (NSs B or B̄)** |  |  |  |  |  |  |  |  | **n** |
|---|---|---|---|---|---|---|---|---|---|---|---|---|---|---|---|---|---|---|---|---|---|
| **No. of MLOs** | **0** |  | **1** |  | **2** |  | **3** |  | **4** |  | **Mean** | **0** |  | **1** |  | **2** |  | **3** |  | **Mean** |  |
| **A** | 1 | 0% | 3 | 1% | 379 | 78% | 98 | 20% | 4 | 1% | 2.21 | 485 | 100% | - |  | - |  | - |  | 0.00 | 485 |
| **Ā** | 315 | 76% | 68 | 16% | 28 | 7% | 2 | 0% | - |  | 0.31 | 413 | 100% | - |  | - |  | - |  | 0.00 | 413 |
| **B** | 482 | 100% | - |  | - |  | - |  | - |  | 0.00 | 7 | 1% | 12 | 2% | 423 | 88% | 40 | 8% | 2.03 | 482 |
| **B̄** | 505 | 100% | - |  | - |  | - |  | - |  | 0.00 | 366 | 72% | 124 | 25% | 15 | 3% | - |  | 0.30 | 505 |
| **AB** | 3 | 1% | 327 | 81% | 72 | 18% | - |  | - |  | 1.17 | 33 | 8% | 277 | 69% | 92 | 23% | - |  | 1.15 | 402 |
| **AB̄** | 1 | 0% | 128 | 31% | 283 | 69% | 1 | 0% | - |  | 1.69 | 301 | 73% | 109 | 26% | 3 | 1% | - |  | 0.28 | 413 |
| **ĀB** | 257 | 92% | 23 | 8% | - |  | - |  | - |  | 0.08 | 9 | 3% | 193 | 69% | 78 | 28% | - |  | 1.25 | 280 |
| **ĀB̄** | 380 | 95% | 21 | 5% | - |  | - |  | - |  | 0.05 | 394 | 98% | 7 | 2% | - |  | - |  | 0.02 | 401 |

Through image segmentation applied to single cells, we determined the integrated fluorescence intensity of individual condensates, *I*_cond_, expected to be proportional to their volume. Similarly, we computed the average intensity in the condensate-free region, *i*_disp_, proportional to the concentration of the dispersed NS phase, and the average intensity across the entire cell, *i*_tot_, proportional to the overall NS concentration (Methods). In Supplementary Fig. 15, we show that *i*_disp_ is weakly correlated with *I*_cond_ (*R*^2^ = 0.21 and 0.24 for A and B, respectively), consistent with the notion that the NS behave similarly to an equilibrium phase-separating system, in which the concentration of the dispersed phase is independent on the volume of the dense phase. In addition, consistently with the lever rule, *i*_tot_ is strongly linearly correlated with *I*_cond_ (*R*^2^ = 0.70 and 0.83 for A and B, respectively).

When identical staining and imaging settings are applied, typical fluorescence intensity recorded from condensates expressed in vivo closely matches the one measured in vitro, suggesting similar FLAP response and NS concentration in the condensed phase (Supplementary Fig. 16). Inspection of the time evolution of individual cells reveals occasional instances in which small condensates form in the central section, before merging with one of the polar MLOs, suggesting a pathway for condensate accretion alternative to single-monomer addition (Fig. 1D(i) and Extended Data Fig. 3A), as previously observed for repeat-RNA MLOs [38]. We propose that the rapid migration of mid-cell nascent condensates toward the poles is driven by the energetic cost associated with nucleoid deformation (Supplementary Note 1). These coalescence events hint at a liquid-like state of the MLOs. Consistent with the liquid-like state, Fluorescence Recovery After Photobleaching (FRAP) experiments revealed partial recovery (Extended Data Fig. 3B-C). It should however be noted that recovery rates may be enhanced by the exchange of FLAP dyes [69].

Time lapses additionally suggest that a third MLO may appear near the equator of cells with two polar condensates as they approach division, when the central section is free from the nucleoid (Fig. 1D(ii) and Extended Data Fig. 4). Consistent with this picture, for both A and B designs, we find a significant correlation between the length of the cells and the number of hosted MLOs (Supplementary Fig. 17). In some instances, the central condensates then appear to split between the two daughter cells, which emerge with two polar condensates of asymmetric intensity (Fig. 1D(ii) and Extended Data Fig. 4(i)). Over time, smaller condensates grow towards parity with the larger ones, progressively reducing size asymmetry (Supplementary Fig. 18). This evidence excludes Ostwald ripening as a dominant mechanism for condensate coarsening, consistent with observations made on DNA nanostar condensates in vitro [84]. In other cases, the newly formed MLO partitions entirely in one of the daughter cells, leaving the other with a single condensate (Extended Data Fig. 4(ii)). Evidence of division confirms the viability of the MLO-expressing cells.

Expressing NS designs with scrambled KLs, Ā and B̄, largely results in uniformly distributed fluorescence (Figs. 1B(i)-(ii), 1C(i)-(ii) and Supplementary Figs. 19, 20), with a small fraction of the cells (23% for Ā and 28% for B̄, Fig. 1B(iii)-C(iii)) displaying regions of increased signal. These clusters likely arise from non-sequence-specific interaction between NSs or with other cellular RNA or proteins [85, 86]. Average fluorescence maps (Fig. 1B(ii)-C(ii)) and location analysis (Extended Data Fig. 2) show that, like specific MLOs, non-specific NS clusters are preferentially located at the cell poles.

Owing to imaging noise and cell-to-cell variation, it is sometimes challenging to distinguish slight non-uniformities in the fluorescent signal from NS condensates, which may lead to false-positive MLO detections for Ā and B̄ (Methods). To discriminate more reliably, we analyse the frequency distribution of pixel values within single cells and extract values for the kurtosis (Methods). A positive kurtosis reflects highly localised fluorescence, characteristic of MLO formation, whereas a negative kurtosis reflects a more uniform fluorescence distribution (Extended Data Fig. 5). Consistently, we observe positive kurtosis for NS A and B, and negative values for Ā and B̄ (Fig. 1E, and Supplementary Tables 4, 5). Alternative to the kurtosis, fluorescence-distribution skewness also discriminates well between MLO-forming and scrambled NS designs (Extended Data Figs. 5, 6).

Optical density measurements performed in bulk cultures reveal an expected decrease in growth rate upon induction, which is marginally more prominent in *E. coli* expressing MLO-forming NSs compared to those producing non-sticky designs (Extended Data Fig. 7A(i) and Supplementary Figs. 21, 22A). Simultaneous detection of NS fluorescence shows the anticipated signal growth upon induction, with stronger emission observed for A and B NSs compared to Ā and B̄ (Extended Data Fig. 7A(ii) and Supplementary Fig. 22B). Uninduced controls do not show significant induction leakage.

Given the limited half life of mRNA in *E. coli* [87], it is expected that nanostars may also be progressively degraded. We thus investigated stability and processing of the expressed A and B NSs with nanopore RNA sequencing and Polyacrylamide Gel Electrophoresis (PAGE) (Extended Data Figs. 8, 9). We find that both constructs are cleaved at specific sites, located on two of the arms of NS A and on one arm of NS B. For both nanostructures, fragments corresponding to all sections of the designed transcript are detected, implying that the observed fragmentation is likely the result of endonucleases acting on folded constructs, rather than abortive transcription. In this scenario, cleavage of one of the dsRNA arms would reduce NS valency without impacting the folded construct downstream of the restriction site. This picture is consistent with the evidence of robust MLO formation: while a fraction of NS population may have reduced valency, a sufficient proportion of the constructs must retain three or four KLs to support phase separation. Despite occurring at two distinct sites, enzymatic degradation is less pronounced for NS A compared to NS B (Extended Data Figs. 8B, 9B). Denaturing PAGE (Extended Data Figs. 8C, 9C) shows substantial quantities of the full, uncleaved transcript for NS A, which are not detected for NS B. Native PAGE shows the full-length construct as the dominant species for NS A, indicating that most cleavage events detected by sequencing may only nick the dsRNA arms, rather than fully detaching them, allowing the construct to retain the designed secondary structure (Extended Data Figs. 8D). For NS B, a moderate amount of the complete, folded construct is present (Extended Data Fig. 9D), likely featuring a nicked but not fully cleaved arm. The most abundant species, however, has lower molecular weight and likely corresponds to the NS lacking one KL and having, therefore, reduced valency (Extended Data Fig. 9D). This observation highlights the advantage of using tetravalent rather than trivalent NS designs: given that a minimum valency of three is required to sustain phase separation [69], the built-in redundancy of tetravalent constructs makes them intrinsically more resilient. In both NS designs, cleavage was not detected on NS arms that include the FLAP or the T7 terminator, possibly due to these structural features hindering RNase access [88]. Both NSs, however, feature a cleavage site in the terminator loop itself, which is thus likely to be detached in many transcripts (Extended Data Figs. 8A-B, 9A-B). The apparent protective effect of altering the duplex structure of NS arms could be leveraged to design against degradation. In addition, stability may be enhanced by circularising the constructs to prevent digestion by exonucleases [89].

### Expression of orthogonal membrane-less organelles in *E. coli*

KL orthogonality between A and B designs should enable co-expression of non-mixing MLOs. We test this hypothesis by designing plasmids containing two distinct NS sequences (Fig. 2A and Supplementary Fig. 10). Different T7 terminator sequences [81] are used for the two NS to minimise the potential for non-specific affinity between the constructs induced by the terminator loops.

**Fig. 2:**
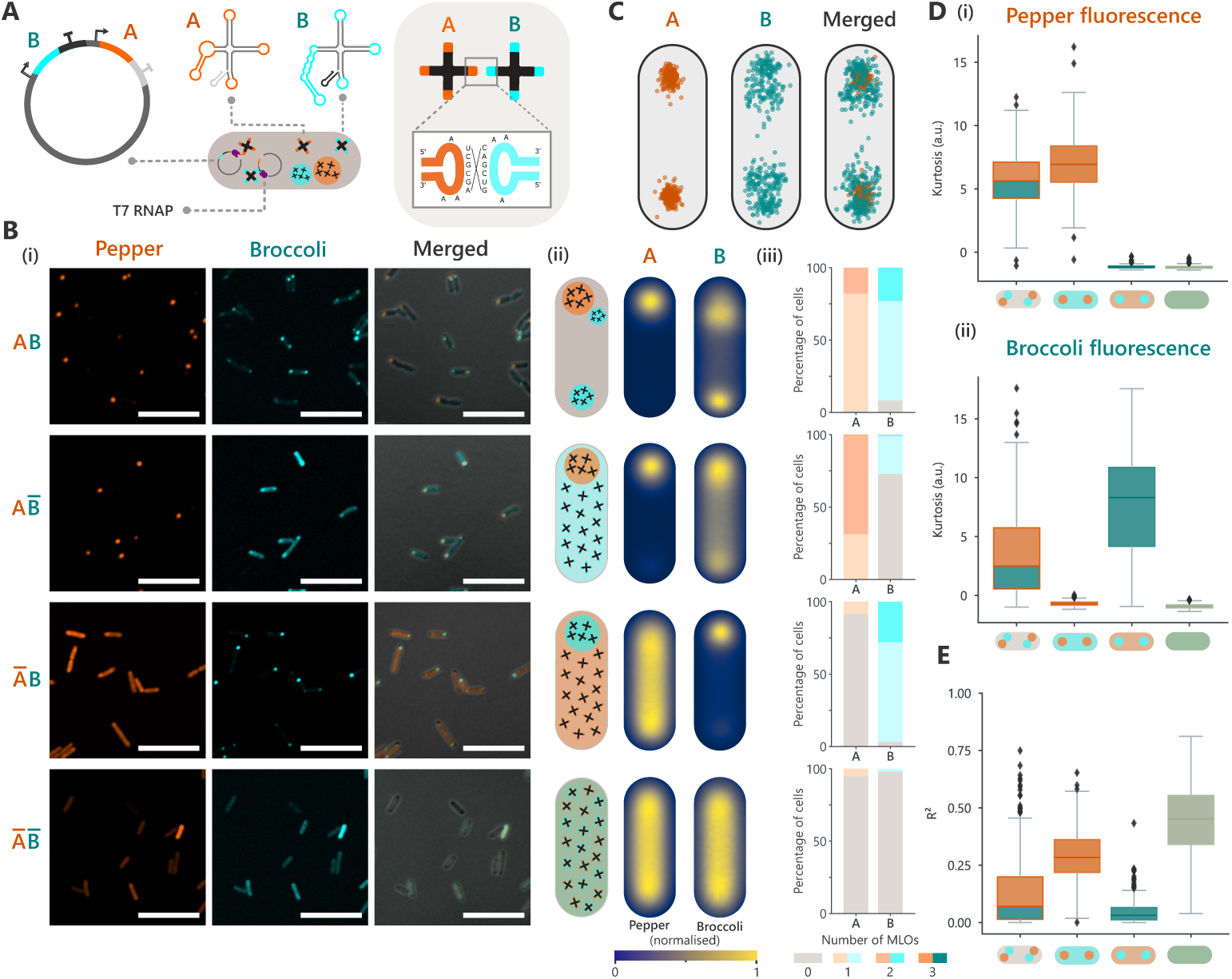
Co-expression of orthogonal, non-mixing RNA MLOs in *E. coli*. (**A**) Schematic of the co-expression of A and B NSs in *E. coli*. A and B motifs have different kissing loop sequences and cannot interact by base-pairing, therefore forming separate MLOs in *E. coli*. Both NS sequences are cloned into a single plasmid for T7 RNA polymerase transcription, each controlled by its own promoter and a distinct terminator [81]. (**B**) Co-expression of NS pairs in *E. coli*. Top to bottom: A+B, A+B̄, Ā +B and Ā +B̄ NSs. (i) Confocal micrographs. Left: Pepper fluorescence (NSs A and Ā). Middle: Broccoli fluorescence (NSs B and B̄). Right: Merged Pepper fluorescence, Broccoli fluorescence, and bright-field. Scale bars: 10µm. (ii) Schematic representation (left) and population-averaged maps of Pepper fluorescence distribution (middle), and Broccoli fluorescence distribution (right) in *E. coli* (Methods). (iii) Percentage of *E. coli* containing 0, 1, 2 or 3 A or B MLOs per cell. (**C**) Scatter plot of the locations of MLOs in *E. coli* co-expressing both A and B NSs. Each symbol represents the position of a detected MLO (Methods). (**D**) Box plots of the values of kurtosis extracted from the pixel-value distribution of fluorescence intensity in single cells. Analysis is performed on (i) Pepper and (ii) Broccoli fluorescence for cells expressing A+B, Ā +B, A+B̄ and Ā +B̄ NSs (left to right). Outcomes of statistical tests are provided in Supplementary Tables 6 and 7. (**E**) Box plots of the correlation coefficient (*R*^2^) values computed between Broccoli and Pepper fluorescent signals. Outcomes of statistical tests are provided in Supplementary Table 3. In **D**-**E**, each data point refers to an individual *E. coli*. In **B**-**E**, data are obtained from 3 separate cultures, each starting from a single colony.

When co-expressing A and B, confocal microscopy confirms the formation of non-mixing MLOs (Fig. 2B(i), Extended Data Fig. 10, and Supplementary Fig. 23). Similar to single-NS systems, condensates are preferentially located towards the poles, as evident from both average fluorescence maps (Fig. 2B(ii), Supplementary Fig. 24) and the locations of individual MLOs (Fig. 2C). Of the cells containing one A-type and B-type MLO (60% of all cells co-expressing NSs A and B), 66% had both MLOs positioned at the same pole. This asymmetric distribution may stem from the cells’ division history, as newly formed poles are less likely to harbour condensates. Alternatively, or in addition, it may be driven by weak, non-specific affinity between A and B condensates. We also observe a clear asymmetry between the exact positioning of A and B condensates: the former are nearly always in contact with a pole, while the latter may occupy more central locations (Extended Data Fig. 2). Inspection of individual micrographs confirms that B condensates tend to accumulate around polar A MLOs (Extended Data Fig. 10). We ascribe this behaviour to the tendency of A NSs to form larger condensates (Supplementary Fig. 25), which may appear faster than B MLOs and grow to occupy the entire polar region, pushing smaller B condensates more towards the central segment of the cell. Slower condensation kinetics for the B design may be linked to its greater susceptibility to degradation and consequent reduced valency.

Most cells express one MLO of each kind, with a smaller percentage containing two A and/or two B MLOs (Fig. 2B(iii) and Table 1). The overall number of MLOs in cells co-expressing A and B (averaging 2.3) is similar to that of cells expressing only A (2.2) or only B (2.0), indicating that co-expression does not significantly alter condensate formation.

Cells co-expressing one sticky NS and one non-sticky NS, A + B̄ or Ā + B, behave as expected, with MLOs of the intended NS species (A or B) coexisting with a largely diffuse phase of the soluble NSs (B̄ or Ā, Fig. 2B(i) and Supplementary Figs. 26, 27). The fluorescent signal distribution (Fig. 2B(ii)) and condensate locations (Extended Data Fig. 2) of A and B MLOs are similar to those observed for cells that do not simultaneously express B̄ or Ā (Fig. 1B(ii)-C(ii) and Extended Data Fig. 2), indicating that soluble NSs do not interfere with the condensation of sticky motifs. Condensate numbers are, however, lower, with a greater proportion of cells expressing a single A or B MLO (Fig. 2B(iii)), arguably due to a lower overall expression of the MLO-forming motif due to the simultaneous production of the soluble NSs.

The fluorescence distribution of B̄, when co-expressed with A, is largely uniform, with a small fraction of the cells (27%) displaying regions of increased B̄ signal at the poles (Fig. 2B(ii)-(iii) and Table 1), similar to that observed in cells expressing B̄ only (28%). The fluorescence distribution of Ā, when co-expressed with B, is almost completely uniform, with only 8% of cells displaying regions of increased Ā signal (Fig. 2B(ii) and Table 1), compared to 25% for *E. coli* expressing only Ā. Cells co-expressing the two soluble NSs, Ā and B̄, display near-uniform fluorescence distribution (Fig. 2B(ii) and Supplementary Fig. 28), with non-specific Ā and B̄ clusters detected only in 5% and 2% of the cells, respectively (Fig. 2B(iii) and Table 1).

The lesser tendency of Ā and B̄ to form non-specific clusters is consistent with a reduced NS concentration caused by the simultaneous expression of two constructs. The greater abundance of non-specific clusters seen when non-sticky NS are co-expressed with sticky motifs, particularly for A+B̄ expression, may be due to MLO-templated clustering of the otherwise soluble NSs. The soluble NSs may indeed weakly partition within, or coat the specifically-assembled MLO, or the condensates may serve as heterogeneous nucleation sites for non-specific aggregation. Either process may be facilitated by weak affinity between the soluble and sticky NSs due to base pairing, misfolding, enzymatic degradation, or the action of native RNA-binding proteins. This interpretation is supported by the evidence of weak partitioning of soluble NS B̄ in NS A condensates observed in vitro (Extended Data Fig. 1), and quantified in Supplementary Fig. 7 (Methods).

As done for single-NS systems, we use the kurtosis of pixel-intensity distributions to assess the tendency of individual NS components to condense. In line with observations made on Fig. 2B, for all combinations tested, cell populations expressing A or B NSs have positive kurtosis in the relevant fluorescent channels, while those expressing Ā or B̄ have negative kurtosis (Fig. 2D and Supplementary Tables 6, 7). Skewness analysis can similarly distinguish between cells expressing condensate-forming and non-sticky NSs (Extended Data Fig. 6).

To further assess the relative distribution of co-expressed NS components, we study the cross-correlation between Pepper (NS A) and Broccoli (NS B) fluorescence signals (Supplementary Fig. 29A), extracting distributions of the coefficient of determination, *R*^2^, across *E. coli* populations (Fig. 2E and Supplementary Table 3). In cells expressing A and B, the low *R*^2^ confirms that MLOs are orthogonal and non-overlapping. When expressing Ā and B̄, *R*^2^ is highest, as both NSs are evenly distributed throughout the cell. In *E. coli* producing A and B̄, a low *R*^2^ follows from the localised Pepper (NS A) signal in a diffused Broccoli (NS B) background. The higher *R*^2^ value found for cells expressing A+B̄ supports the hypothesis that B̄ may non-specifically partition within MLOs, consistent with observations made in vitro (Extended Data Fig. 1B, Supplementary Fig. 7).

Bulk growth curves collected for *E. coli* expressing two NS species are similar to those recorded for single-NS expression (Extended Data Fig. 7B(i) and Supplementary Fig. 30A). Samples show no significant differences in the trend and magnitude of the fluorescent traces recorded simultaneously (Extended Data Fig. 7B(ii) and Supplementary Fig. 30B). We also observe that some cells still contain MLOs after a week at room temperature, indicating long-term stability of the condensates despite enzymatic degradation of individual NSs (Supplementary Fig. 31).

### Selective protein recruitment in RNA membrane-less organelles

Owing to their modularity and controlled structure, the RNA motifs can be straightforwardly modified to feature additional functional motifs, including aptamers enabling the capture of molecular clients. As proof-of-concept, we embedded the GFP-binding AP3 aptamer [90] in one of the arms of NS A, opposite to the arm hosting the Pepper FLAP (Fig. 3A). Note that A_AP3_ remains tetravalent, as the kissing loop is inserted between two stem loops of the aptamer.

**Fig. 3:**
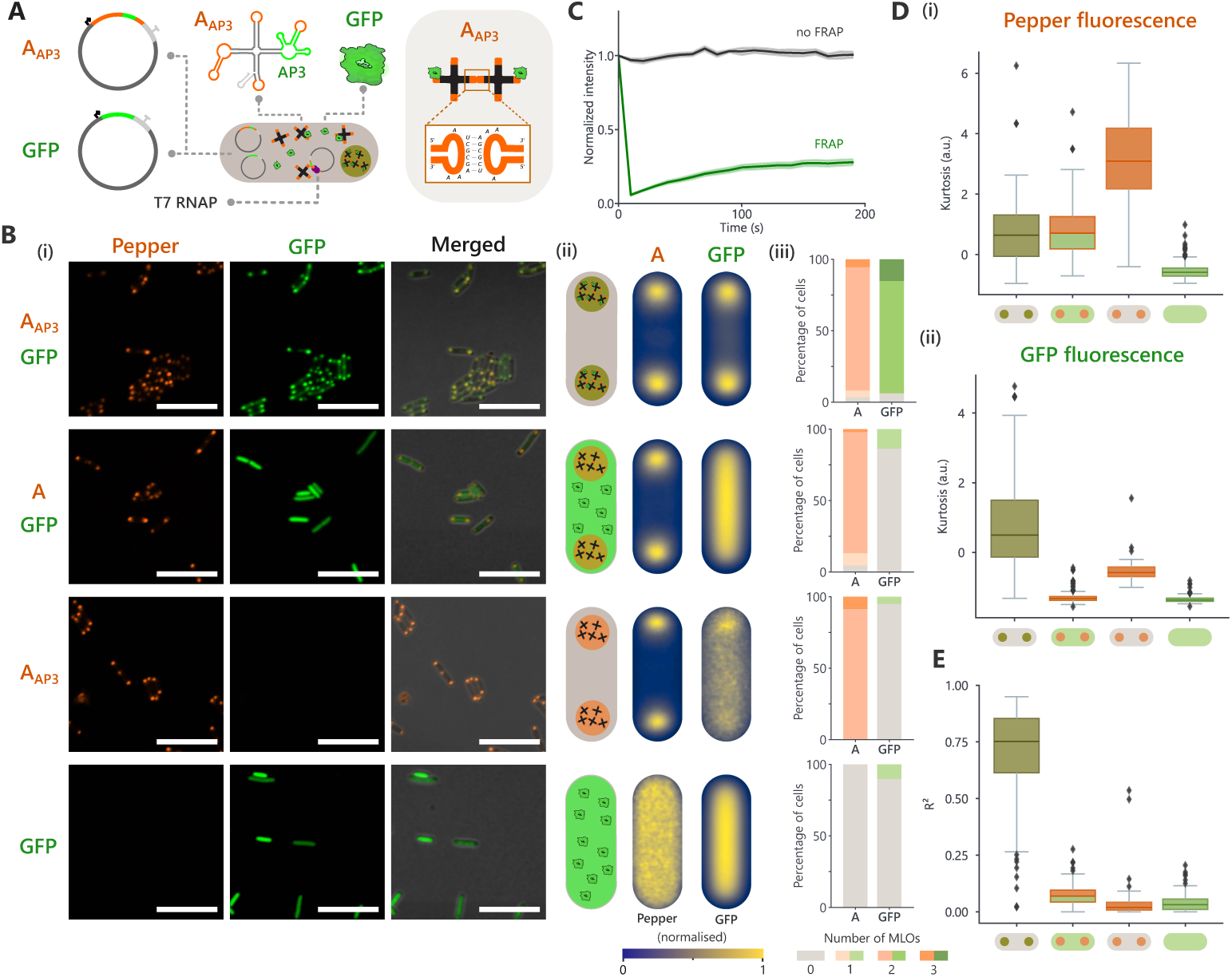
Proteins can be selectively captured in RNA MLOs expressed in *E. coli*. (**A**) Schematic of GFP co-expression with NS A_AP3_, carrying the GFP-binding AP3 aptamer [90], in *E. coli*. (**B**) Co-expression of NSs and GFP in *E. coli*. Top to bottom: A^AP3^ + GFP, A + GFP, A_AP3_ only, and GFP only. (i) Confocal micrographs. Left: Pepper (NS) fluorescence. Middle: GFP fluorescence. Right: Merged Pepper fluorescence, GFP fluorescence, and bright-field. Scale bars, 10 µm. (ii) Schematic representation (left) and population-averaged maps of Pepper fluorescence distribution (middle), and GFP fluorescence distribution (right). (iii) Percentage of *E. coli* containing 0, 1, 2 or 3 MLOs per cell. (**C**) Fluorescence Recovery After Photobleaching (FRAP) curves of GFP captured in A_AP3_ MLOs. For each MLO, fluorescence intensity prior to photobleaching is normalised to 1. The green solid line is the mean intensity of photobleached samples (*n* = 25 condensates), while the black solid line is the mean intensity of non-photobleached samples (*n* = 15 condensates). Shaded areas indicate the standard error of the mean. (**D**) Box plots of the values of kurtosis extracted from the pixel-value distribution of fluorescence intensity in single cells. Analysis is performed on (i) Pepper and (ii) GFP fluorescence for cells expressing A_AP3_ + GFP, A + GFP, A_AP3_ only, and GFP only. Outcomes of statistical tests are provided in Supplementary Tables 8, 9. (**E**) Box plots of the coefficient of determination (*R*^2^) values computed between Pepper and GFP fluorescent signals. Outcomes of statistical tests are provided in Supplementary Table 3. In **D**-**E**, each data point refers to an individual *E. coli*. In **B**-**E**, data are obtained from 3 separate cultures, each starting from a single colony.

Confocal micrographs and image analysis confirm that A_AP3_ expression in *E. coli* produces MLOs with similar spatial distribution and numbers as those measured for NS A (Fig. 3B(i)-(iii), Table 2, and Supplementary Fig. 32). Co-expression of GFP results in a clear co-localisation of the protein with the A_AP3_ signal (Fig. 4B(i)-(ii) and Supplementary Fig. 33). Cells expressing A_AP3_ alone and those co-expressing A_AP3_ and GFP both contained an average of 2.0 condensates per cell (Table 2), indicating that GFP expression does not noticeably affect A_AP3_ condensate formation.

**Fig. 4:**
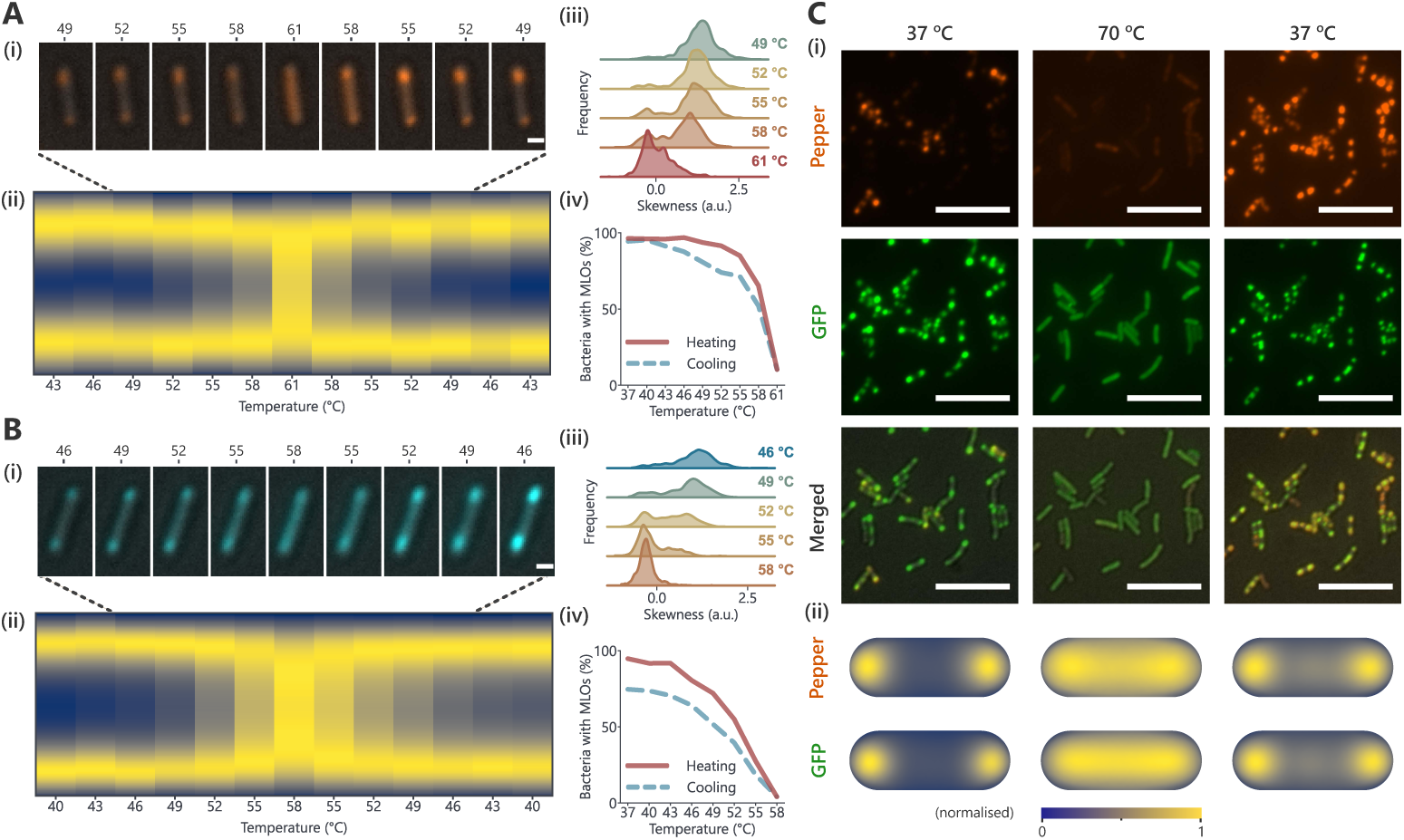
Synthetic RNA MLOs reversibly disassemble and reassemble in *E. coli* upon temperature cycling. (**A**) Reversible disassembly and reassembly of A MLOs. (i) Merged epifluorescence and bright-field micrographs showing disassembly and reassembly of A MLOs in a single cell. Scale bars, 1 µm. (ii) Population-averaged maps of Pepper (NS) fluorescence as a function of temperature upon heating and subsequent cooling. (iii) Cell-population distribution of the values of skewness computed for the fluorescence-intensity distributions of Pepper signal in individual *E. coli* on heating and cooling. (iv) Percentage of single cells containing MLOs as extracted from the skewness distributions (Methods). (**B**) Reversible disassembly and reassembly of B MLOs. Data are presented as in panel **A**, using Broccoli fluorescence to image NSs. (**C**) Disassembly and reassembly of A_AP3_ MLOs with captured GFP. (i) Confocal micrographs of samples at 37 *^◦^*C (left), after heating to 70 *^◦^*C (middle), and after cooling back to 37 *^◦^*C (right). Top: Pepper (NS) fluorescence; middle: GFP fluorescence; bottom: merged Pepper fluorescence, GFP fluorescence, and bright-field. Scale bars, 10 µm. (ii) Population-averaged maps of Pepper (top) and GFP (bottom) fluorescence at the temperatures shown in (i). In **A**–**C**, fluorescence maps and quantitative analyses are based on *E. coli* from 3 independent cultures, each starting from a single colony.

**Table 2:** Statistical snapshot of the number and percentage of cells expressing a given number of MLOs for protein capture experiments. . “Dataset” indicates experiment types with *E. coli* expressing NSs and/or GFP as indicated. “Mean” indicates the mean number of MLOs per cell in each dataset. *n* is the number of cells analysed for each dataset. For each dataset, data are collated from 3 separate cultures, each starting from a single colony.

| **Dataset** | **Pepper channel (NSs A or A<sub>AP3</sub>)** |  |  |  |  |  |  |  |  | **GFP channel** |  |  |  |  |  |  |  |  | **n** |
|---|---|---|---|---|---|---|---|---|---|---|---|---|---|---|---|---|---|---|---|
| **No. of MLOs** | **0** |  | **1** |  | **2** |  | **3** |  | **Mean** | **0** |  | **1** |  | **2** |  | **3** |  | **Mean** |  |
| **A<sub>AP3</sub>+GFP** | 5 | 3% | 7 | 5% | 125 | 86% | 8 | 6% | 1.94 | 9 | 6% | - |  | 114 | 79% | 22 | 15% | 2.03 | 145 |
| **A+GFP** | 9 | 4% | 18 | 9% | 174 | 85% | 4 | 2% | 1.84 | 177 | 86% | 28 | 14% | - |  | - |  | 0.14 | 205 |
| **A<sub>AP3</sub>** | - |  | - |  | 75 | 91% | 7 | 9% | 2.09 | 78 | 95% | 4 | 5% | - |  | - |  | 0.05 | 82 |
| **GFP** | 189 | 100% | - |  | - |  | - |  | 0.00 | 170 | 90% | 19 | 10% | - |  | - |  | 0.10 | 189 |

In contrast, GFP remains evenly distributed when co-expressed with NS A lacking the GFP-binding aptamer, confirming the specificity of the recruitment mechanism (Fig. 3B(i)-(ii) and Supplementary Fig. 34). Expressing GFP in the absence of NSs confirms the lack of spontaneous protein aggregation and the formation of inclusion bodies (Fig. 3B(i)-(ii) and Supplementary Fig. 35) [91]. The small number of GFP clusters detected by our segmentation algorithm in the absence of A_AP3_ NSs come from segmentation artefacts linked to the 3D shape of the cell [92], which results in brighter central sections that may be occasionally misclassified as MLOs.

Partial recovery of the GFP signal in FRAP experiments demonstrates that the recruited protein can exchange with cytosolic GFP (Fig. 3C).

Analysis of the kurtosis of the pixel-fluorescence distribution in Fig. 4D show the expected trends: positive kurtosis is measured in the Pepper (NS) channel for all MLO-forming NS designs, and in the GFP channel only when the protein-binding A_AP3_ is expressed (Supplementary Tables 8, 9). Consistent trends are observed when analysing skewness (Extended Data Fig. 6). Finally, image correlation analysis shows a strong fluorescence co-localisation between GFP and A_AP3_ MLOs but not between GFP and A MLOs (Fig. 3E, Supplementary Fig. 29, and Supplementary Table 3). Incidentally, the strong co-localisation between GFP and A_AP3_ confirms the limited impact of potential chromatic aberrations between the red fluorescence channel (used for Pepper-labelled NSs) and the green channel (used for GFP and Broccoli-labelled NSs.)

### Temperature-dependent dissolution and re-assembly of MLOs

Since the NSs self-assemble through base-pairing interactions, we anticipated that the MLOs would dissolve at sufficiently high temperature, as observed in vitro (Supplementary Figs. 36, 37, 38). Consistent with this expectation, heating *E. coli* containing A-MLOs led to their dissolution at 61 *^◦^*C. This process was reversible, with A-MLOs reassembling with similar polar distribution as the cells were cooled (Fig. 4A(i)-(ii) and Supplementary Fig. 39). Sampling the skewness of the pixel-intensity distribution across the bacteria population provides information of the condensate melting process (Fig. 4A(iii)). At the starting temperature, the skewness is high across the population, reflecting the presence of MLOs. As temperature is increased, the distribution becomes bimodal, with a second population of cells displaying low skewness following MLO dissolution. The intra-population variability in the onset of melting may be due to cell-to-cell variation in NS expression and therefore concentration. At sufficiently high temperature most cells display low skewness. By fitting the skewness distributions, we can extract a temperature-dependent fraction of cells hosting MLOs, which shows a small hysteresis between heating and cooling ramps. This may be caused by slow assembly kinetics, delays in thermal equilibration, or changes in protein crowding due to protein denaturation during heating, which could alter the local environment of the MLOs (Fig. 4A(iv) and Supplementary Fig. 40). B-MLOs showed a reversible disassembly behaviour similar to the A design, but with a lower melting temperature: condensates begin to melt at 49 *^◦^*C and fully dissolve by 58 *^◦^*C (Fig. 4B(i)-(iv) and Supplementary Figs. 40, 41). Single-cell skewness-temperature curves were used to estimate the melting temperature for each cell. Linear regression between the melting temperature and cellular fluorescence gave low coefficients of determination (*R*^2^ *<* 0.2), indicating no detectable correlation between NS concentration and melting temperature (Supplementary Fig. 42). The difference in melting temperature between the two constructs is unlikely to result from different KL affinity, having observed that NSs with KL A disassemble at slightly lower temperature (39.9 *^◦^*C) compared to those with KL B in vitro (40.9 *^◦^*C, Supplementary Figs. 36, 37, 38). Instead, it may result from the greater degree of enzymatic degradation experienced by NS B (Extended Data Figs. 8, 9). A reduction in NS valency is also consistent with the broader melting transition seen in NS B compared to the A design. Different from A NSs, a small fraction (∼ 20%) of cells expressing B MLOs do not reassemble condensates when cooled, possibly due to nanostructure misfolding or failure to refold upon cooling. For both A and B NSs, condensate fluorescence intensity appears to increase after the heating and cooling ramps (Fig. 4A(i), B(i)), which may be attributed to thermally induced alterations in membrane permeability, facilitating greater intracellular accumulation of FLAP dyes. We further note that melting temperatures measured in vivo are substantially higher than those we observe in vitro, where the MLOs melted at 39.9-40.9 *^◦^*C (Supplementary Fig. 36, 37, 38, 43). This deviation likely originates from intracellular molecular crowding promoting condensation [93] or difference in ionic conditions between the in vitro transcription buffer and the cytoplasm.

We then demonstrate temperature-triggered release of captured proteins by heating MLOs that had sequestered GFP. When cells expressing the A_AP3_ NSs and GFP were briefly heated to 70 *^◦^*C for 2 minutes, the GFP and nanostar fluorescent signal dispersed throughout the cell. Upon cooling, the MLOs reassembled at the poles, with GFP relocalising accordingly (Fig. 4C(i)-(ii) and Supplementary Figs. 44, 45). Approximately 10% of the cells retained the ability to grow and divide following thermal treatment (Supplementary Fig. 46). Growth and division is found to resume after a 3-hour delay, which is likely the time required for bacterial recovery [94].

## Conclusion

We demonstrate the expression of RNA nanostructures, capable of forming non-natural membrane-less organelles in live *E. coli*. Sequence selectivity enables the production of non-mixing and addressable organelle types with distinct composition, while embedding aptamers facilitates protein uptake. The genetically encoded, synthetic organelles are liquid-like and exhibit upper critical solution temperature behaviour, enabling reversible disassembly and reassembly upon heating and subsequent cooling. The system provides valuable insights on the biophysics of biomolecular condensation in prokaryotes, as we show by probing organelle number, spatial distribution, fluidity and response upon division. Compared to platforms reliant on engineered proteins and peptides, or simpler RNA repeats, the nano-structured motifs offer advantages in terms of rational and modular design. The observation that the RNA nanostars retain programmability and functionality within the complex intracellular environment is noteworthy, particularly for expression in *E. coli* that, despite its biotechnological relevance, poses substantial challenges due to rapid RNA degradation.

By overcoming the hurdle of RNA instability, our contribution paves the way to deploying nano-structured RNA condensates to metabolic engineering and biomanufacturing, leveraging the possibility of sequestering arbitrary proteins or nucleic acids within the condensates. One may indeed aim to optimise biocatalytic pathways by (co)localising enzymes and/or substrates, enhance expression yields by sequestering toxic products or intermediates, or improve separation and purification pipelines. Beyond industrial applications, the synthetic RNA condensates may be used to modulate biological functions in vivo by influencing gene regulation, buffering stress responses, and generally mimicking native phase-separated assemblies. For example, the condensates could be engineered to sequester transcriptional regulators or RNAs to alter gene expression profiles, protect key transcripts under stress, or serve as synthetic analogues of bacterial RNA granules [6]. These capabilities will add to the significant opportunities already offered by genetically-encoded RNA nanostructures, which have been deployed to regulate transcription [95], translation [96], and metabolic pathways [97, 98]. The platform could further be adapted to other specialised prokaryotic expression strains, including *Bacillus subtilis*, *Pseudomonas putida* and *Lactococcus lactis*. We expect that, taken together, these characteristics of robustness, modularity, structural programmability and interaction selectivity, will establish nano-structured RNA condensates as a prominent tool for next-generation cellular engineering.

## Supplementary information

Supplementary figures, supplementary tables, supplementary notes, and gene sequences are available in the supplementary information.

## Acknowledgements

L.D.M., L.M., and C.F. acknowledge support from the European Research Council (ERC) under the Horizon 2020 research and innovation programme (ERC-STG No 851667 - NANOCELL). L.D.M. acknowledges support from a Royal Society University Research Fellowship (UF160152, URF\R\221009). L.D.M., B.N. and M.T. acknowledge support from the UK Biotechnology and Biological Sciences Research Council (BBSRC) through a Japan Interdisciplinary Partnership Award Plus (BB/X012557/1) and from JST ASPIRE (JPMJAP24B4). M.T. acknowledges support from MEXT/JSPS KAKENHI (Nos. JP24H00070 and JP25H01361). R.R.S. acknowledges funding from the Biotechnology and Biological Sciences Research Council through a BBSRC Discovery Fellowship (BB/X010228/1). M.D. acknowledges support for a studentship awarded by Ernest Oppenheimer Fund (School of Physical Sciences, University of Cambridge). A.K. acknowledges the University of Cambridge Harding Distinguished Postgraduate Scholars Programme and the UK Engineering and Physical Sciences Research Council (EPSRC, 2928387). G.F. acknowledges funding from the Department of Chemistry at Imperial College London. S.P.N. acknowledges support from EPSRC (EP/S023518/1). The team thank Yuval Elani, Elisa Franco and Hirohide Saito for useful discussions and the support staff of the Department of Chemical Engineering and Biotechnology, University of Cambridge, especially Danita Pearson, for their dedicated assistance.

## Data availability

Raw sequencing data has been deposited to NCBI Bioproject database under BioProject accession number PRJNA1307508.

## Code availability

The data analysis code used in this publication is available at https://github.com/adamkni/Condensate Segmentation.

## Methods

### Chemicals and reagents

All reagents were obtained from Merck unless otherwise specified. *E. coli* Bl21(DE3) and XL10-Gold cells were obtained from Agilent. DH5*α* cells were purchased from Thermo Fisher Scientific. TBI was purchased from Torcis Bioscience. HBC620 was purchased from MedChem Express. DFHBI-1T was purchased from Cayman Chemical. Recombinant GFP was purchased from Caltag Medsystems. Pico-Surf fluorinated oil was purchased from Sphere Fluidics. CellBrite Fix 640 membrane dye was purchased from biotium. The T7-FlashScribe™ Transcription Kit was purchased from Cellscript. The RNeasy Mini kit and Bacteria Protect Reagent were obtained from Qiagen. GeneRuler 100 bp DNA Ladder and RiboRuler Low Range RNA Ladder were obtained from Thermo Fisher Scientific. LB-agar plates, glycerol, ultra-pure water, and SOC media were obtained from the technical support team of the Department of Chemical Engineering and Biotechnology, University of Cambridge. Linear DNA strands were obtained from Integrated DNA Technologies (IDT) as gBlocks. Gene-frames were obtained from Thermo Fisher Scientific. Glass capillaries were purchased from VitroCom. Nunc MicroWell 96-Well Optical-Bottom Plates with Polymer Base (Cell culture, black, Surface Treatment: Cell Culture, Bottom: Flat) were obtained from Thermo Fisher Scientific. The pET plasmid (53137) was obtained from Addgene. The NEBuilder HiFi kit and the Monarch miniprep kit were obtained from New England Biolabs (NEB).

### Expression and characterisation of RNA nanostars in vitro

In vitro transcription reactions were prepared at room temperature using the T7-FlashScribe™ Transcription Kit and 40 nM of the relevant linear DNA template with T7 terminator or 5 nM of the relevant plasmid (for samples used in PAGE analysis). The final reaction mixture contained 9 mM of each ribonucleotide triphosphate, 0.05 units per µL RNase inhibitor, and 10 mM dithiothreitol. In addition, the reaction contained 50 µM of the Broccoli aptamer-binding dye DFHBI-1T, and 0.5 µM of the Pepper aptamer-binding dye HBC620. Mixtures were loaded into borosilicate glass capillaries with rectangular cross section (0.2 mm × 2 mm, VitroCom) and capped with paraffin oil. Capillaries were then immediately placed at 37 *^◦^*C and incubated for 4 hours, until imaging. Epifluorescence micrographs were obtained using a Nikon Eclipse Ti2-E inverted microscope equipped with a Hamamatsu ORCA-Flash4.0 V3 sCMOS camera, a tuneable light source (Lumencor SPECTRA X LED engine), and a Plan Apo *λ* 20 × 0.75 N.A. dry objective. The HBC620 dye was excited using a 590 nm LED source with an LED-DA/FI/TR/Cy5-B filter, 10 ms excitation time, and fluorescence was recorded through an FF01-624/40 emission filter. DFHBI-1T was excited using a 470 nm LED source with an LED-DA/FI/TR/Cy5-B filter, 500 ms excitation time, and fluorescence was recorded through an EM510 emission filter. To determine the melting temperature of NS A and NS B, samples were incubated for 15 h at 30 *^◦^*C and subsequently heated using a Peltier-controlled microscope stage (Temikra Ltd.). The temperature was increased from 30 *^◦^*C to 50 *^◦^*C and then cooled back to 30 *^◦^*C.

During this process, the temperature was adjusted by 1 *^◦^*C every 5 min, with imaging performed after each temperature change. For directly comparing the fluorescence intensity of in vitro condensates with that recorded *E. coli* (Supplementary Fig. 16), the in vitro transcription reaction was performed as previously described, except for replacing DFHBI-1T with 10 µM TBI, as used for *E. coli*, followed by a 15 h incubation at 30 *^◦^*C. For the in vitro transcription of NS A_AP3_, the reaction was prepared as described above with the addition of 750 nM recombinant GFP. The transcription mixture was then encapsulated within water-in-oil droplets by mixing 22 µL of the solution with 90 µL of 2% w/w Pico-surf fluorinated oil and vortexing at 2,500 rpm for 30 s. The top layer, containing the emulsion droplets, was extracted, loaded into capillaries, and incubated at 37 *^◦^*C for 8 h before being imaged on the Nikon Eclipse Ti2-E inverted microscope. The settings described above were used for imaging the Pepper channel (NS A_AP3_), while GFP fluorescence was imaged using the 470 nm LED source with an LED-DA/FI/TR/Cy5-B filter and 2 ms excitation time. To induce thermal disassembly of the in vitro A_AP3_+GFP samples, the loaded capillaries were heated using a Peltier microscope stage (Temikra Ltd.) to 20 *^◦^*C, 37 *^◦^*C and 40 *^◦^*C, and subsequently cooled to 20 *^◦^*C. To test the behaviour of NSs in buffer mimicking ionic conditions in *E. coli* (Supplementary Fig. 8), transcribed NSs were purified by Monarch Spin RNA Cleanup Kit (NEB), then 500 nM NS was added to 15 µL of buffer (40 mM Tris pH 8.0, 200 mM KCl, 1 mM MgCl_2_). The solution was incubated at 37 *^◦^*C in 384-Well Optical-Bottom Microplate, black, non-treated PS surface (ThermoFisher) for 1 h before epifluorescence imaging.

### Image analysis of NS condensates in vitro

Images (16-bit) were extracted from .nd2 (Nikon) or .czi (Zeiss) files. Fluorescence intensities reported in Supplementary Fig. 7 and for the comparison of in vitro and in vivo fluorescence intensities were quantified as the mean pixel intensity on mask-segmented images. Segmentation masks were generated using Otsu thresholding on flattened images, which were then applied to the corresponding raw images for intensity measurements. For melting-temperature analysis (Supplementary Fig. 36), raw 16-bit images were used to calculate the ratio between the standard deviation of the pixel values and the mean pixel intensity (*σ*/I). The melting and condensate-formation temperatures were determined as the points of intersection between the ratio curves and the plateau, defined as the mean of data points from 41 *^◦^*C onwards plus a constant of 0.05. To generate the displayed micrographs, images were rescaled to facilitate visualisation by dividing the pixel intensities by the intensity of the brightest pixels (99.9-100th percentile), and then exported as coloured micrographs. For micrographs showing condensate melting, condensate formation, and TBI dye comparisons, the brightest pixels across the entire image set were used for division. For all other micrographs, the brightest pixels within each individual image was used for division. The Python packages tifffile and numpy were used for handling and processing the microscopy images, scikit-image was used for image masking and thresholding, Pillow was used for micrograph export, and Matplotlib was used for creating plots.

### Plasmid construction

#### Nanostar design

RNA NSs A, Ā, B, and B̄ were adapted from Ref. [69]. Each construct consists of a four-way junction with 25 bp double-stranded RNA arms. One arm in each design contains a FLAP, Pepper for A and Ā [74], and Broccoli for B and B̄ [76]. Each arm in NS A ends with a KL with sequence 5’-AUCGCCAAA-3’, while arms in NS B terminate with KL 5’-AGUCGACAA-3’. The KL sequences were designed in Ref. [69] to ensure similar binding strength while retaining mutual orthogonality and self-complementarity. NSs Ā and B̄ have KL nucleotide sequences scrambled to eliminate self-complementarity. NS A_AP3_ additionally features the GFP-binding AP3 aptamer in one of the arms, opposite to the one hosting the Pepper aptamer [90]. NS arms that include fluorescent or GFP-binding aptamers retain KLs, ensuring that the constructs remain tetravalent. All NS designs were verified with Nupack [99] and Kinefold [100]. Sequences of all NS designs are reported in Supplementary Table 1.

#### Plasmid design and construction

To construct plasmids expressing single NSs (A, Ā, B, B̄, A_AP3_), the natural T7 terminator sequence [81] was appended to the 3’ end of the NS sequences. The NS+terminator sequences were cloned into a pET plasmid with ampicillin resistance gene (Addgene #53137), using NEBuilder HiFi kit, following the manufacturer’s instructions. To construct plasmids expressing two NSs (A+B, A+B̄, Ā +B, or Ā +B̄), the natural T7 terminator was appended to the 3’ end of NSs A and Ā, while a T7 terminator variant [81] was used for NSs B, and B̄. Different terminators were used to reduce chances of non-specific interactions between the two constructs induced by the terminators. As for single NSs, the resulting sequences were cloned into the pET plasmid (Addgene #53137) using the NEBuilder HiFi kit. A 44 bp linker separates the two NS sequences on the plasmid. All plasmid maps are provided in Supplementary Figs. 9, 10. The GFP expressing plasmid was constructed by cloning the eGFP sequence from pEGFP-N1 (Takara) into the pACYCDuet-1 backbone (Novagen), from which LacI had been removed. Both DNA fragments were obtained as linear sequences and assembled using the NEBuilder HiFi kit. The plasmids containing ampicillin resistance genes were propagated with XL-10 Gold cells, while plasmids containing chloramphenicol resistance genes were propagated in DH5*α* cells. Plasmids were extracted from the cloning strains using the Monarch miniprep kit. The sequences of all of the constructed plasmids were verified by whole-plasmid sequencing (Full Circle Labs).

#### RNA nanostars expression in *E. coli*

Throughout all experiments, antibiotics matching the relevant resistance genes were used, namely ampicillin (50 µg mL*^−^*^1^), and chloramphenicol (34 µg mL*^−^*^1^).

#### Transformation of BL21(DE3) strains

To transform BL21(DE3) bacteria with plasmids containing the ampicillin resistance gene, 10 ng of the target plasmid were mixed with 10 µL of BL21(DE3) competent cells, and incubated on ice for 15 min. The samples were plated on LB-agar containing carbenicillin and incubated at 37 *^◦^*C for 16 h. To transform BL21(DE3) bacteria with plasmids containing chloramphenicol resistance gene, or for co-transformation with plasmids containing different antibiotic resistance genes, 10 ng of each plasmid were added to 10 µL of BL21(DE3) competent cells and incubated on ice for 30 min. The samples were placed in a 42 *^◦^*C water bath for 30 s, followed by 2 min incubation on ice, and addition of 250 µL Super Optimal broth with Catabolite repression (SOC) medium. The samples were placed in a shaking incubator (Multitron Standard, Infors HT, 37 *^◦^*C, 225 rpm) for 1 h, before plating on LB-agar with matching antibiotics, and incubating at 37 *^◦^*C for 16 h. LB-agar plates with transformed cells were stored at 4 *^◦^*C for up to two weeks before RNA expression experiments.

#### RNA and protein expression in *E. coli*

A single colony was transferred to 5 mL M9 minimal medium, supplemented with 0.4% glycerol, 0.2% casamino acids, 2 mM MgCl_2_, 100 µM CaCl_2_ (hereby referred to as M9 medium with supplements), and matching antibiotics. The culture was grown for 4 h (225 rpm, 37 *^◦^*C) in a shaking incubator, to reach an OD_600_ around 0.6-0.8, measured by spectrophotometer (SmartSpec Plus, Bio-Rad). IPTG (0.4 mM), TBI (10 µM), HBC620 (5 µM) were added to 2 mL of the sample, and the samples were further incubated for 1 h (225 rpm, 37 *^◦^*C) in a shaking incubator. TBI dye was used instead of DFHBI-1T for imaging in cells due to its improved photostability in vivo, which makes it better suited for cellular imaging applications [76]. 10 µL of the sample were diluted with 10 µL of the M9 medium with supplements, including the same concentrations of dyes, inducer, and antibiotics. The samples were stained with 0.2 µL of CellBrite Fix 640 membrane dye (1000 ×) at 25 *^◦^*C for 10 minutes, before transferring to agarose pads for imaging.

#### Extraction of RNA from *E. coli*

Bacteria samples were prepared as described previously. RNA was extracted from bacteria using Qiagen RNeasy Mini kit with Bacteria Protect Reagent, following the manufacturer’s instructions. In the cell lysis step, the cells were sonicated for three cycles at maximum speed for 30 s, with 1 min rest between the cycles (Soniprep 150, MSE supplies LLC).

#### Quantifying NS expression in *E. coli* with RT-qPCR

Extracted RNA samples were reverse-transcribed and amplified using the Takyon No ROX SYBR (Eurogentec) mastermix and Takyon One Step converter (Eurogentec) in MicroAmp Optical 96-well reaction plates with optical adhesive covers (Applied Biosystems) on the QuantStudio 6 Pro qPCR (ThermoFisher). 60 ng of total RNA per sample was loaded into each well in a total reaction volume of 7.1 µL. Three biological replicates and three technical replicates were used for each sample. The cycle number at which a sample’s fluorescence signal crosses a predefined threshold (Cq) was recorded for each sample. To quantify the relative amount of target transcripts, a housekeeping gene (*adk*) was used as a reference. The relative transcript number (FD) is then calculated by:

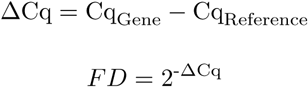

Primers B fwd and B rev targeted NS B, while primers adk fwd and adk rev targeted *adk*. Primers where selected to amplify the first half of the NS, to facilitate the detection of cleaved constructs (Extended Data Fig. 9). The primer sequences were: B fwd: 5’-GCACAGTGCTATGAGTGTC-3’, B rev: 5’-CGACTGTGGCATACAGCGAC-3’, adk fwd: 5’-ATTCTGCTTGGCGCTCCGGG-3’, adk rev: 5’-CCGTCAACTTTCGCGTATTT-3’. The thermocycling parameters were as follows: 48 *^◦^*C (10 min), followed by 95 *^◦^*C (3 min), then 40 cycles of 95 *^◦^*C (10 s) and 60 *^◦^*C (1 min). The temperature ramp rate was 1.6 *^◦^*C s*^−^*^1^ for all steps. To construct the calibration curve, in vitro transcribed NS RNA was purified using the Monarch Spin RNA Cleanup Kit (NEB). The concentration of purified NS was quantified by UV absorbance using a Nanodrop One spectrophotometer. Calibration curves was created by measuring the Cq values at different NS concentrations. The intracellular concentration of NS was estimated by dividing the amount of extracted RNA by the overall intracellular volume. The overall intracellular volume A calibration curve created using known purified RNA concentrations and was used calculate the concentration of RNA per mL of culture or per cell. To compute the latter, we estimated the total number of cells in the culture using optical density (OD) (1 OD = (1.0 ± 0.3) × 10^8^ colony forming units mL*^−^*^1^, based on 4 independent measurements), and multiplied by the expected single-cell volume of 3 *µ*m^3^, obtained by approximating the cell as a cuboid with dimensions 1 µm × 1 µm × 3 µm.

#### Preparation of agarose pads for imaging

For *E. coli* imaging on agarose pads, we prepared 2% molten low gelling temperature agarose in M9 media with supplements and added IPTG (0.4 mM), TBI (10 µM), HBC620 (5 µM), and matching antibiotics. The molten agarose solution was kept at 50 *^◦^*C, and 100 µL were added to a smaller gene frame (10 mm × 10 mm) adhered to a microscope slide (Clear ground 0.8-1.0 mm, Fisher Scientific). The molten agarose was sandwiched between two glass slides and left at 25 *^◦^*C for 10 min to solidify. The solidified agarose was transferred to larger gene frames (17 mm × 28 mm) adhered to a glass slide. 2 µL of the prepared cells were added to each agarose pad, and after 5 minutes at 25 *^◦^*C, a coverslip (24 mm × 60 mm, No.1, Epredia) was adhered to the other side of the gene frame.

#### Confocal imaging of *E. coli*

The prepared agarose pads were imaged with a Zeiss LSM 800 confocal microscope, using a Plan-Apochromat 63× 1.4 N.A. Oil DIC M27 objective. The fluorescent signal from the Pepper aptamer was acquired using a 561 nm laser set to 1.0% intensity and a Master Gain of 500 V, with an emission window of 580 to 650 nm. The fluorescent signal from the Broccoli aptamer and GFP was acquired using a 488 nm laser set at 2.0% intensity for A+B̄ and 1.0% for all other samples and a Master Gain of 500 V, with an emission window of 470 to 580 nm for all samples other than for A+B̄, which had a restricted emission window of 470 to 530 nm. The settings for samples A+B̄ were optimised to minimise bleed-through of the Pepper fluorescence in the Broccoli channel. While this fluorescence is weak and generally not noticeable in other samples, it becomes noticeable here because A is concentrated at MLOs, and B̄ is mostly evenly distributed (Supplementary Fig. 47). The image frame size was set to 1192 pixels × 1192 pixels with 16 bit depth. Line-averaging was enabled and set to 4×. The scan mode along the *x* direction was selected to be bidirectional. The scanning rate was set to produce a scanning time of 1.77 µs per pixel. This protocol was also used for imaging the in vitro samples when comparing fluorescence intensities between the in vitro and *E. coli* samples.

#### FRAP of *E. coli*

The 561 nm laser was set at 100% intensity for photobleaching A MLOs, while the 488 nm laser was set at 100% intensity for photobleaching B MLOs and GFP-containing MLOs. A total of 25 A MLOs, 12 B MLOs, and 25 GFP-containing MLOs were chosen for photobleaching, with the entire condensate bleached in each case. A total of 15 A MLOs, 8 B MLOs, 15 GFP-containing MLOs were chosen as non-photobleached controls. Fluorescence recovery was monitored at 10 s intervals. Condensate intensity was quantified as the mean intensity of the 10 brightest pixels within the MLO.

#### Epifluorescence imaging of *E. coli*

##### Time-lapse images of *E. coli*

Time-lapse images of cells expressing NS A were acquired with a Ti2 Eclipse Epi-fluorescent microscope (Nikon), equipped with an sCMOS camera (Hamamatsu ORCA-Flash4.0 V3), a tunable light source (Lumencor SPECTRA X LED engine), and CFI Plan Apo *λ*D 60× 1.42 N.A. oil immersion objective. The Pepper aptamer was excited using a 555 nm LED with an LED-DA/FI/TR/Cy5-B filter, and fluorescence was collected through an FF01-624/40 emission filter with a 500 ms exposure time. Images were taken every 5 minutes. Samples were incubated at 37 *^◦^*C using a heating stage (Temikra Ltd.).

##### Temperature-dependent dissolution and re-assembly of MLOs

To study the melting and reforming of MLOs in bacteria, bacteria with NS A or NS B were prepared for imaging as described previously, except that 2% agarose low EEO was used for the agarose pads. *E. coli* samples on agarose pads were attached to a Peltier microscope stage (Temikra Ltd.) equipped with sapphire microscope window to ensure optical transparency and good thermal conduction (UQGOptics), and imaged with a Ti2 Eclipse Epifluorescent microscope (Nikon), equipped with an sCMOS camera (Hamamatsu ORCA-Flash4.0 V3), a tuneable light source (Lumencor SPECTRA X LED engine), and a CFI Plan Apo *λ*D 60× 1.42 N.A. oil immersion objective. The Pepper aptamer was excited using a 555 nm LED with an LED-DA/FI/TR/Cy5-B filter, and fluorescence was collected through an FF01-624/40 emission filter with a 500 ms exposure time. The Broccoli aptamer was excited using a 470 nm LED with an LED-DA/FI/TR/Cy5-B filter, and fluorescence was collected through an EM510 emission filter with a 2 s exposure time. The samples were heated from 37 *^◦^*C to either 58 *^◦^*C or 60 *^◦^*C with increments of 3 *^◦^*C every 2 min. Samples were then cooled from their maximum temperature to 37 *^◦^*C with increments of 3 *^◦^*C every 2 min. For bacteria co-expressed with NS A_AP3_ and GFP, cells were imaged using the same Pepper aptamer settings as for Nanostar A, while the GFP signal was captured on the 470 nm LED with an LED-DA/FI/TR/Cy5-B filter, and fluorescence was collected through an EM510 emission filter with a 50 ms exposure time. Cells were heated to 37 *^◦^*C, 60 *^◦^*C, 70 *^◦^*C and cooled to 37 *^◦^*C, with 2 min incubation periods between each temperature.

##### Analysis of *E. coli* micrographs

Images and metadata were extracted from .czi (Zeiss) or .nd2 (Nikon) files using pylibCZIrw and Nd2Reader, respectively. The brightfield image underwent flat-field correction and non-local means denoising. Initial binary masks of the cells were generated using a calculated threshold and adjusted manually if needed. Small holes were filled, and objects with an area smaller than half of a typical *E. coli* cell (2 µm × 0.5 µm) were removed. Since the algorithm struggled to segment individual bacteria within microcolonies, bacteria inside these regions were filtered out based on object proximity. Objects on image borders were discarded. The remaining objects were filtered based on eccentricity (higher than 0.85), aspect ratio (between 2.5 and 4.5), area (smaller than 7.07 µm^2^), and width of *E. coli* (smaller than 1.5 µm) [101–104]. The final segmented set was further filtered manually via an interactive GUI. For *R*^2^ calculation, pixel intensities from one fluorescent channel were compared against those from another, yielding the coefficient of determination (*R*^2^). To compute kurtosis and skewness for each bacterium, pixel intensities in segmented single cells were normalised so that the top 5% of pixels had a value of 1. These pixels were then removed, and kurtosis and skewness of the pixel-intensity distribution were calculated using Scipy [105]. To determine the number of MLOs per bacterium, each segmented bacterium was rotated so that its long axis aligned horizontally. To minimise interpolation loss, pixels were subdivided into 16 smaller pixels before rotation. Intensities were then averaged vertically to obtain a linear profile, from which peaks were detected using Scipy. The sensitivity of peak detection was manually adjusted based on noise levels. For heatmaps of fluorescence intensity, horizontally aligned bacteria were scaled to a standardised size of 48 pixels × 130 pixels, and pixel intensities were averaged across aligned positions. For Fig. 2B(ii), cells were aligned with their most fluorescent region oriented upwards. This was determined by the Pepper channel for cells co-expressing nanostar A and B/B̄, and by the Broccoli channel for cells co-expressing nanostar Āand B. To determine locations of MLOs, the condensates were segmented using adaptive thresholding (cv2) [106]. MLO positions and sizes were validated by comparing them to peaks detected at similar locations within the cell to prevent misclassification. The MLO position was recorded as the highest-intensity pixel within the mask. For quantification of condensate and dispersed-phase intensities, local intensity maxima were identified in each segmented cell. Only cells in which the detected number of peaks matched the expected condensate count were retained. Around each peak, a 2D Gaussian was fitted and an elliptical condensate mask was generated, which defined the condensate region. The integrated condensate intensity, *I*_cond_, was computed as the sum of pixel intensities within this region. The dispersed-phase intensity, *i*_disp_, was calculated as the mean intensity in the remaining cell area after excluding the condensate part. *i*_tot_ was computed as the mean intensity within the entire cell area. All analyses were performed using Python 3.10, and visualisations were generated with Matplotlib [107] and Seaborn [108]. To generate the displayed fluorescence micrographs, images were rescaled to facilitate visualisation by dividing the pixel intensities by the intensity of the brightest pixels (99.9-100th percentile). To generate the displayed bright-field micrographs, intensities were rescaled by subtracting the intensity of the dimmest pixel, and dividing by the range between the brightest and dimmest pixels. For microgrpahs showing condensate melting and condensate formation, the brightest pixels across the entire image set were used for division. For all other micrographs, the brightest pixels within each individual image was used for division.

#### Analysis of the temperature-dependent dissolution and reassembly of MLOs

Micrographs were analyzed fully automatically to obtain segmented bacteria. Heatmap progression across different temperatures was determined by extracting the maximum intensity along the long axis of standard heatmaps, computed as discussed above. The skewness of the intensity distribution of the pixel intensity within single cells was used to monitor the progression of dissolution and reassembly. To estimate the proportion of cells with and without MLOs, the distribution of skewness values across the cell population at a given temperature was fitted using a mixture of two Gaussians (using skimage), with one Gaussian fitting the sub-population of cells with high skewness (featuring condensates) and the second fitting the sub-population of cells with low skewness (lacking condensates) [109]. For each cell, the probability of belonging to either sub-population was calculated. If the probability of belonging to the MLO-forming distribution exceeded 0.5, the cell was classified as MLO-forming. To estimate *T*_m_ for single cells, cells with skewness above 0.25 at 37 *^◦^*C (indicating the presence of condensates) were selected, and their skewness-temperature profiles were fitted with a sigmoid curve. The inflection point of the sigmoid fit was taken as the estimated *T*_m_.

### Growth and RNA production curves in bulk cultures

To characterise growth and RNA production in bulk cell samples, *E. coli* from single colonies were grown in M9 media with supplements for 4 h as described previously. *E. coli* samples were then diluted to OD_600_= 0.05. 100 µL of the diluted solutions were loaded in a Nunc MicroWell 96-Well Optical-Bottom well plate, together with TBI (10 µM), HBC620 (5 µM), and matching antibiotics. The samples were cultured at 37 *^◦^*C in a plate reader (Tecan M2000). Every 7 minutes, the plate was shaken (orbital) for 5 s with 3.5 mm amplitude, followed by absorbance and fluorescence measurements. For absorbance, wavelength was set to 600 nm and bandwidth was set to 9 nm. For detecting Broccoli fluorescence we set excitation wavelength to 470 nm, excitation bandwidth to 9 nm, emission wavelength to 520 nm and emission bandwidth to 20 nm. For Pepper fluorescence, we set excitation wavelength to 570 nm, excitation bandwidth to 9 nm, emission wavelength to 620 nm and emission bandwidth to 20 nm. When the OD_600_ reached 0.7, we added 0.4 mM IPTG to the samples requiring induction, before continuing the experiment with unchanged settings.

### RNA sequencing of *E. coli*

*E. coli* expressing A or B nanostars were induced for 1 h as described previously. Cells were pelleted and shipped to Full Circle Labs for RNA extraction, poly(A) tailing, and nanopore sequencing. To identify start and end positions of RNA raw reads [110], the reads were aligned to the expected RNA sequence with Needleman-Wunsch algorithm [111], and reads with alignment scores corresponding to ≥ 7 matched nucleotides were retained for further analysis. Due to known imprecision in basecalling near the 5’ ends of direct nanopore RNA reads [112], fragment start positions were defined as the first aligned nucleotide in a stretch of at least 3 consecutive matches between the read and the expected sequence.

### Polyacrylamide gel electrophoresis

The gels were prepared with a 19:1 w/v acrylamide:bis-acrylamide solution in Tris-Borate-EDTA (TBE) buffer. Acrylamide fractions (w/w) of 6% and 8% were used for native and denaturing gels, respectively. The denaturing gel contained 7 M urea. For both native and denaturing PAGE, samples transcribed in vitro were diluted 1:10, while for *E. coli* RNA extracts, 500 ng of RNA were used. Samples were then mixed with 0.83 µL of the DNA Gel Loading Dye (6×) for native PAGE and 2.5 µL of the RNA Gel Loading Dye (2×) for denaturing PAGE before loading a total of 5 µL of mixture into the respective lanes. For the denaturing gels, samples were heated to 70 *^◦^*C for 10 min and subsequently cooled on ice for 3 min before loading. For native gels, one lane contained 1.25 µL of the GeneRuler 100 bp DNA Ladder, while 1.25 µL of the RiboRuler Low Range RNA Ladder was used for denaturing gels. For both native and denaturing gels, sample stacking was performed by running the gel at 40 V for 20 min immediately after loading. The native gels were then run at 150 V for 60 min, while the denaturing gels were run at 270 V for 60 min. All gels were run in TBE. Gels were stained after electrophoresis with 1 × SYBR Gold nucleic acid gel stain and imaged using a Syngene G:BOX Chemi XRQ gel documentation system. Lane intensities were quantified using ImageJ [113]. The full, un-cropped gel images are shown in Supplementary Fig. 48.

## Extended data figures

**Extended Data Figure 1:**
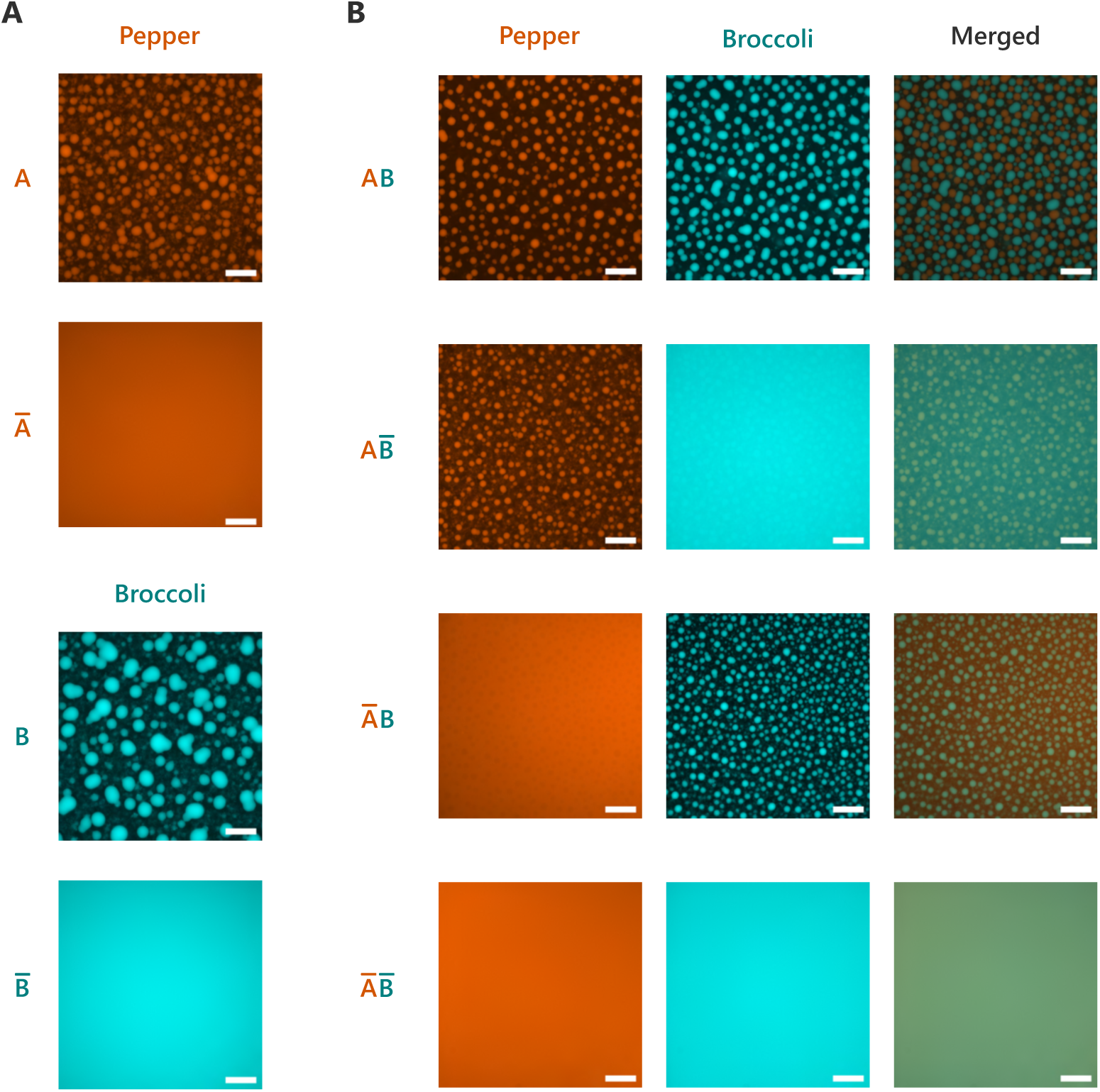
Epifluorescence micrographs of in vitro expression of single NSs and co-expression of two NSs. (**A**) Epifluorescence images of samples expressing single NSs, from top to bottom: A, Ā, B, B̄. (**B**) Epifluorescence images of samples co-expressing two NSs, from top to bottom: A+B, A +B̄, Ā +B, Ā +B̄. Images are shown in the Pepper fluorescent channel (NS A, left), in the Broccoli fluorescent channel (NS B, centre) and merging both channels (right). All samples were expressed from linear DNA templates with terminators (Methods) and imaged 4 hours after initiating the reactions. Scale bars, 100 µm.

**Extended Data Figure 2:**
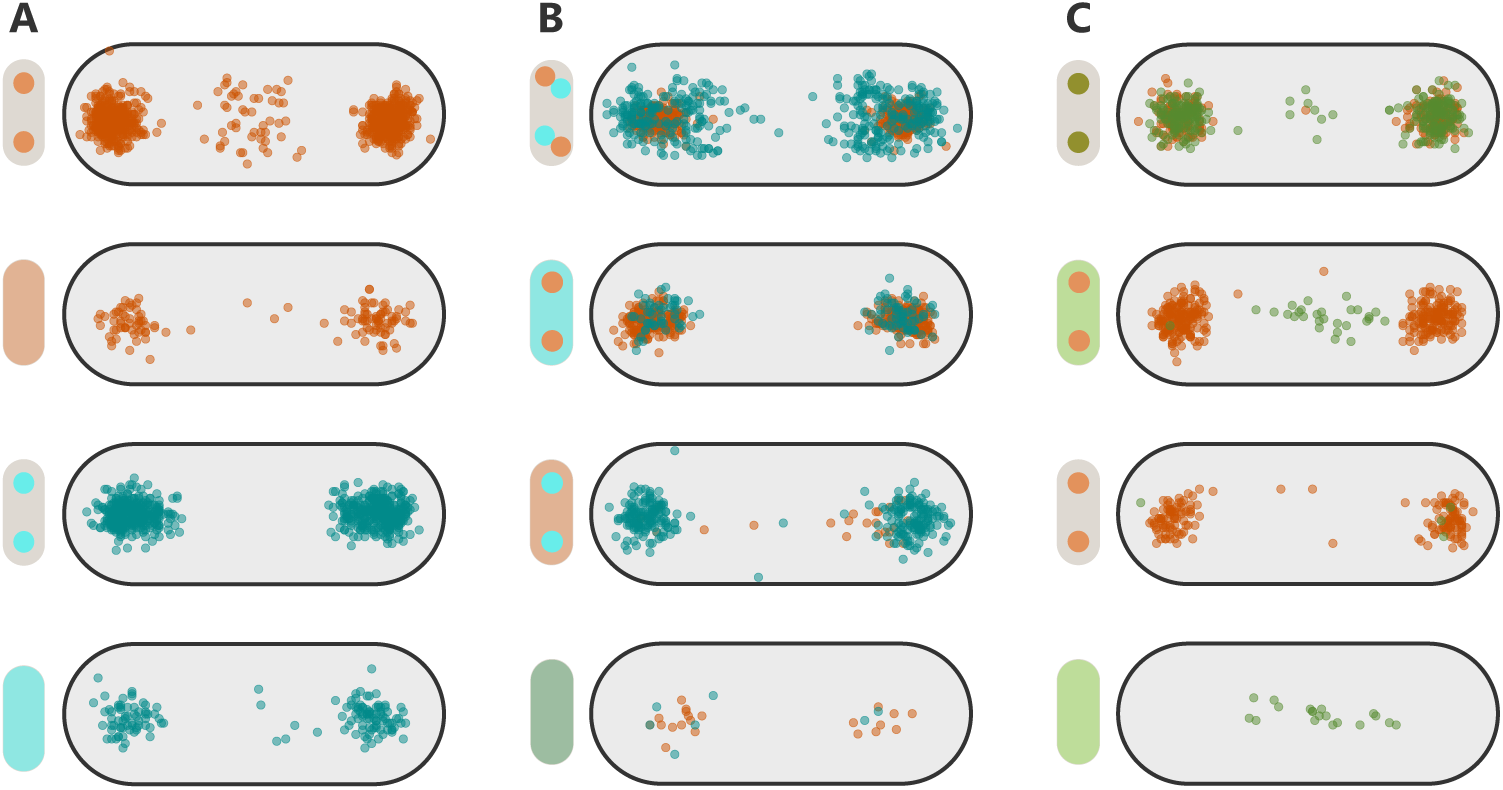
Locations of MLOs within bacteria. (**A**) Data for cells expressing a single NS, from top to bottom: A, Ā, B, B̄. NSs A and B display MLOs tightly localised at the poles. In NS A, some MLOs were also found near the centre of the cell. In contrast, NSs Ā and B̄ showed few clusters at the poles. (**B**) Data for cells expressing two NSs, from top to bottom: A+B (also shown in Fig. 2), A +B̄, Ā +B, Ā +B̄. In cells expressing A+B, A-MLOs are always found at the poles, while B NSs are sometimes closer to the centre of the cell. In cells expressing A+B̄ and B+Ā both specific MLOs and non-specific clusters are located at the poles. (**C**) Data relative to GFP capture experiments. From top to bottom, cells express A_AP3_ + GFP, A + GFP, A_AP3_ only, and GFP only. When A_AP3_ and GFP are expressed, we observe polar co-localisation of proteins and NSs. Expressing either NS A (in the presence of GFP) or A_AP3_ (in the absence of GFP) produces polar MLOs. No significant fluorescent localisation is observed if only GFP is expressed. Colour-coding is as follows. Orange: NS A (left and middle) and NS A_AP3_ (right); Cyan: NS B; Green: GFP. MLO locations were determined using segmentation and, as a result, not all MLOs identified by peak-picking were successfully located (Methods).

**Extended Data Figure 3:**
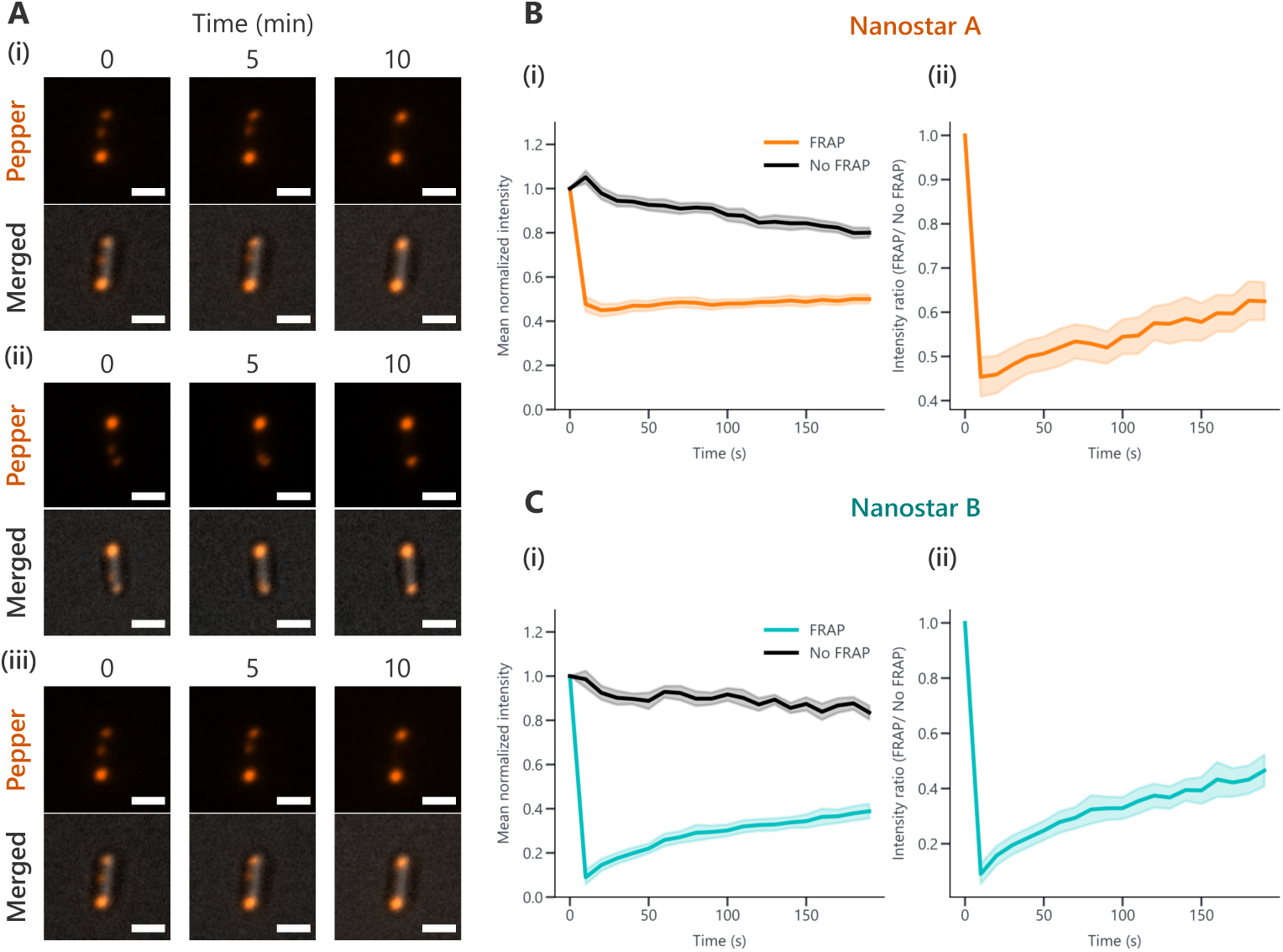
Fluidity of MLOs in *E. coli*. (**A**) Timelapse micrographs of three fusion events (i-iii) involving A MLOs. For each sequence, the top row shows confocal images and the bottom row shows merged confocal and bright-field images. Scale bars, 2 µm. (**B**) Fluorescence recovery after photo-bleaching (FRAP) for A MLOs. (i) FRAP curves collected for A MLOs. For each condensate, fluorescence intensity prior to photobleaching was normalised to 1. MLOs were obtained from three separate cultures, each starting from an individual single colony. The orange solid line (FRAP) is the mean normalised intensity of photobleached condensates (*n* = 25 MLOs), while the black solid line (No FRAP) is the mean normalised intensity of non-photobleached condensates (*n* = 15 MLOs). The decreasing trend in the intensity of non-photobleached condensates is caused by a degree of photobleaching occurring during imaging in the recovery phase. Shaded areas indicate the standard error of the mean. (ii) Ratio of the mean normalised intensity of photobleached A MLOs to the mean normalised intensity of non-photobleached A MLOs, correcting for photobleaching occurring in the recovery phase. The shaded area represents the standard error of the mean of the intensity ratios. (**C**) Data as in panel **B**, but for FRAP of B MLOs. (i) The cyan solid line (FRAP) is the mean normalised intensity of photobleached condensates (*n* = 12 MLOs), while the black solid line (No FRAP) is the mean normalised intensity of non-photobleached condensates (*n* = 8 MLOs).

**Extended Data Figure 4:**
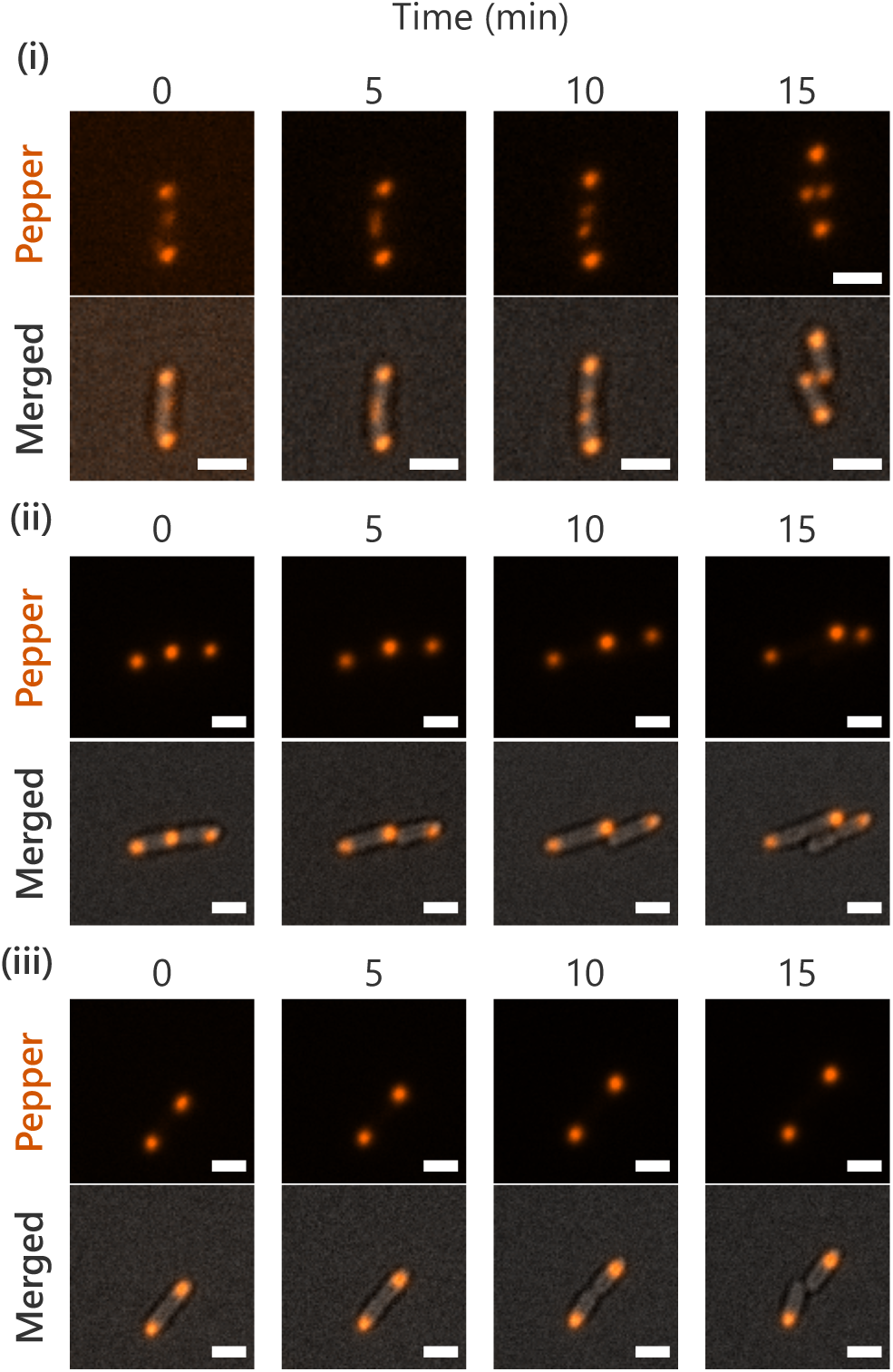
Timelapse micrographs of three division events in cells expressing A MLOs. For each sequence, the top row shows confocal images and the bottom row shows merged confocal and bright-field images. The three examples show the different behaviours observed upon division. A typical cell featuring two polar MLOs may grow a third condensate in the central section as it approaches division (i-ii). The central condensate can then be split between the two daughter cells, resulting in both having two polar condensates (i). Alternatively, the central condensate may be fully encased in one of the daughter cells, leaving the other with a single MLO. In other cases, a cell may divide without forming a central condensate, leaving both daughter cells with a single MLO. Scale bars, 2 µm.

**Extended Data Figure 5:**
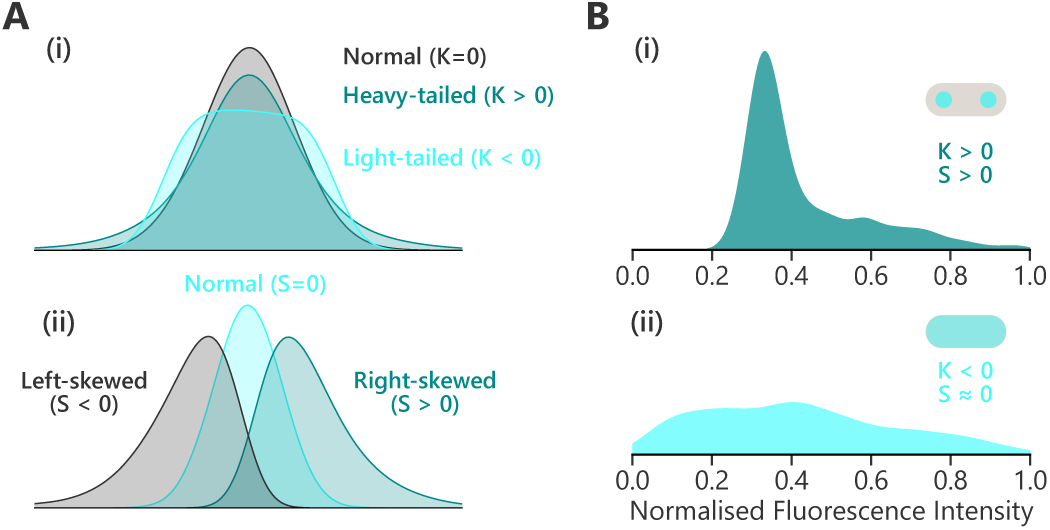
Interpretation of kurtosis and skewness of the pixel-fluorescence distributions of single cells, used to assess MLO formation. (**A**)(i) Examples of neutral, positive, and negative kurtosis (K). Positive kurtosis indicates a heavy-tailed distribution, while negative kurtosis indicates a light-tailed one. (ii) Examples of neutral, positive, and negative skewness (S). Positive skewness reflects a right-skewed distribution, whereas negative skewness reflects a left-skewed one. (**B**) Examples of pixel-fluorescence distributions for MLO-forming (i) and non-MLO-forming cells (ii). To obtain the distributions used for calculating kurtosis and skewness, fluorescence intensities are normalised such that 5% of the data points have a value of 1; these points are then excluded from the distribution. In MLO-forming cells (i), the distribution shows a sharp peak corresponding to background fluorescence and a long right-skewed tail originating from the brighter MLOs, resulting in positive kurtosis and skewness. In contrast, non-MLO-forming cells (ii) exhibit a more uniform fluorescence distribution across the cell, leading to near-zero skewness and negative kurtosis, which reflect the absence of distinct intensity peaks or heavy tails.

**Extended Data Figure 6:**
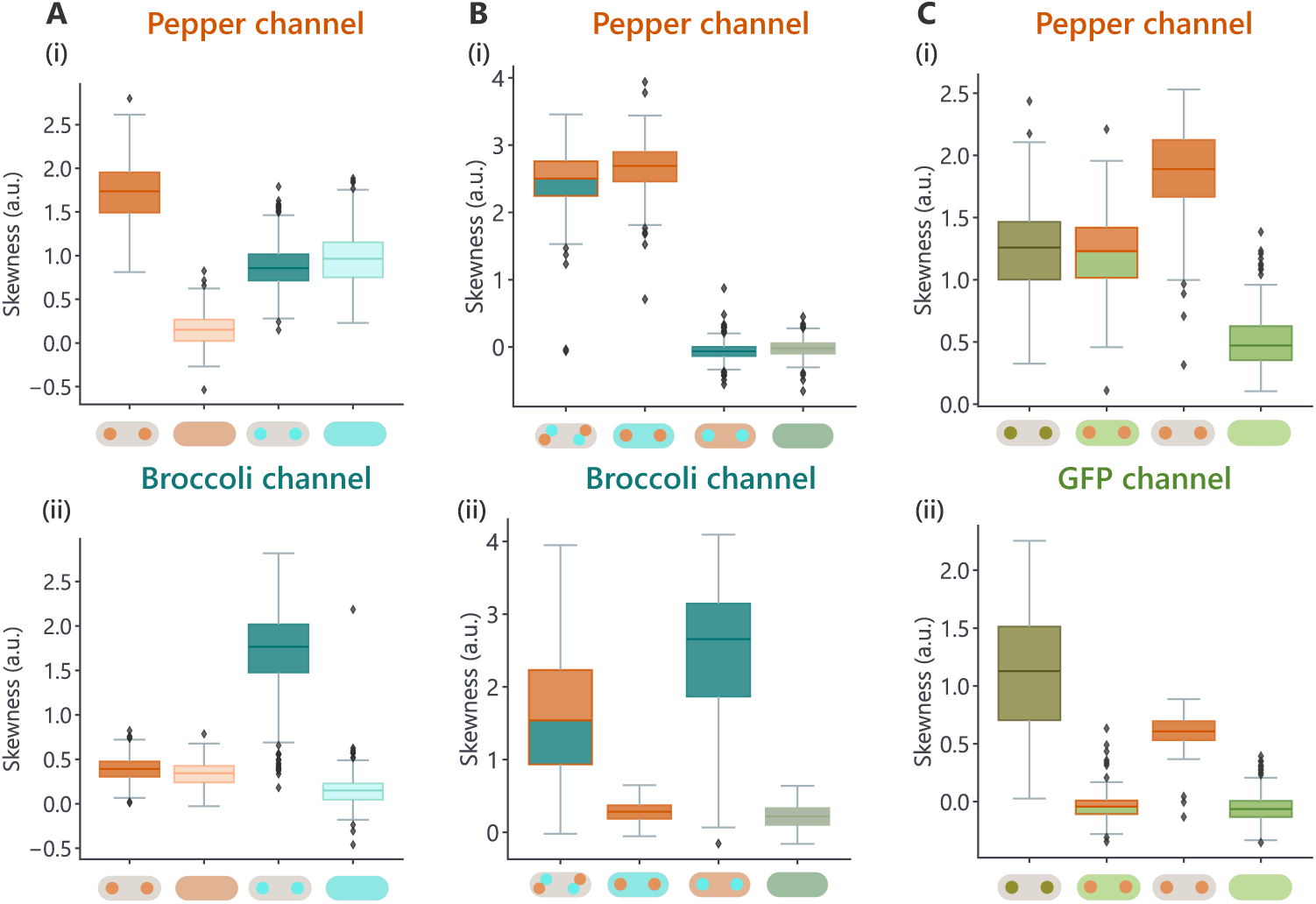
Comparing pixel-intensity-distribution skewness for NS-expressing *E. coli*. (**A**) Cells expressing single NSs, from left to right: A, Ā, B, and B̄. In the Pepper channel (i), NS A exhibits high skewness compared to NS Ā. Similarly, in the Broccoli channel (ii), NS B shows higher skewness than NS B̄. Cells that do not express NSs fluorescent in the respective channel display skewness values around 0.5. (**B**) Cells expressing two NSs, from left to right: A+B, A+B̄, Ā +B, and Ā +B̄. Cells expressing NS A show high skewness in the Pepper channel (i) relative to those expressing NS Ā. Likewise, cells expressing NS B exhibit high skewness compared to those expressing NS B̄ in the Broccoli channel (ii). (**C**) Experiments designed to demonstrate GFP capture. From left to right, cells express A_AP3_ + GFP, A + GFP, A_AP3_ only, and GFP only. Cells expressing NSs A or A_AP3_ display high skewness in the Pepper channel (i) compared to those not expressing either NS. Cells that capture GFP into MLOs exhibit high skewness in the GFP channel (ii) compared to cells where GFP remains evenly distributed throughout the cytoplasm. Cells lacking GFP entirely show intermediate skewness values around 0.6.

**Extended Data Figure 7:**
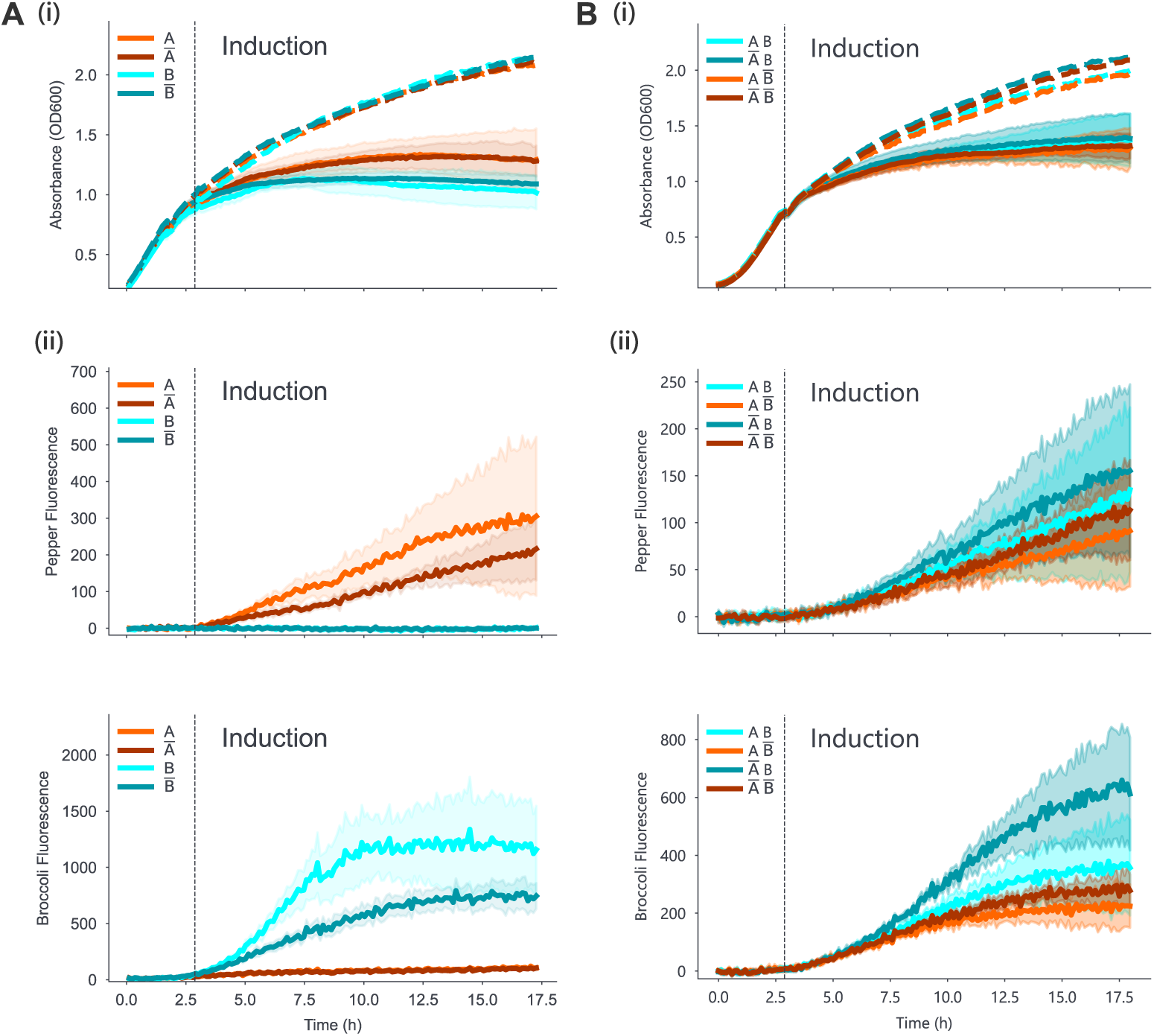
Growth and fluorescence curves of bulk *E. coli* cultures expressing single NSs or co-expressing two NSs. (**A**) Single NSs. (i) Optical Density (OD) curves for samples with (solid line) and without IPTG induction (dashed line, Supplementary Fig. 22). (ii) Fluorescence curves collected in the Pepper (NSs A and Ā) and Broccoli (NSs B and B̄) channels for induced samples. (**B**) Data as panel **A**, but for growth and fluorescence curves of cultures expressing two NSs (Supplementary Fig. 30). For each condition, measurements were performed for three biological replicates from separate single colonies, with three technical replicates studied per biological replicate. Solid lines represent the mean values across all replicates, while the shaded regions indicate the standard deviation across all replicates.

**Extended Data Figure 8:**
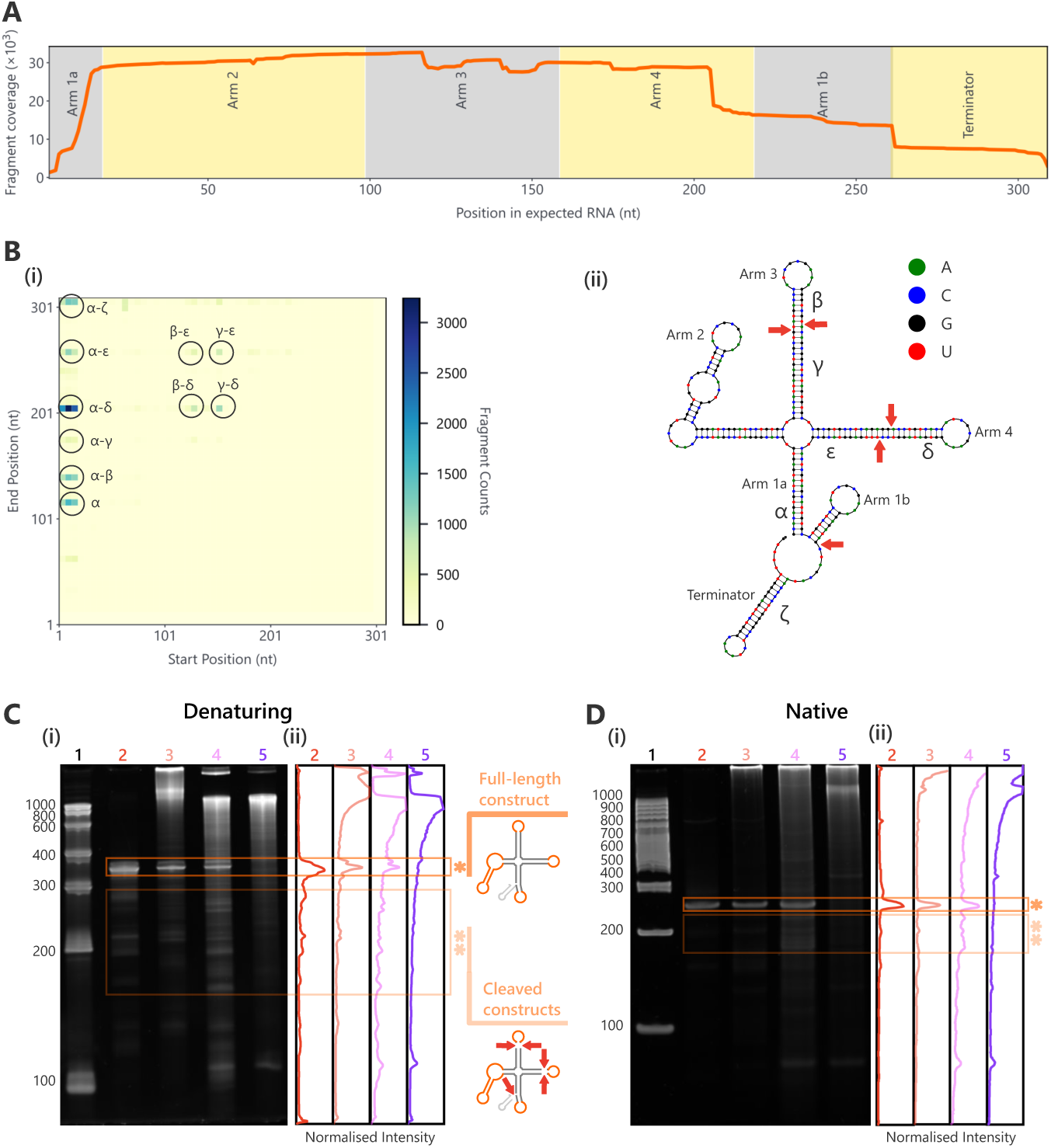
Sequencing and PAGE analysis of NS. **A.** (**A**-**B**) Nanopore sequencing of NS A as extracted from *E. coli*. (**A**) Coverage plot showing the number of aligned RNA fragments overlapping the expected NS A transcript. Coloured sections indicate the structural elements of the NS, with Arm 2 including the FLAP. The coverage profile confirms that the entire NS is transcribed, and highlights cleavage at specific sites ascribed to the post-transcriptional action of endonucleases. Fewer counts are obtained for the smaller fragments, which may be a result of bias in library preparation for sequencing [114, 115]. Relatively low coverage at the 5’ end (Arm 1a) is consistent with known basecalling limitations in direct RNA nanopore reads. Specifically, the helicase enzyme controlling translocation through the nanopore sits at a distance equivalent to 10-15 nt from the sensing region, causing the final 10-15 nt of a strand being sequenced to rapidly translocate once the 5’ is released by the enzyme, preventing accurate reading [112]. (**B**) (i) Binned scatter plot displaying the frequency of sequencing reads starting and ending at specified positions of the expected NS A sequence. High frequency locations mark RNA fragments, which are circled and labelled. (ii) NUPACK [99] schematic of the NS structure highlighting the fragments labelled in (i) and the preferential cutting sites (red arrows). The full-length transcript (without the terminator sequence: *α*-*ε*; with the terminator sequence: *α*-*ζ*) can be found. (**C**)(i) Denaturing PAGE of NS A samples. Lane legend: 1, RiboRuler Low Range RNA Ladder with marked band sizes (nt); 2, NS A transcribed in vitro from a linear DNA template; 3, NS A transcribed in vitro from a plasmid; 4, total RNA extract from *E. coli* induced to express NS A; 5, total RNA extract from uninduced *E. coli*. (ii) Lane intensity profiles. * marks the full-length construct, ** cleaved constructs. The full-length transcript is detected in lanes 2-4, with lane 4 (induced *E. coli*) also showing truncated products, consistent with sequencing. (**D**) Native PAGE of NS A samples, with lanes as in **C** except for using the GeneRuler 100 bp DNA Ladder in Lane 1. The full-size construct is detected in lanes 2-4, and constitutes the dominant species. High molecular weight material in lane 3 likely results from non-terminated transcripts, given the lower efficiency of termination in vitro [81], while those in lanes 4 and 5 are ascribed to mRNA.

**Extended Data Figure 9:**
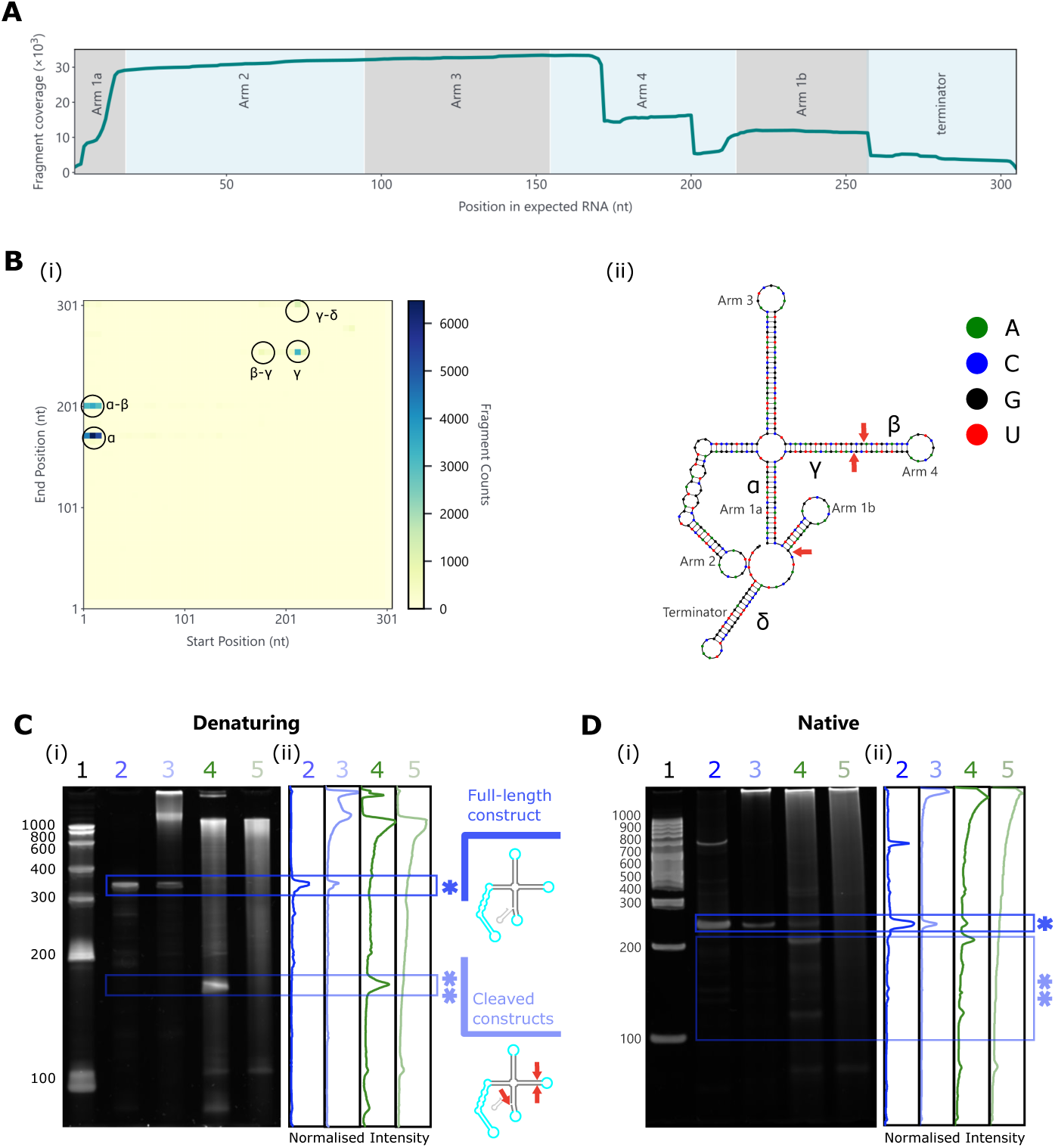
Sequencing and PAGE analysis of NS. **B.** (**A**-**B**) Nanopore sequencing of NS B as extracted from *E. coli*. (**A**) Coverage plot showing the number of aligned RNA fragments overlapping the expected NS B transcript. Coloured sections indicate the structural elements of the NS, with Arm 2 including the FLAP. The coverage profile confirms that the entire NS is transcribed, and highlights cleavage at specific sites ascribed to the post-transcriptional action of endonucleases. Fewer counts are obtained for the smaller fragments, which may be a result of bias in library preparation for sequencing [114, 115]. Relatively low coverage at the 5’ end (Arm 1a) is consistent with known basecalling limitations in direct RNA nanopore reads. Specifically, the helicase enzyme controlling translocation through the nanopore sits at a distance equivalent to 10-15 nt from the sensing region, causing the final 10-15 nt of a strand being sequenced to rapidly translocate once the 5’ is released by the enzyme, preventing accurate reading [112]. (**B**) (i) Binned scatter plot displaying the frequency of sequencing reads starting and ending at specified positions of the expected NS A sequence. High frequency locations mark RNA fragments, which are circled and labelled. (ii) NUPACK [99] schematic of the NS structure highlighting the fragments labelled in (i) and the preferential cutting sites (red arrows). The full-length transcript were not detected. (**C**)(i) Denaturing PAGE of NS B samples. Lane legend: 1, RiboRuler Low Range RNA Ladder with marked band sizes (nt); 2, NS B transcribed in vitro from a linear DNA template; 3, NS B transcribed in vitro from a plasmid; 4, total RNA extract from *E. coli* induced to express NS B; 5, total RNA extract from uninduced *E. coli*. (ii) Lane intensity profiles of the gels. * marks the full-length construct, ** cleaved constructs. The full-length transcript is detected in vitro (lanes 2 and 3) but not in induced *E. coli* (lane 4), consistent with sequencing. (**D**) Native PAGE of NS B samples, with lanes as in **C** except for using the GeneRuler 100 bp DNA Ladder in Lane 1. The full-size construct is detected in lane 4 in moderate quantities, likely nicked and stabilised by base pairing. The shorter dominant construct may lack the KL in Arm 4. High molecular weight material in lane 3 likely results from non-terminated transcripts, given the lower efficiency of termination in vitro [81], while those in lanes 4 and 5 are ascribed to mRNA.

**Extended Data Figure 10:**
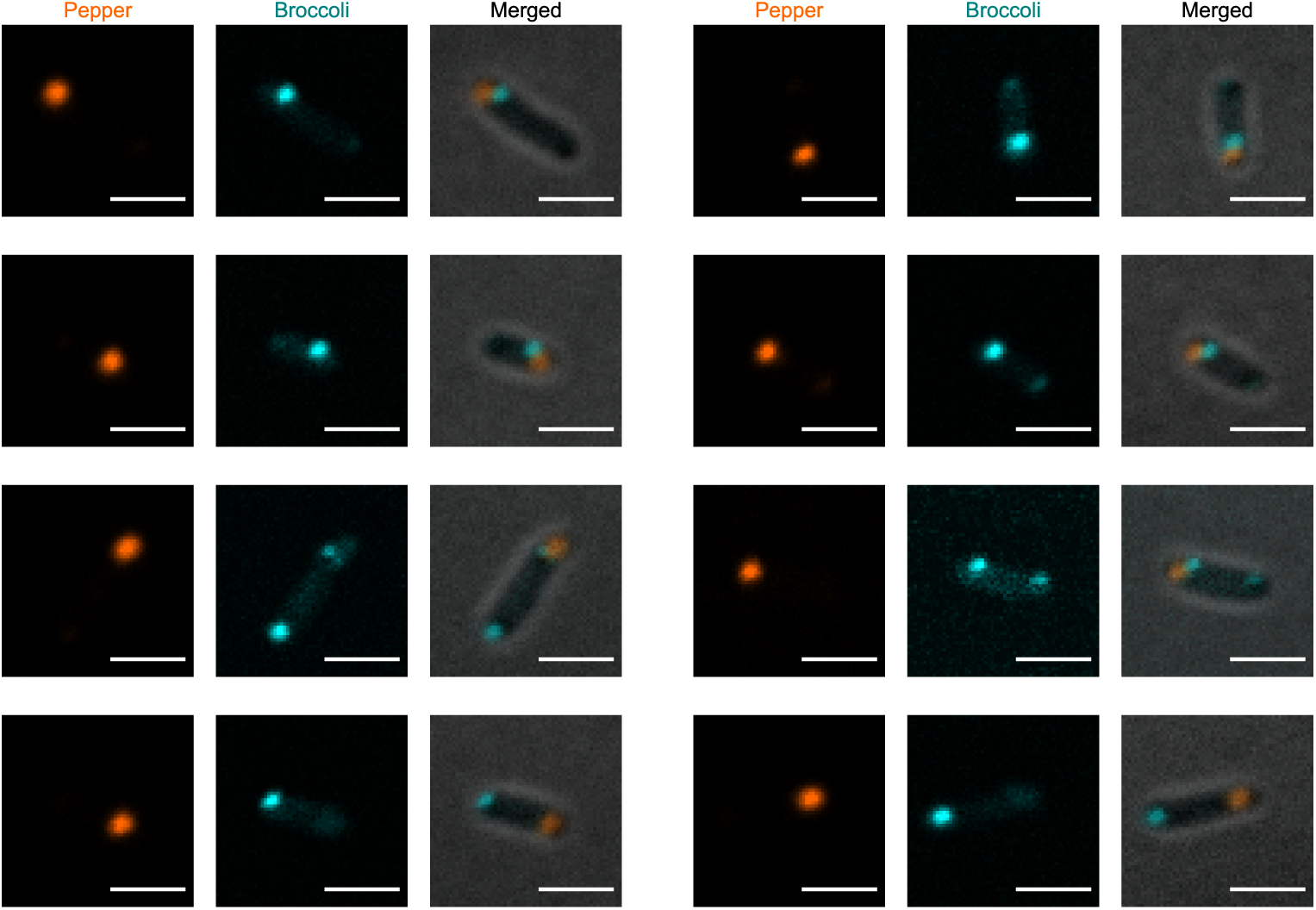
Zoomed in micrographs of cells co-expressing NSs A and. **B.** Zoomed in confocal micrographs recorded in the Pepper channel (NS A, left), the Broccoli channel (NS B, centre) and merging both fluorescence channels with bright-field images (right) of *E. coli* co-expressing NS A and NS B, 1 h after IPTG induction. Scale bars, 2 µm.

