## Supplementary Information for "Expression of nano-engineered RNA organelles in bacteria"

### Supplementary notes

#### Supplementary Note 1: Energetics of condensate formation within the nucleoid

The bacterial nucleoid is a dense, dynamic, and viscoelastic polymeric mesh composed primarily of DNA and associated proteins [1]. It occupies the central region of the cell due to ongoing replication and cohesion of sister chromosomes at the terminus region, which spatially anchors the nucleoid mid-cell [2]. Forming a condensate in the central region of the cell would create a void within the nucleoid, and thus have an associated energetic cost. To estimate this energy, we start from the work of Pelletier et al. [3]. In cells, the chromosome is compressed by the osmotic pressure exerted by cytosolic crowding agents that are excluded from the nucleoid. To imitate the effect of intracellular crowding, Pelletier et al. compressed the *E. coli* nucleoid in a microfluidic channel using a bead driven by optical tweezers, and showed the nucleoid's mechanical response can be modelled as an entropic spring

$$\frac{f}{A} = \left(\frac{R}{R_0}\right) - \left(\frac{R}{R_0}\right)^{-2},$$

where $f$ is the compression force, $A = -2.04$ pN is a scaling constant, $R_0 = 10.1\ \mu$m is the equilibrium length of the nucleoid in the microfluidic channel of width $w = 1.6\ \mu$m, and $R$ is the compressed nucleoid length. Here, lengths refer to the spatial extent of the nucleoid along the channel axis.
The compression pressure is thus given by

$$P = \frac{f}{w^2} = \frac{A}{w^2}\left[\left(\frac{R}{R_0}\right) - \left(\frac{R}{R_0}\right)^{-2}\right].$$

The energy required to create a void of volume $\Delta V$, given an initial nucleoid volume V, can be estimated as

$$\Delta E = \int_V^{V+\Delta V} P\,\mathrm{d}V = \int_V^{V+\Delta V} \frac{A}{w^2}\left[\left(\frac{R}{R_0}\right) - \left(\frac{R}{R_0}\right)^{-2}\right]\mathrm{d}V.$$

For a small $\Delta V$, one can assume that the pressure $P$ is unaffected by the volume change, which allows us to simply express the energy cost of void formation as $\Delta E \approx P\,\Delta V$.

Pelletier et al. report that that a force of 100 pN is required to compress the nucleoid to its in vivo size ($R/R_0 \approx 0.1$), resulting in $P \approx 80\,\mathrm{J\,m^{-3}}$ [3]. Using this value, we can estimate the energy cost of forming a spherical void of 50 nm in diameter as $\approx 4.2 \times 10^{-20}$ J, or $\approx 130\ k_\mathrm{B}T$. This energy cost is roughly 500-fold higher than the hybridisation free energy of the KL domain ($8.8 \times 10^{-23}$ J, estimated using Nupack [4]). This rough estimate suggests that the formation of nanostar clusters within the nuceloid region is strongly disfavoured, consistent with the observed, overwhelming tendency for pole accumulation.

Beyond energetic constraints, it is also likely that nucleoid presents a significant steric barrier for transport of nanostars and nanostar clusters across the cell, as previously reported for protein aggregates [5]. Combined with the energy cost of deforming the nucleoid, these steric effects would make it unlikely for nanostar clusters formed in nucleoid-free regions to diffuse into the nucleoid.

Finally, we note that cell crowding can be modulated, for instance by using sub-lethal concentrations of antibiotics [6]. Changing cell crowding should impact the mechanical characteristics of the nucleoid, which may in turn influence condensate localisation.

### Supplementary figures

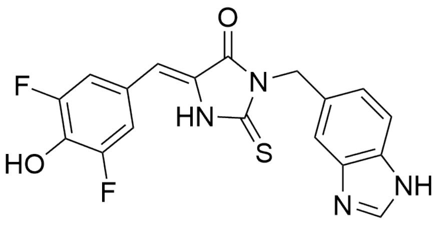


TBI (*Z*)-3-((1*H*-benzo[*d*]imidazol-5-yl)methyl)-5-(3,5-difluoro-4-hydroxybenzylidene)-2-thioxoimidazolidin-4-one

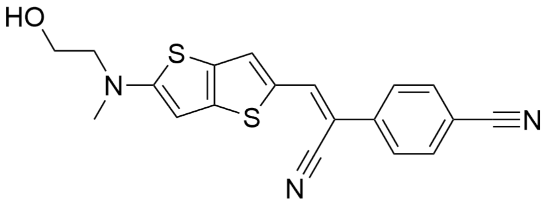


HBC620 ((*Z*)-4-(1-cyano-2-(5-((2-hydroxyethyl)(methyl)amino)thieno[3,2-*b*]thiophen-2-yl)vinyl)benzonitrile)

**Supplementary Figure 1**: **Structures and full chemical names of TBI** [7], **and HBC620** [8]

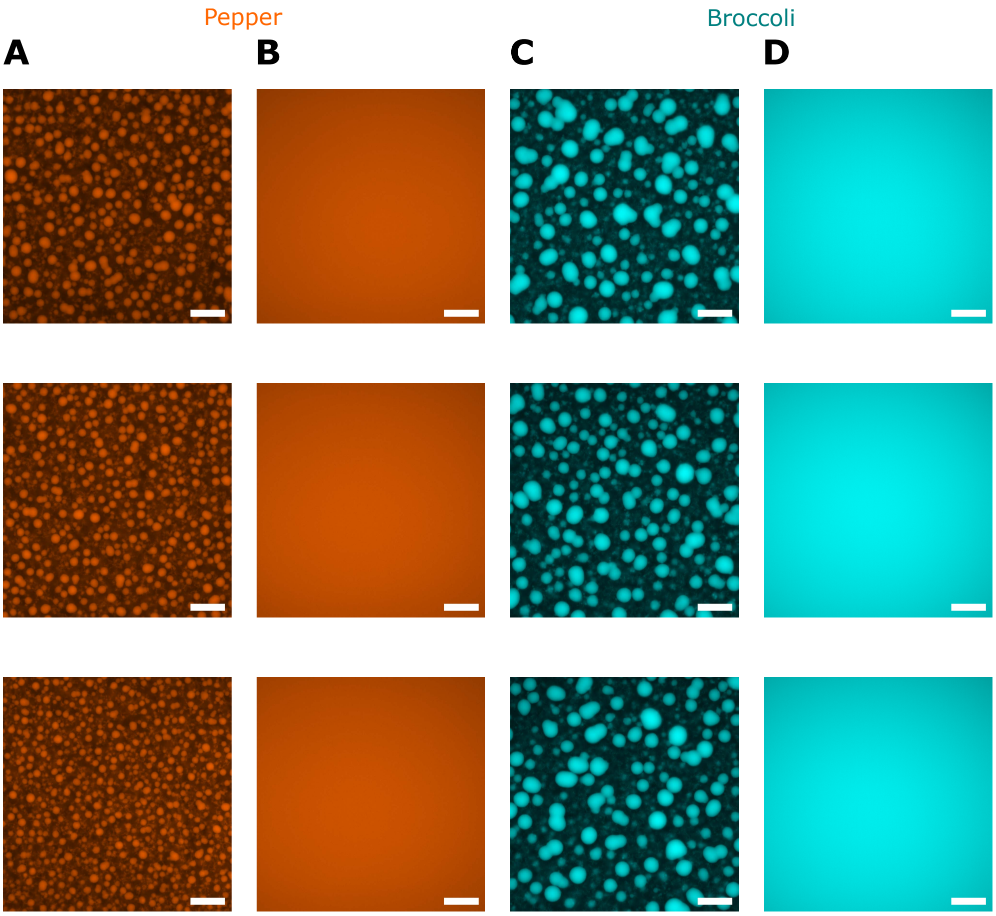


**Supplementary Figure 2**: **Epifluorescence micrographs of in vitro expression of single NSs.** Epifluorescence micrographs of (A) NS A, (B) NS $\bar{\text{A}}$, (C) NS B, and (D) NS $\bar{\text{B}}$, after 4 h incubation. All NSs were expressed from linear DNA templates with terminators. Scale bars, 100 µm.

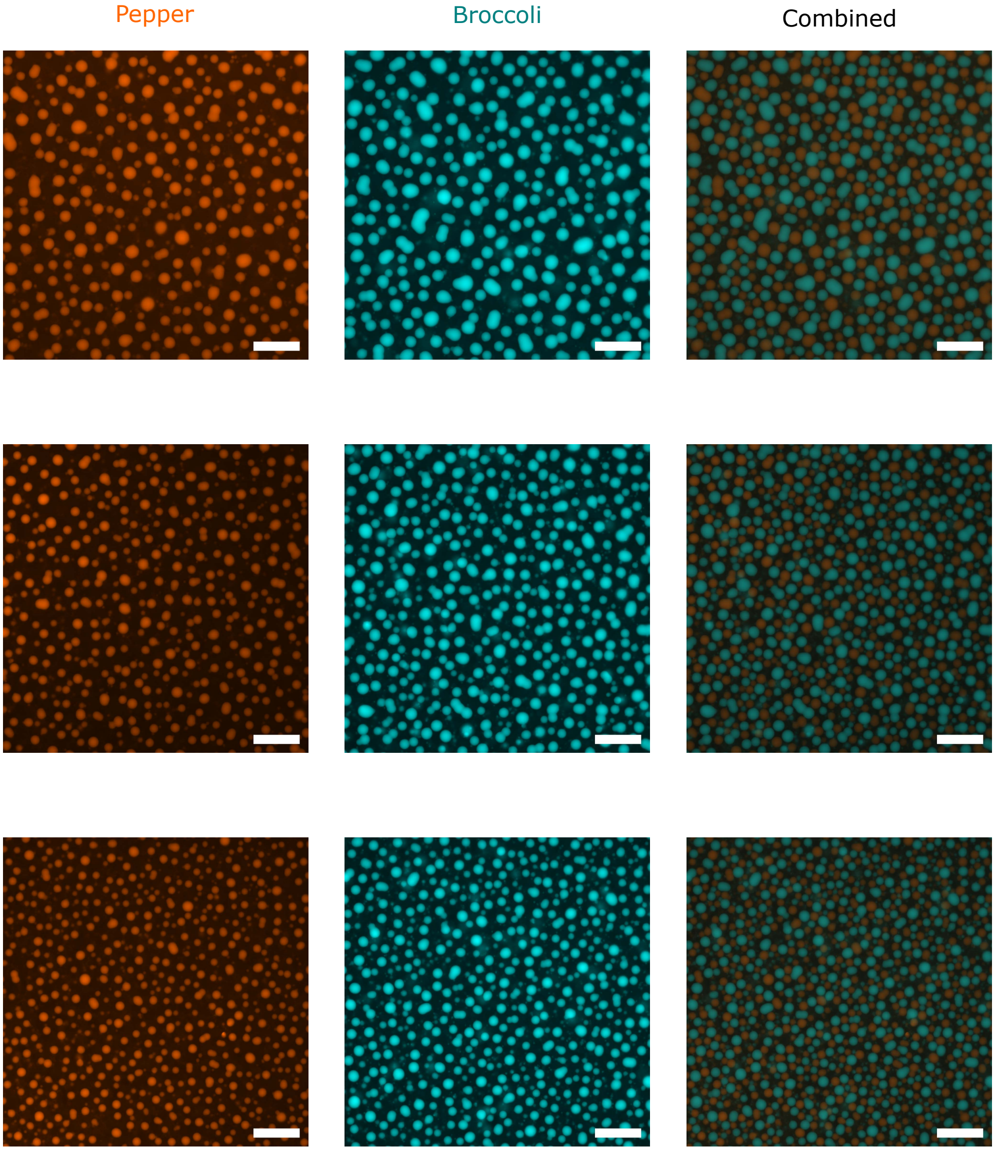


**Supplementary Figure 3: Epifluorescence micrographs of *in vitro* co-expression of NS A and NS B.** The micrographs were collected 4 hours from the beginning of transcription. Images are shown in the Pepper fluorescent channel (NS A, left), in the Broccoli fluorescent channel (NS B, centre) and merging both channels (right). From the merged micrographs one can verify that A and B condensates are orthogonal and non-mixing, as designed. All NSs were expressed from linear DNA templates with terminators. Scale bars, 100 µm.

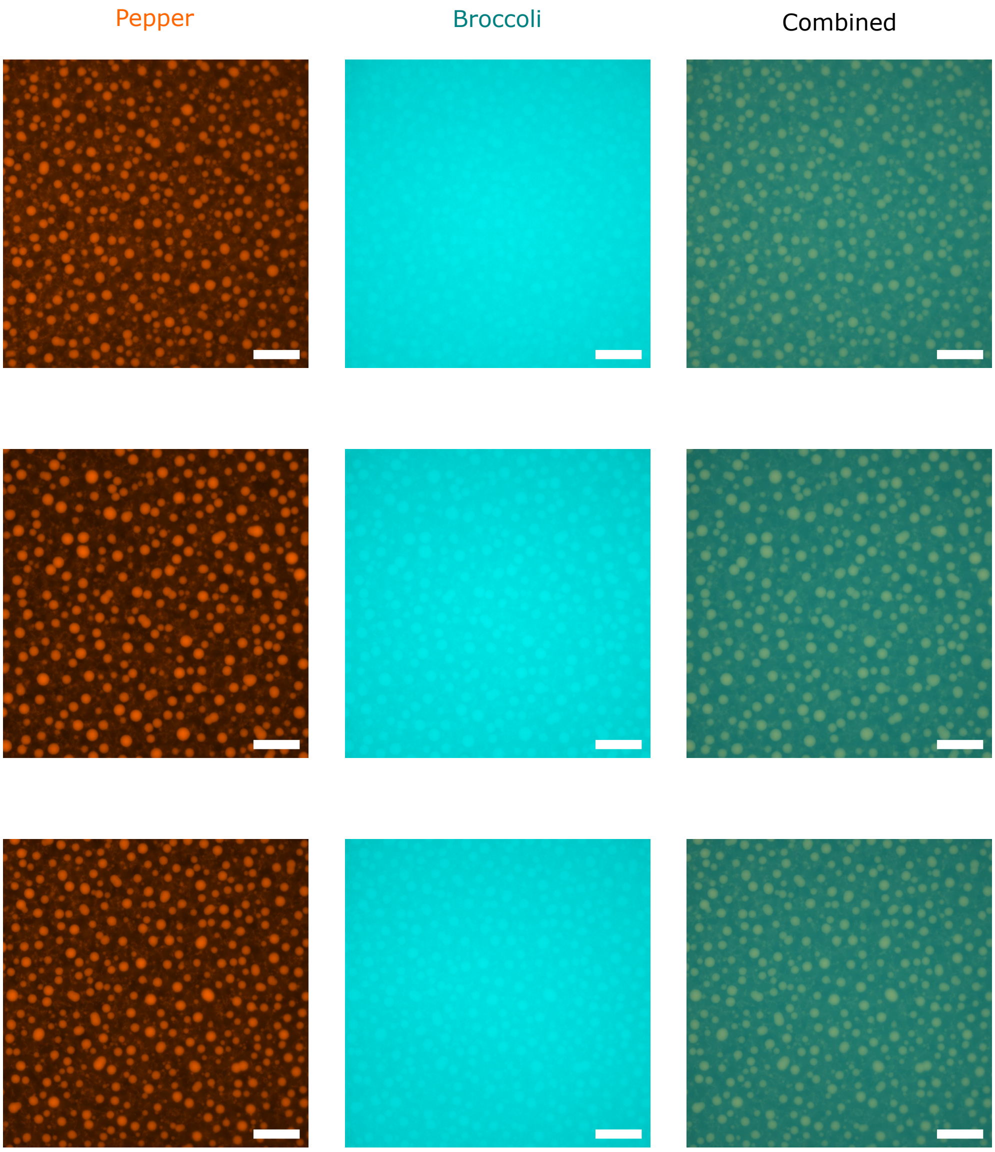


**Supplementary Figure 4: Epifluorescence micrographs of in vitro co-expression of NS A and NS $\bar{\mathrm{B}}$.** The micrographs were collected 4 hours from the beginning of transcription. Images are shown in the Pepper fluorescent channel (NS A, left), in the Broccoli fluorescent channel (NS $\bar{\mathrm{B}}$, centre) and merging both channels (right). NS $\bar{\mathrm{B}}$, featuring scrambled Kissing Loops, produces uniform fluorescence, as expected. All NSs were expressed from linear DNA templates with terminators. Scale bars, 100 µm.

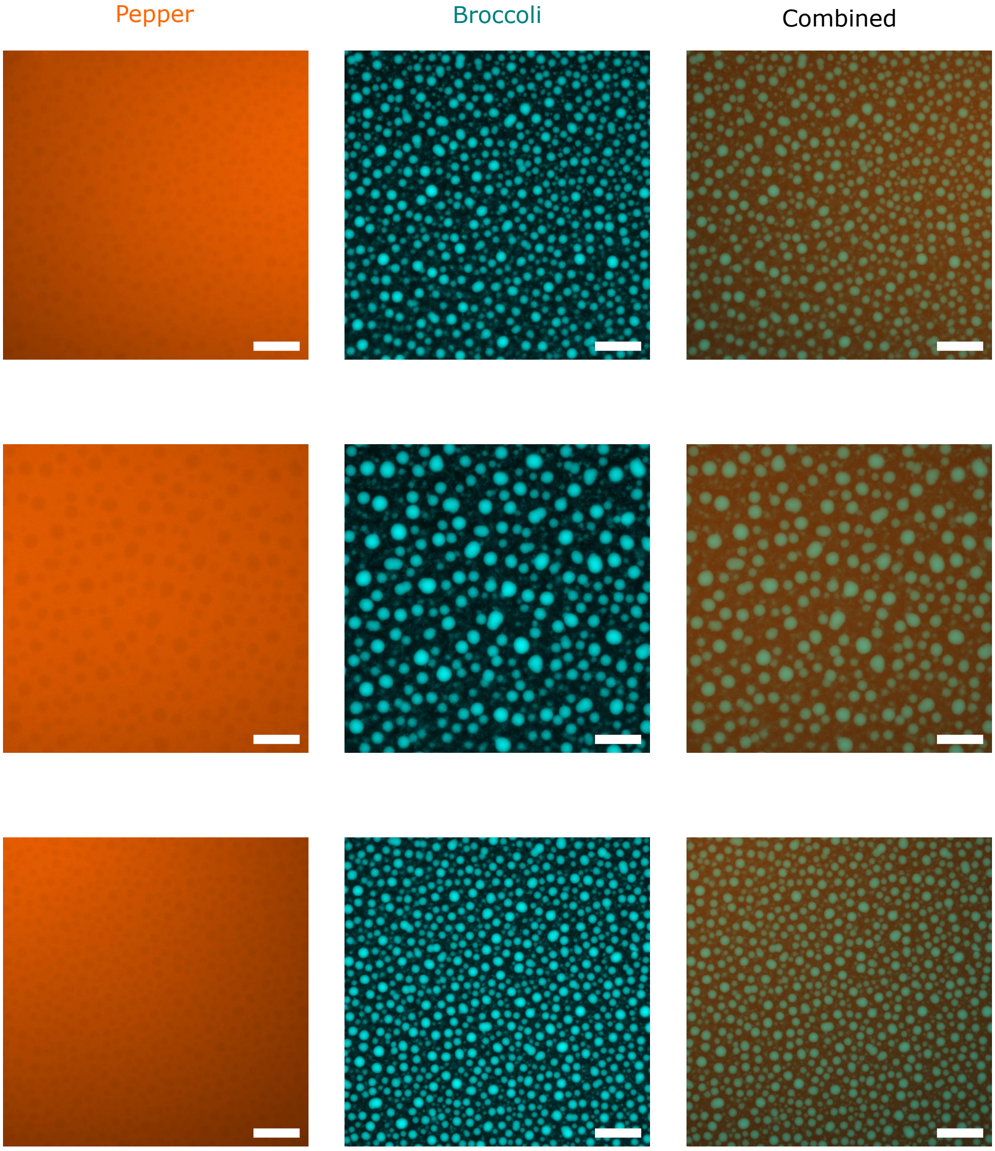


**Supplementary Figure 5**: **Epifluorescence micrographs of in vitro co-expression of NS $\bar{A}$ and NS B.** The micrographs were collected 4 hours from the beginning of transcription. Images are shown in the Pepper fluorescent channel (NS $\bar{A}$, left), in the Broccoli fluorescent channel (NS B, centre) and merging both channels (right). NS $\bar{A}$, featuring scrambled Kissing Loops, produces uniform fluorescence, as expected. All NSs were expressed from linear DNA templates with terminators. Scale bars, 100 µm.

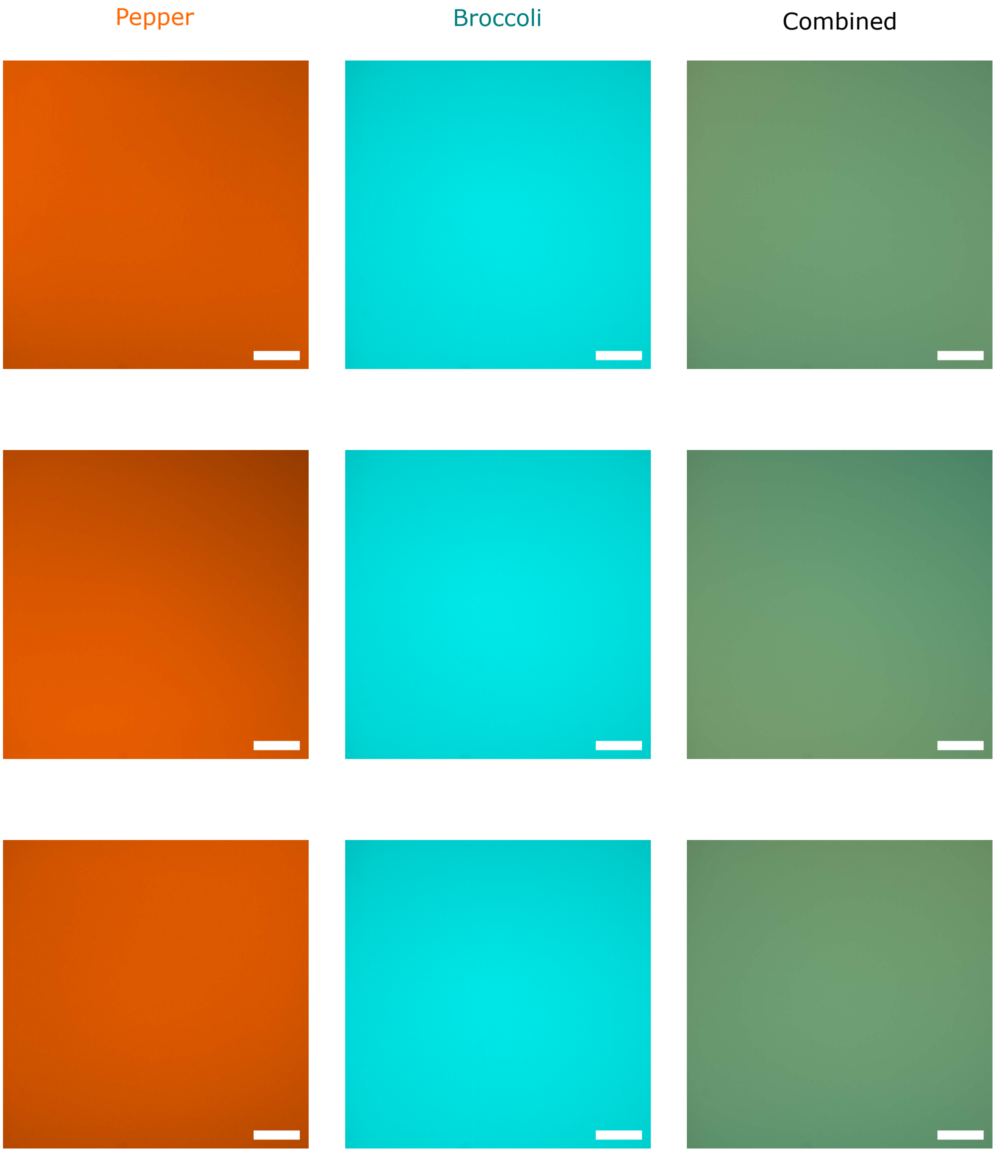


**Supplementary Figure 6: Epifluorescence micrographs of in vitro co-expression of NS $\bar{A}$ and NS $\bar{B}$.** The micrographs were collected 4 hours from the beginning of transcription. Images are shown in the Pepper fluorescent channel (NS $\bar{A}$, left), in the Broccoli fluorescent channel (NS $\bar{B}$, centre) and merging both channels (right). Both NSs feature scrambled Kissing Loops sequences and produce uniform fluorescence, as expected. All NSs were expressed from linear DNA templates with terminators. Scale bars, 100 µm.

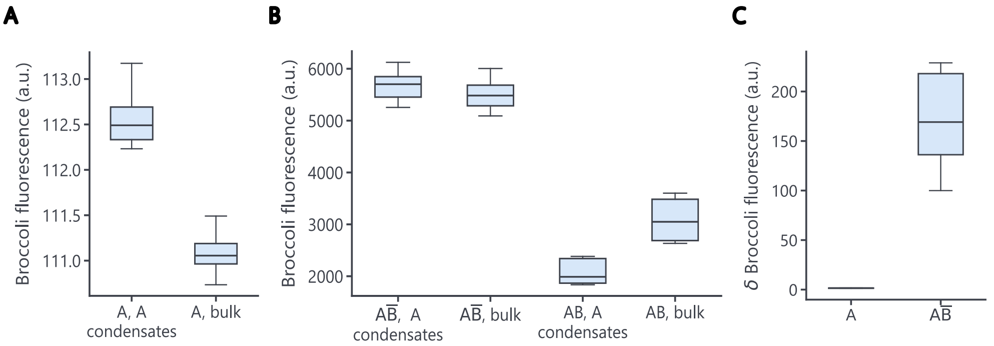


**Supplementary Figure 7**: **Quantification of the Broccoli signal observed in A-type condensates.** (**A**) Mean Broccoli fluorescence within and outside (bulk phase) NS A condensates in NS A-only samples. The Broccoli signal in the denser NS A condensates differs only slightly from that in the bulk phase, indicating that fluorescence bleed-through from the Pepper channel into the Broccoli channel is minimal. (**B**) Mean Broccoli fluorescence within and outside NS A condensates in A+$\bar{\text{B}}$ and A+B samples. In A+$\bar{\text{B}}$ samples, Broccoli fluorescence is higher within NS A condensates than in the bulk, whereas in A+B samples, fluorescence is higher in the bulk phase. The latter observation is consistent with excluded volume effects that lower the concentration of NS B within A condensates relative to the bulk phase. (**C**) Difference, $\delta$, in mean Broccoli fluorescence intensity between NS A condensates and the surrounding bulk phase in A-only and A+$\bar{\text{B}}$ samples. The larger $\delta$ in A+$\bar{\text{B}}$ samples suggests accumulation of NS $\bar{\text{B}}$ within A condensates, likely due to weak affinity between the scrambled KLs of NS $\bar{\text{B}}$ and the KLs of NS A.

(i)

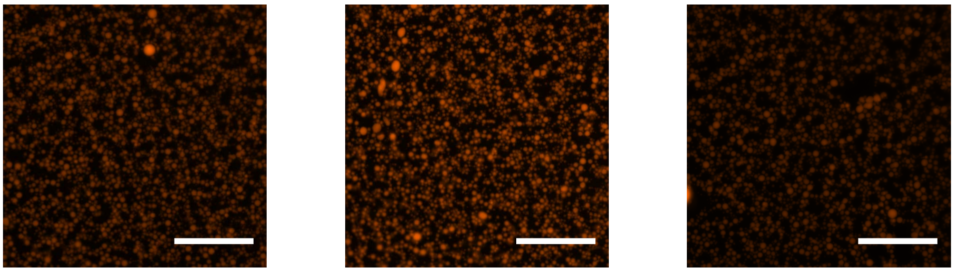

(ii)

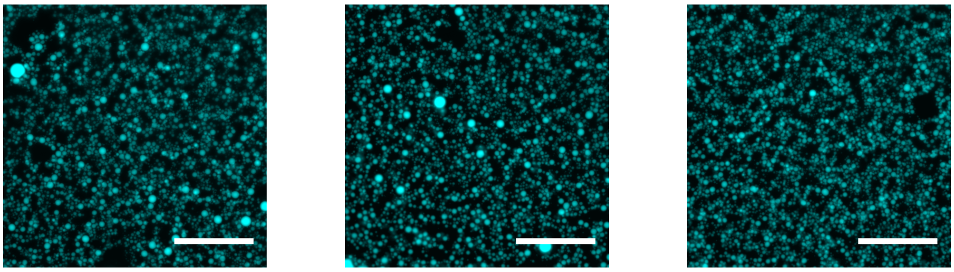

**Supplementary Figure 8**: **Epifluorescence micrographs of in vitro expressed NS incubated in buffer mimicking the intracellular ionic environment of *E.coli*.** Epifluorescence micrographs of (i) NS A. (ii) NS B, in buffer containing 1 mM $MgCl_2$ and 200 mM KCl. Scale bars, 200 µm.

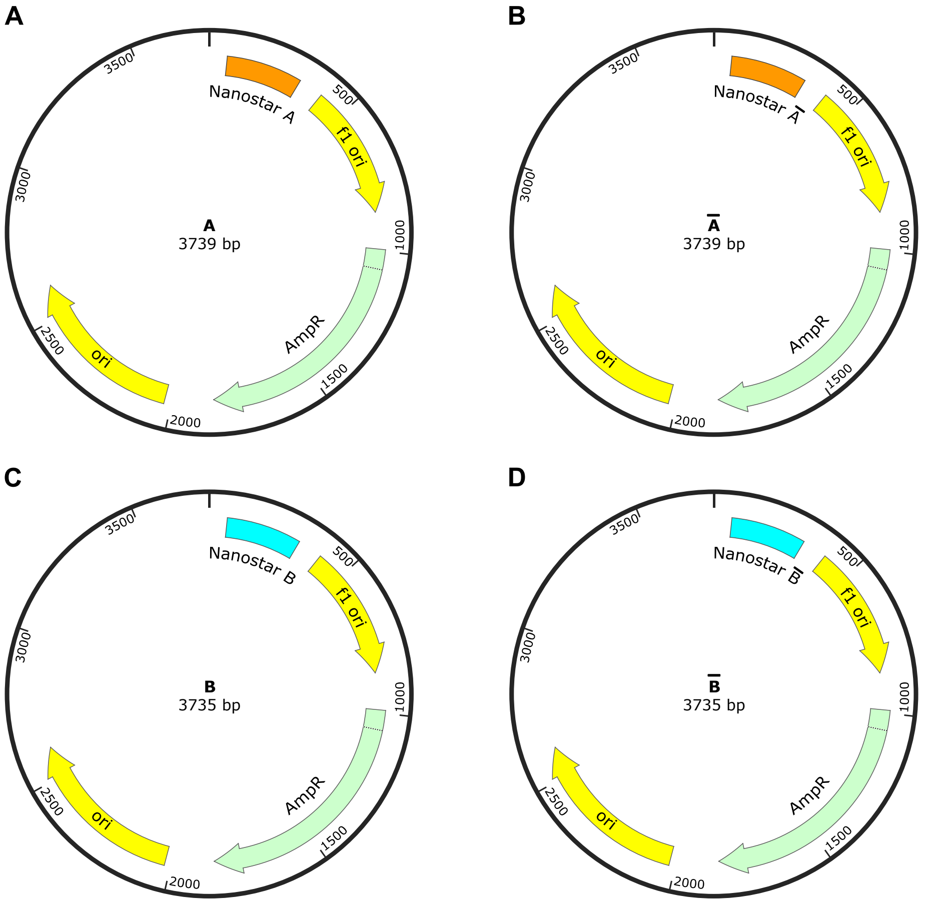


**Supplementary Figure 9**: **Plasmid maps** of plasmids expressing (**A**) NS A, (**B**) NS Ā, (**C**) NS B, (**D**) NS B̄. The plasmid maps were created with Snapgene (version 8.1).

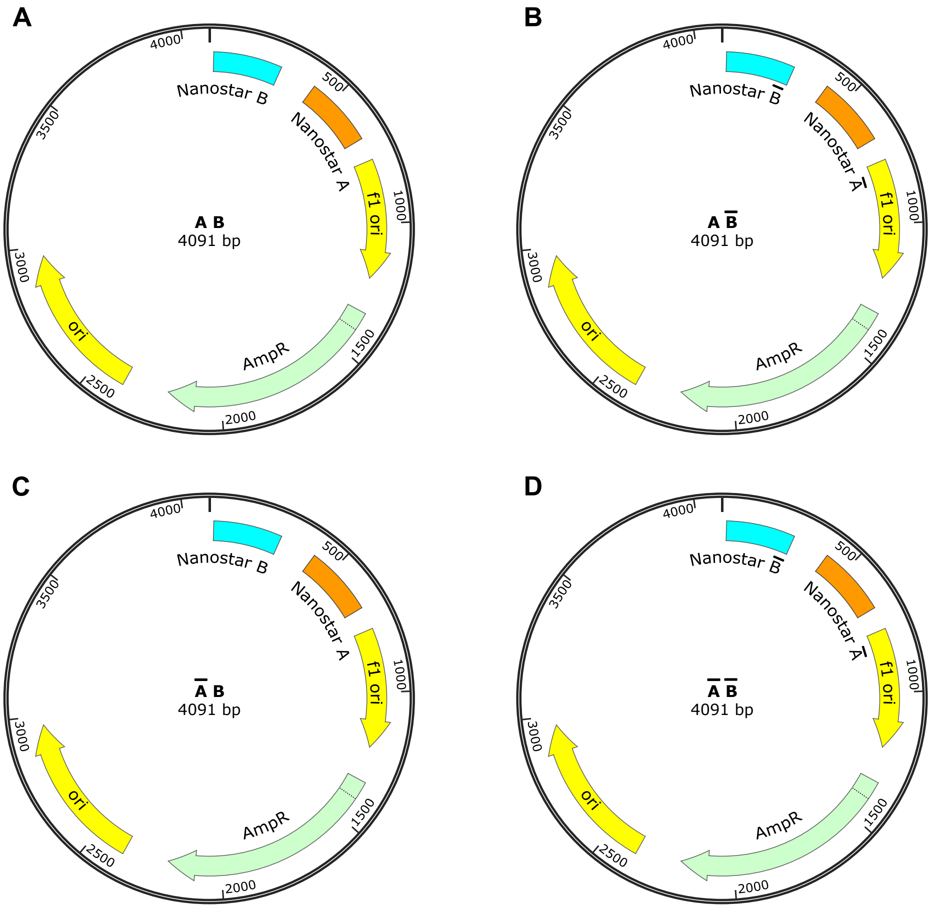


**Supplementary Figure 10**: **Plasmid maps** of plasmid expressing (**A**) NSs A and B, (**B**) NSs A and B̄, (**C**) NSs Ā and B, (**D**) NSs Ā and B̄. The plasmid maps were created with Snapgene (version 8.1).

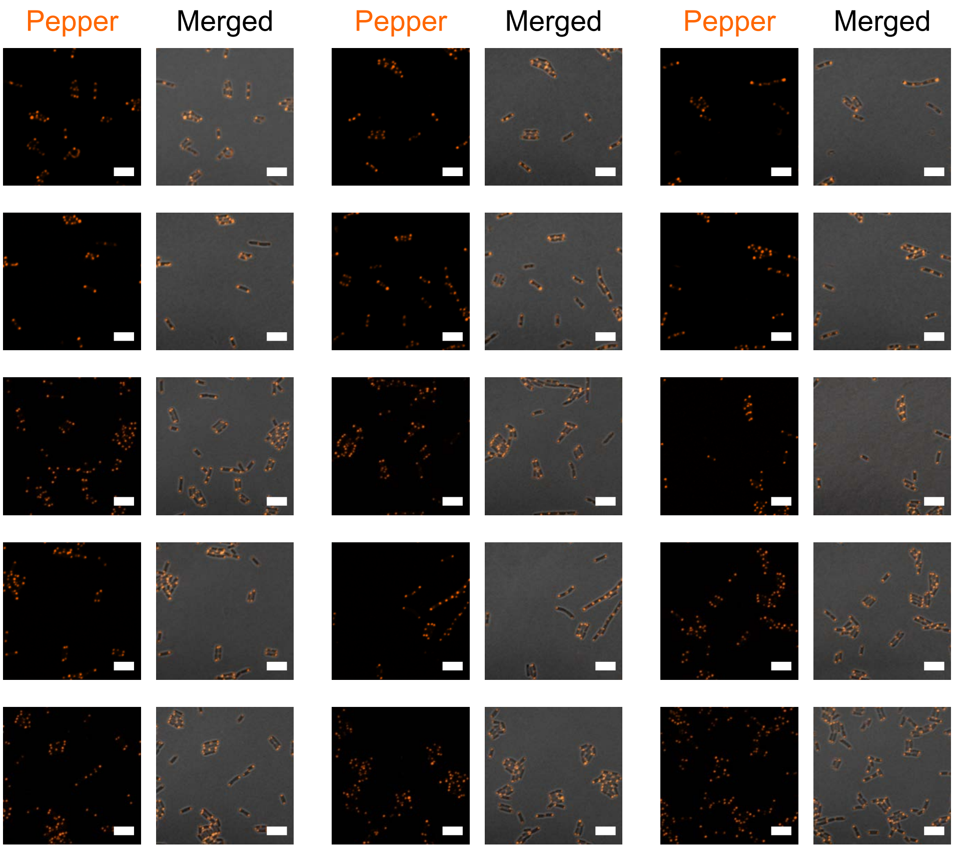


**Supplementary Figure 11**: **Additional micrographs of *E. coli* expressing NS A.** Confocal micrographs (left) and merged confocal/bright-field images (right) of *E. coli* expressing NS A, 1 h after IPTG induction. Scale bars, 5 µm. Micrographs were obtained from three separate cultures, each starting from an individual single colony.

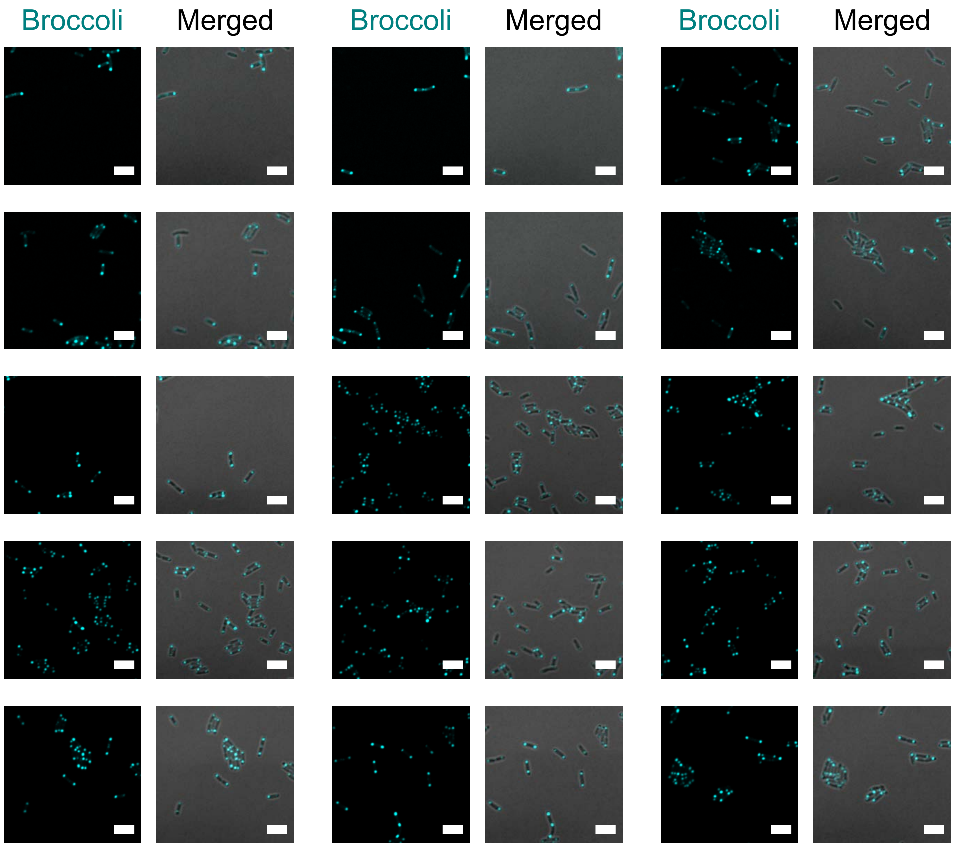


**Supplementary Figure 12**: **Additional micrographs of *E. coli* expressing NS B.** Confocal micrographs (left) and merged confocal/bright-field images (right) of *E. coli* expressing NS B, 1 h after IPTG induction. Scale bars, 5 µm. Micrographs were obtained from three separate cultures, each starting from an individual single colony.

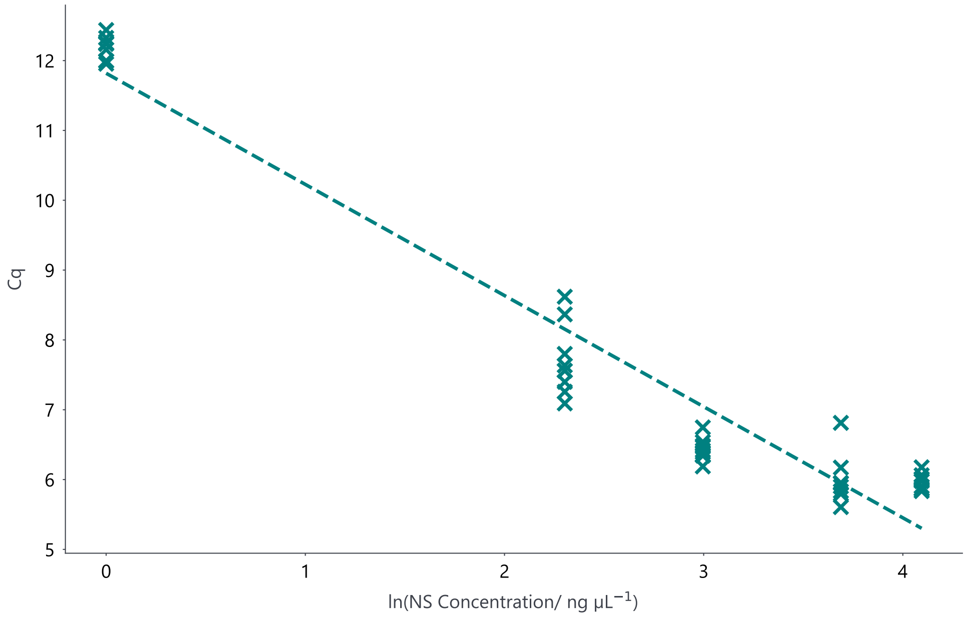


**Supplementary Figure 13**: **RT-qPCR Nanostar concentration calibration curve.** RT-qPCR calibration curve showing the relationship between $Cq$ values and the natural logarithm of RNA concentration for NS B. Linear fitting produced $Cq = -1.59 \times \ln(\text{NS concentration/ng}\,\mu\text{L}^{-1}) + 11.8$. See Methods for experimental details.

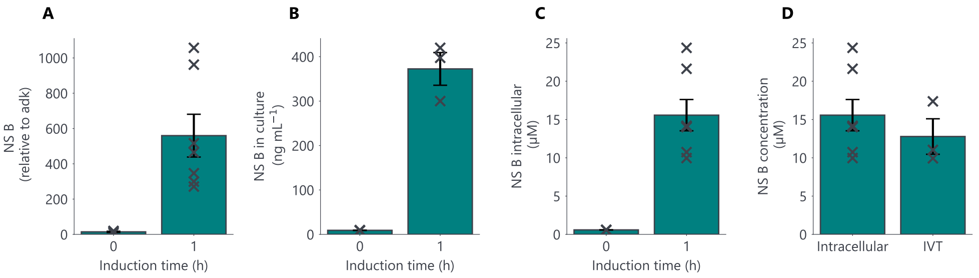


**Supplementary Figure 14**: **Concentration of NSs expressed in *E. coli* as analysed by RT-qPCR.** (**A**) Bar graphs of the relative concentration of NS B, normalised to the housekeeping gene *adk*. (**B**) Absolute concentrations of NS B per mL of cell culture. (**C**) Intracellular concentrations of NS B, estimated by dividing the data in (**B**) by the overall intracellular volume. To compute the latter, we estimated the total number of cells in the culture using optical density (OD) ($1\,\text{OD} = (1.0 \pm 0.3) \times 10^8$ colony forming units $\text{mL}^{-1}$, based on 4 independent measurements), and multiplied by the expected single-cell volume of 3 $\mu m^3$, obtained by approximating the cell as a cuboid with dimensions 1 µm × 1 µm × 3 µm (see Methods). The height of the bars are the mean values of three technical replicates (RT-qPCR measurements) performed on each biological replicate grown from independent colonies. For the 1 h induction condition, seven biological replicates were analysed; for the 0 h induction condition, three biological replicates were analyzed. Error bars indicate standard error of the mean. (**D**) Side-by-side comparison of the intracellular concentration of NS B and that of NS B expressed in vitro 3 h after starting the reaction. For RT-qPCR, a calibration curve produced using known concentrations of purified RNA was used to calculate the concentrations in (**B**) and (**C**) (Supplementary Fig. 13). NS concentration in the in vitro sample was determined with UV absorbance using a Nanodrop ONE spectrophotometer, following RNA purification. We note that, while RT-qPCR is widely adopted for measuring intracellular RNA concentrations due to its sensitivity, specificity, and relative ease of use, outcomes can still be influenced by variability in RNA extraction, secondary structure, and enzymatic processing (Extended Data Figs. 8A-B, 9A-B) and post-transcriptional modifications.

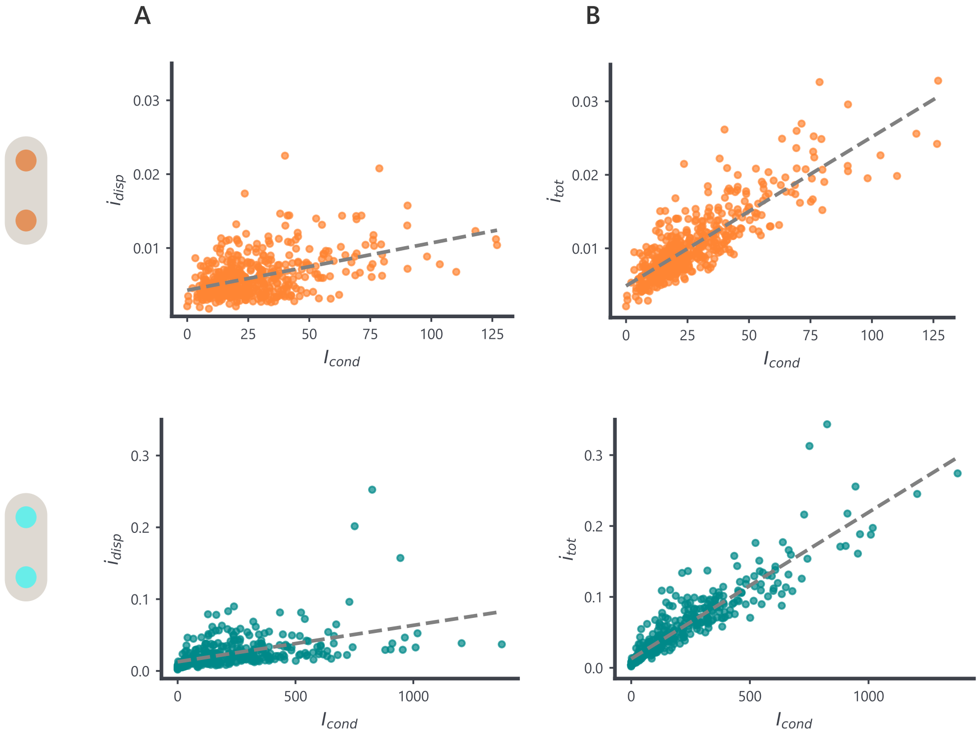


**Supplementary Figure 15**: **Probing of the thermodynamic behaviour of NSs within cells.** (**A**) Scatter plot of the average fluorescence intensity measured outside condensates ($i_{\rm disp}$) against the integrated fluorescence inside segmented condensates ($I_{\rm cond}$). $i_{\rm disp}$ is proportional to the concentration of dispersed NSs, while $I_{\rm cond}$ is expected to be proportional to condensate volume, under the assumption of uniform NS density and brightness in the condensed phase. Linear fits (dashed gray line) are shown with the following parameters: NS A (top) ($y = 6.4{\times}10^{-5}x + 4.3{\times}10^{-3}$, $R^2 = 0.21$) and NS B (bottom) ($y = 5.1{\times}10^{-5}x + 1.3{\times}10^{-2}$, $R^2 = 0.24$). Weak correlation between $i_{\rm disp}$ and $I_{\rm cond}$ is consistent with NS concentration in the dispersed phase being independent on condensate volume, as expected at thermodynamic equilibrium. (**B**) Scatter of the average fluorescence of the whole cell, $i_{\rm tot}$, proportional to the overall NS concentration, against $I_{\rm cond}$. Linear fits (dashed gray line) are shown with the following parameters: NS A (top) ($y = 2.0{\times}10^{-4}x + 4.9{\times}10^{-3}$, $R^2 = 0.70$) and NS B (bottom) ($y = 2.1{\times}10^{-4}x + 1.3{\times}10^{-2}$, $R^2 = 0.83$). At thermodynamic equilibrium, a linear dependence of condensate volume on overall NS concentration is consistent with the lever rule.

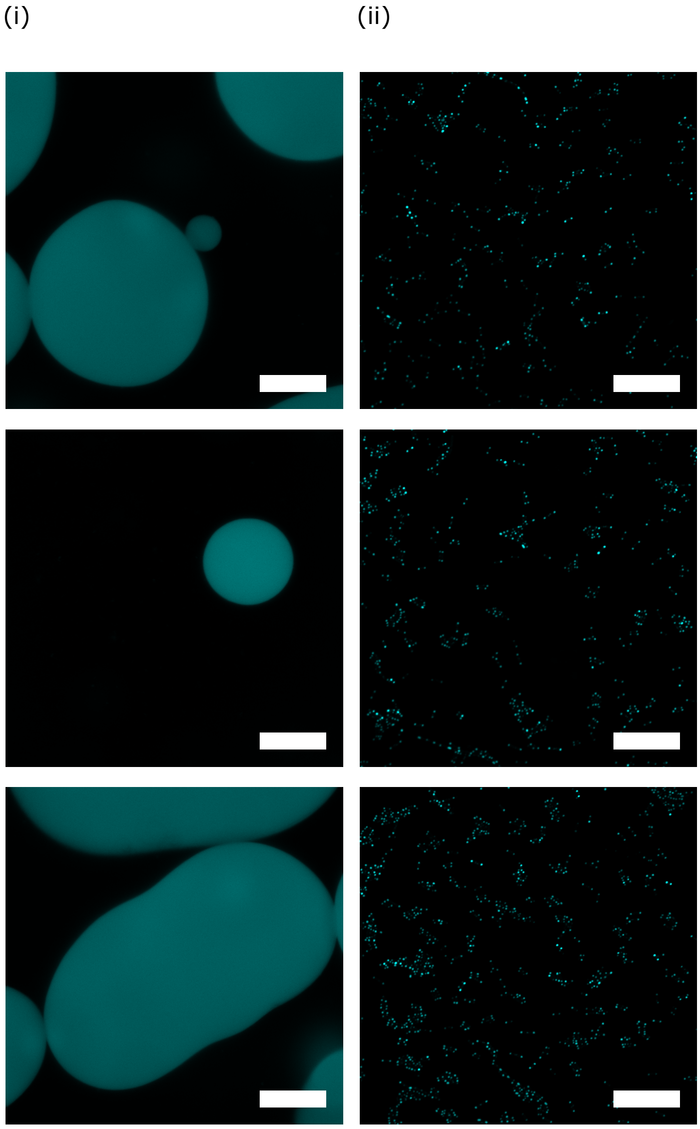


**Supplementary Figure 16**: **Confocal micrographs of in vitro and in vivo NS B condensates stained using the same TBI concentration (10 µM).** The micrographs were obtained using the same imaging protocol and subjected to the same post-processing settings (Methods). The segmented images exhibit comparable fluorescence, with the mean pixel intensity of NS B condensates expressed in vitro (**i**) being 23% higher than that of NS B condensates expressed in *E. coli* (**ii**). This suggests similarities in FLAP response and overall NS concentration in the condensed phase. Scale bars, 20 µm.

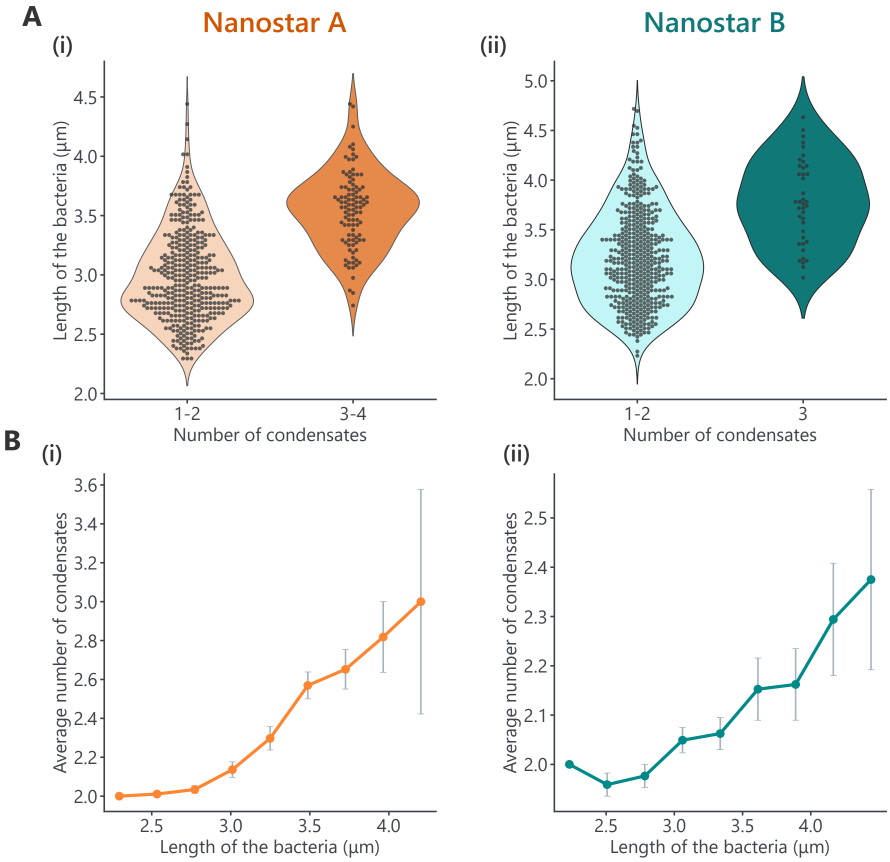


**Supplementary Figure 17**: **Correlation between the length of bacterial cells and the number of MLOs per cell.** (**A**) Violin plot of length of bacteria where cells have been grouped based on the number of identified MLOs per cell. (i) Cells expressing NS A. (ii) Cells expressing NS B. For both NSs the difference in length of the bacteria between cells with 1-2 MLOs and cells with 3+ MLOs is significant. For NS A, a Shapiro-Wilk test indicated that the group with 1-2 MLOs was not normally distributed ($p < 0.0001$). Levene's test showed unequal variances between groups ($p = 0.043$). Given these violations of parametric assumptions, a Kruskal-Wallis test was conducted, revealing a significant difference between the group with 1-2 MLOs ($n = 382$) and the group with 3-4 MLOs ($n = 102$), $H = 130.332$, $p < 0.0001$. For NS B, a Shapiro-Wilk test indicated that the group with 1-2 MLOs was not normally distributed ($p < 0.0001$). Levene's test showed no significant difference in variances between groups ($p = 0.410$). Given the non-normality in the group with 1-2 MLOs, a Kruskal-Wallis test was conducted, revealing a significant difference between the group with 1-2 MLOs ($n = 435$) and the group with 3 MLOs ($n = 40$), $H = 40.78$, $p < 0.0001$. (**B**) Average number of condensate versus cell length. Cells were binned into 10 groups by length of the cell and the average number of MLOs was calculated in each group. The error bars show the standard error of the mean. Cells with no MLOs were not included in the dataset.

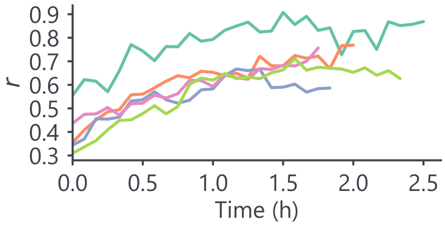


**Supplementary Figure 18**: **Ratio of condensate sizes within single cells expressing NS A over time.** The ratio $r$ was calculated as $r = I_{\text{cond}}^{\text{small}}/I_{\text{cond}}^{\text{large}}$, where $I_{\text{cond}}^{\text{small}}$ is the integrated fluorescence intensity of the smaller MLO and $I_{\text{cond}}^{\text{large}}$ is the integrated fluorescence intensity of the larger MLO. Each line represents one cell, tracked until the onset of cell division. The area of MLOs was selected manually using ImageJ. Images were acquired every 5 minutes.

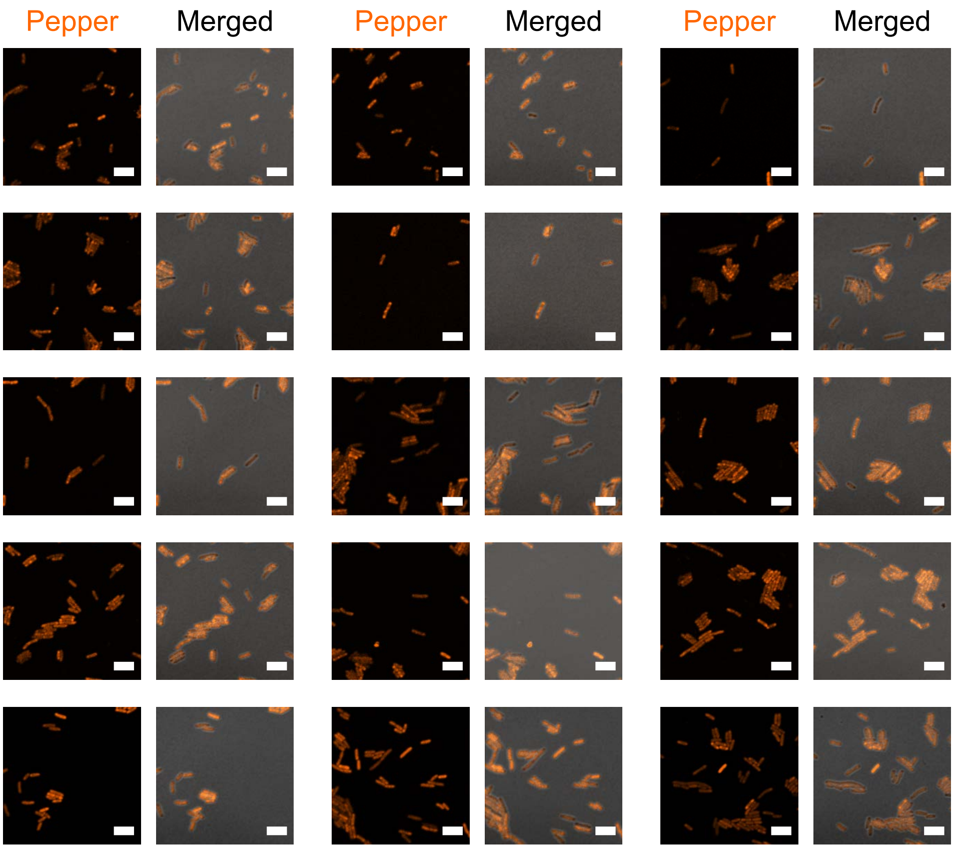


**Supplementary Figure 19**: **Additional micrographs of *E. coli* expressing NS $\bar{\text{A}}$.** Confocal micrographs (left) and merged confocal/bright-field images (right) of *E. coli* expressing NS $\bar{\text{A}}$, 1 h after IPTG induction. Scale bars, 5 µm. Micrographs were obtained from three separate cultures, each starting from an individual single colony.

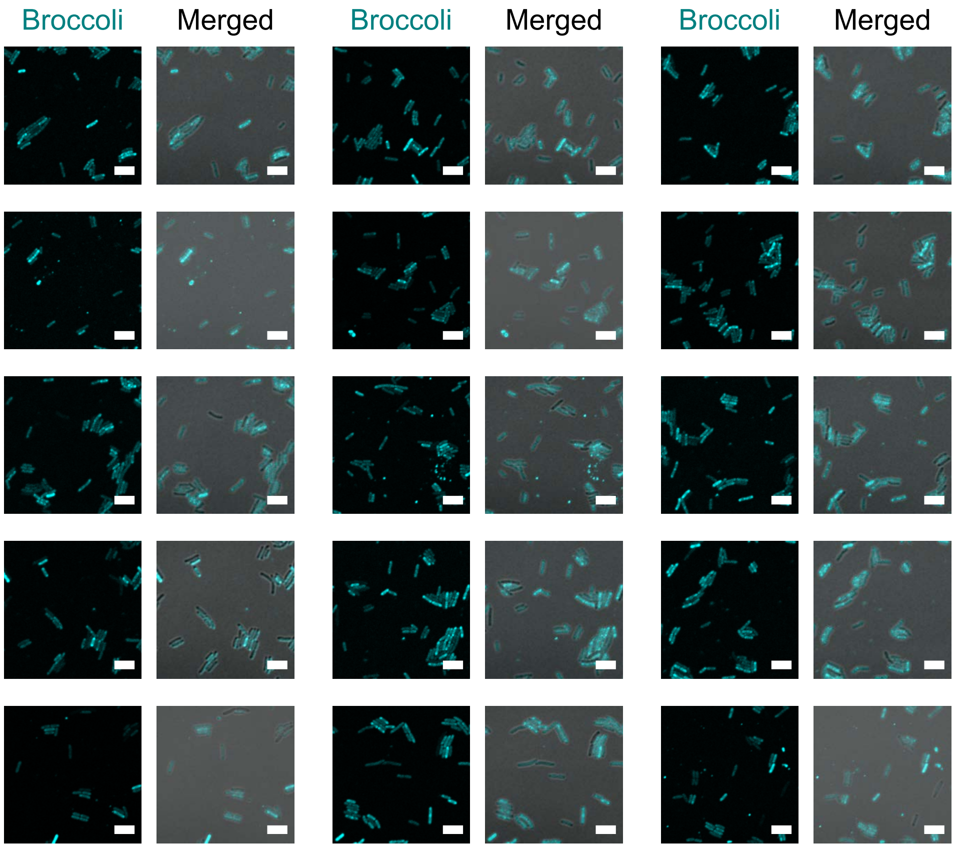


**Supplementary Figure 20**: **Additional micrographs of *E. coli* expressing NS $\bar{\mathbf{B}}$.** Confocal micrographs (left) and merged confocal/bright-field images (right) of *E. coli* expressing NS $\bar{\mathrm{B}}$, 1 h after IPTG induction. Scale bars, 5 µm. Micrographs were obtained from three separate cultures, each starting from an individual single colony.

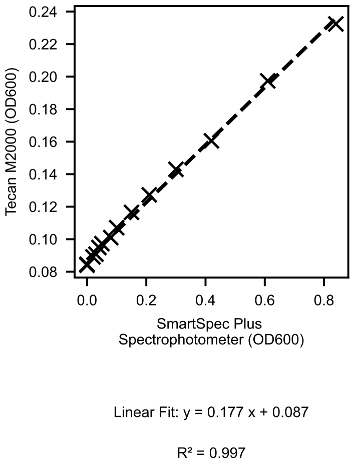


**Supplementary Figure 21**: **Calibration curve showing $OD_{600}$ values as measured on a Tecan M2000 plate reader and Bio-Rad SmartSpec Plus spectrophotometer.** Absolute OD values can vary between instruments due to difference in optical pathlength. In particular, platereaders like the Tecan M2000 typically have a shorter pathlength compared to cuvette-based spectrophotometers such as the Bio-Rad SmartSpec Plus. As a result, OD values from the plate reader are not directly comparable to standard spectrophotometer measurements. To account for this, we generated a calibration curve to convert plate reader measurements to equivalent spectrophotometer readings (Methods).

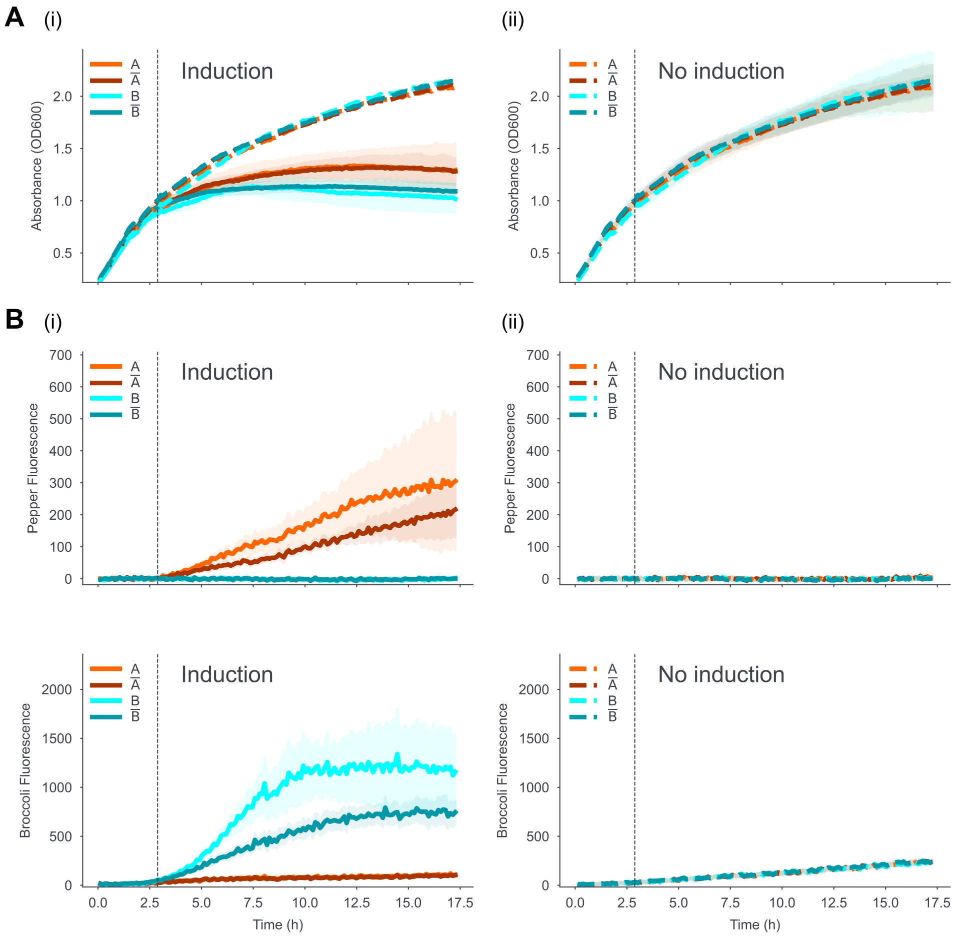


**Supplementary Figure 22**: **Growth and fluorescence curves of *E. coli* samples expressing a single NS.** (**A**) Optical Density (OD) curves for samples with (i) and without IPTG induction. In (i), data for uninduced samples are plotted as dashed lines for comparison. (**B**) Fluorescence curves collected in the Broccoli (NSs B and B̄) and Pepper (NSs A and Ā) for induced (i) and uninduced (ii) samples. For each condition, measurements were performed for three biological replicates from separate single colonies, with three technical replicates studied per biological replicate. Solid lines represent the mean values across all replicates, while the shaded regions indicate the standard deviation across all replicates. Data for induced samples (i) are also shown in Extended Data Fig. 7.

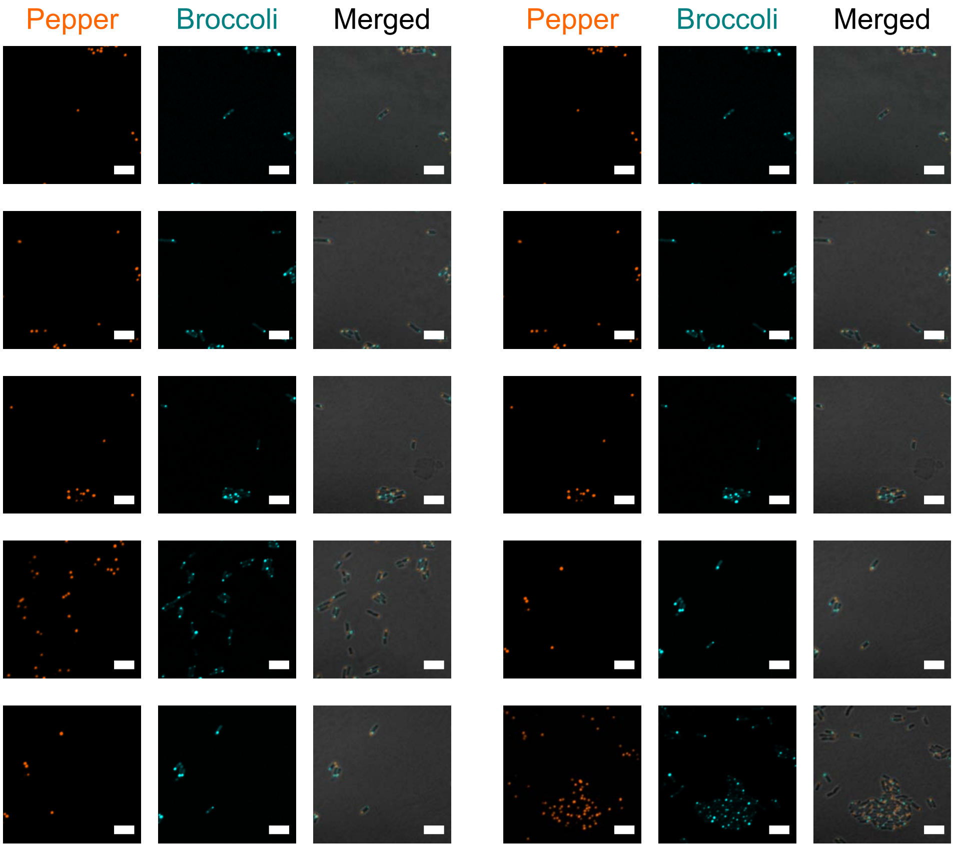


**Supplementary Figure 23**: **Additional micrographs of *E. coli* expressing NSs A+B.** Confocal micrographs recorded in the Pepper channel (NS A, left), the Broccoli channel (NS B, centre) and merging both fluorescence channels with bright-field images (right) of *E. coli* co-expressing NS A and NS B, 1 h after IPTG induction. Scale bars, 5 µm. Micrographs were obtained from three separate cultures, each starting from an individual single colony.

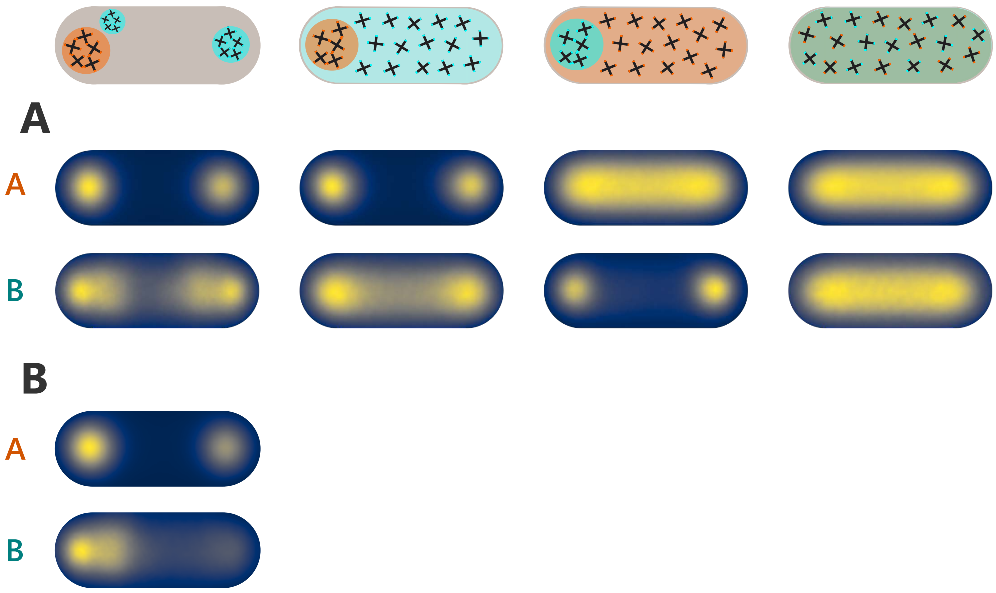


**Supplementary Figure 24**: **Average fluorescence intensity heatmaps of cells co-expressing two NSs** (**A**) Average heatmap generated from cells oriented randomly along the longitudinal axis. From left, cells co-expressing NSs A+B, A+$\bar{B}$, $\bar{A}$+B and $\bar{A}$+$\bar{B}$. (**B**) Average heatmap generated from cells co-expressing NSs A+B aligned such that the pole with highest Broccoli fluorescence is positioned on the left. In Fig. 2B(ii), maps were generated by aligning cells based on the pole with the highest Pepper fluorescence. These alternative maps are shown here for completeness.

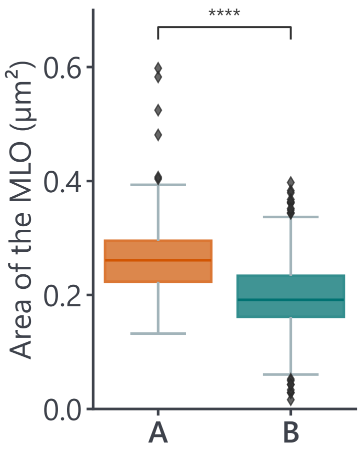


**Supplementary Figure 25**: **Comparison of the segmented cross-sectional area of the MLOs in *E. coli* co-expressing NSs A and B, 1 h after IPTG induction.** A Shapiro-Wilk test indicated that the distribution of areas of both NSs A and B MLOs was not normally distributed ($p < 0.0001$). Levene's test showed no significant difference in variances between groups ($p = 0.308$). Given the non-normality of both groups, a Kruskal-Wallis test was conducted, revealing a significant difference in the apparent cross-sectional area of NS A ($n = 399$) and NS B ($n = 399$), $H = 208.671$, $p < 0.0001$.

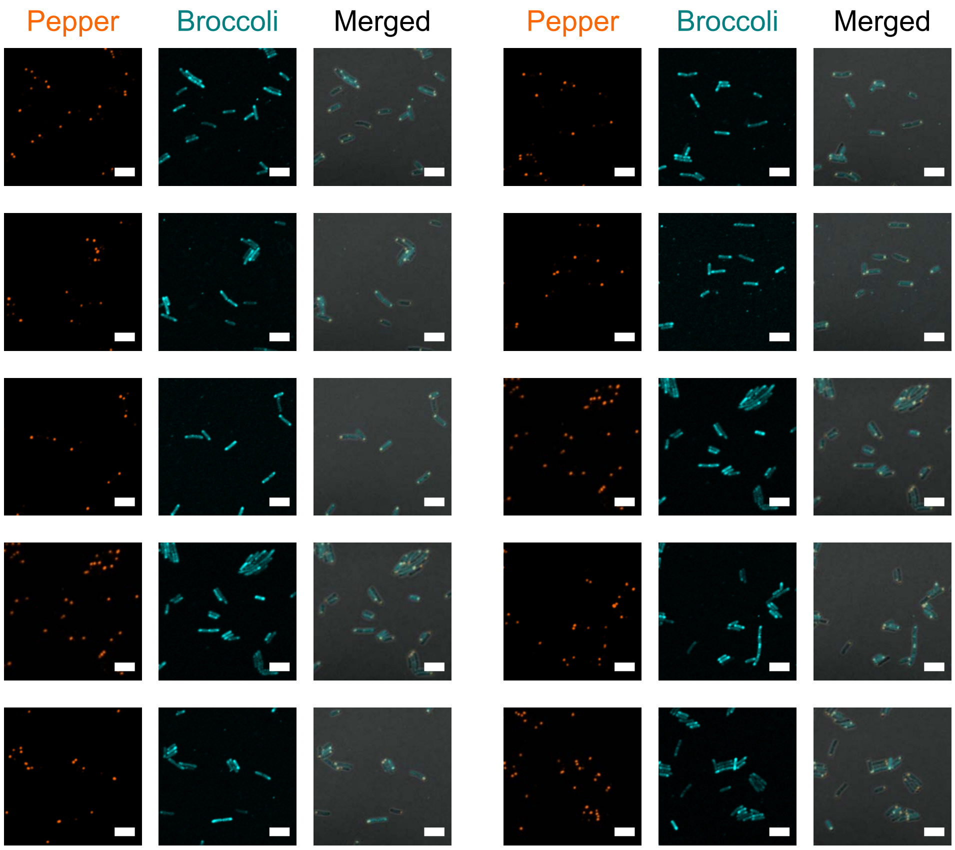


**Supplementary Figure 26**: **Additional micrographs of *E. coli* expressing NSs A+$\bar{\text{B}}$.** Confocal micrographs recorded in the Pepper channel (NS A, left), the Broccoli channel (NS $\bar{\text{B}}$, centre) and merging both fluorescence channels with bright-field images (right) of *E. coli* co-expressing NS A and NS $\bar{\text{B}}$, 1 h after IPTG induction. Scale bars, 5 µm. Micrographs were obtained from three separate cultures, each starting from an individual single colony.

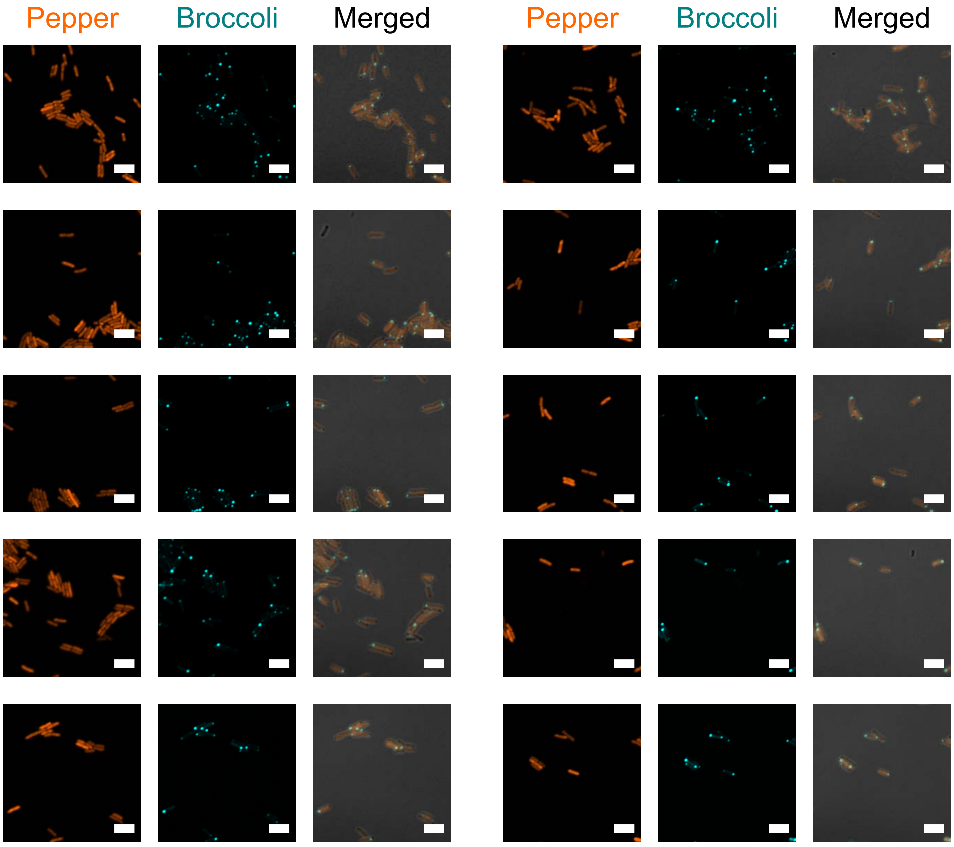


**Supplementary Figure 27**: **Additional micrographs of *E. coli* expressing NSs $\bar{A}$+B.** Confocal micrographs recorded in the Pepper channel (NS $\bar{A}$, left), the Broccoli channel (NS B, centre) and merging both fluorescence channels with bright-field images (right) of *E. coli* co-expressing NS $\bar{A}$ and NS B, 1 h after IPTG induction. Scale bars, 5 µm. Micrographs were obtained from three separate cultures, each starting from an individual single colony.

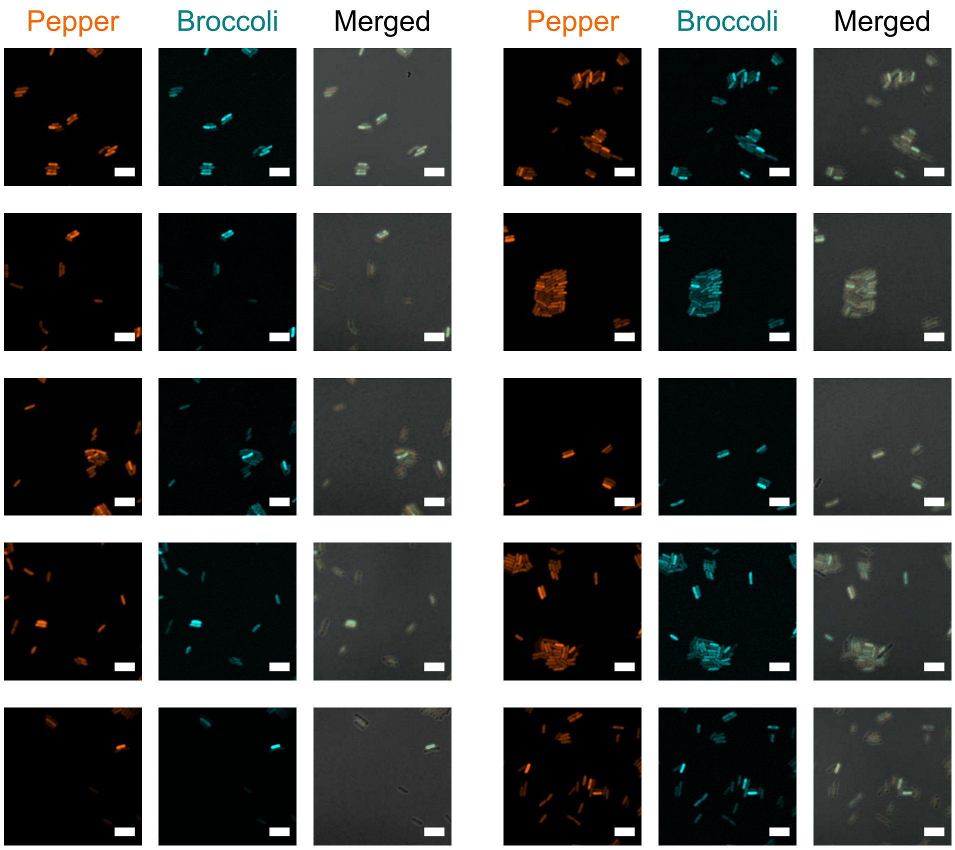


**Supplementary Figure 28**: **Additional micrographs of *E. coli* expressing NSs $\bar{\text{A}}$+$\bar{\text{B}}$.** Confocal micrographs recorded in the Pepper channel (NS $\bar{\text{A}}$, left), the Broccoli channel (NS $\bar{\text{B}}$, centre) and merging both fluorescence channels with bright-field images (right) of *E. coli* co-expressing NS $\bar{\text{A}}$ and NS $\bar{\text{B}}$, 1 h after IPTG induction. Scale bars, 5 µm. Micrographs were obtained from three separate cultures, each starting from an individual single colony.

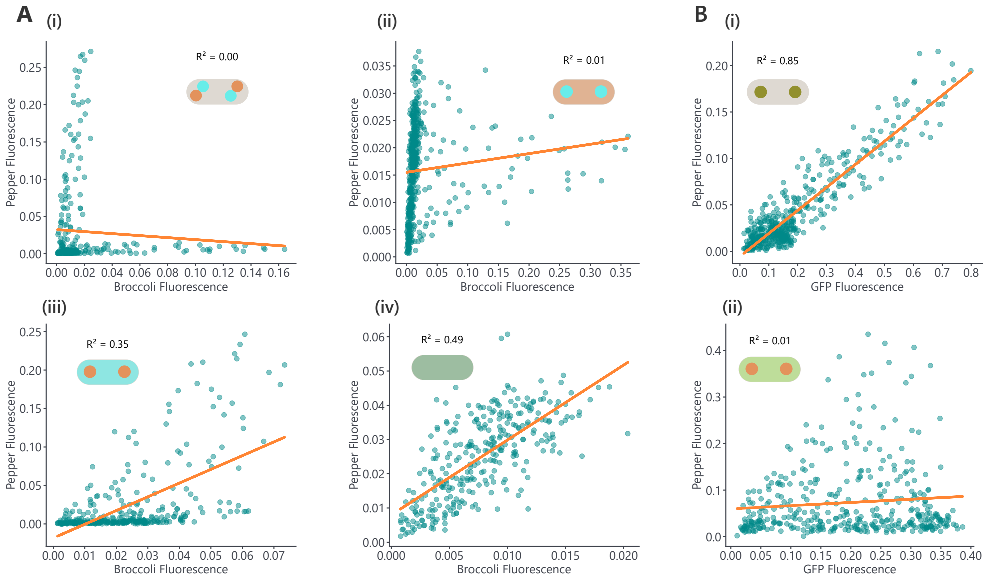


**Supplementary Figure 29**: **Examples of correlation plots between fluorescent channels used to calculate coefficients of determination ($R^2$).** (**A**) Correlation between pixel fluorescent intensity in the Broccoli and Pepper channels for single cells. (i) Example of a cell expressing NSs A and B. High intensity in one channel occurs when the other channel shows no fluorescence as the orthogonality of kissing-loop interactions produces de-mixed A and B MLOs. The resulting scatter plot shows an "L" shape, producing low $R^2$. (ii) Example of a cell expressing NSs $\bar{\text{A}}$ and B. The soluble NS $\bar{\text{A}}$ produces a uniform background in the Pepper channel, while B MLO results in a sub-set of brighter pixels, leading to low correlation. (iii) Example of a cell expressing NS A and $\bar{\text{B}}$. The behaviour is analogous to panel (iii), but correlation is higher due to possible co-assembly of $\bar{\text{B}}$ on A MLOs and fluorescence bleed-through (see main text for discussion). (iv) Example of a cell expressing NSs $\bar{\text{A}}$ and $\bar{\text{B}}$. Fluorescent signal from both soluble NSs is uniformly distributed across the cell, leading to medium correlation. (**B**) Correlation between pixel fluorescent intensity in the GFP and Pepper channels. (i) Example of a cell expressing GFP-binding NSs $\text{A}_{\text{AP3}}$ and GFP. Co-localisation of the two signals yields high correlation. (ii) Example of a cell expressing non-GFP-binding NSs A and GFP. The uniform fluorescent background produced by the proteins is poorly correlated with the localised signal from the NSs. In all plots, data are shown for a single, representative cell.

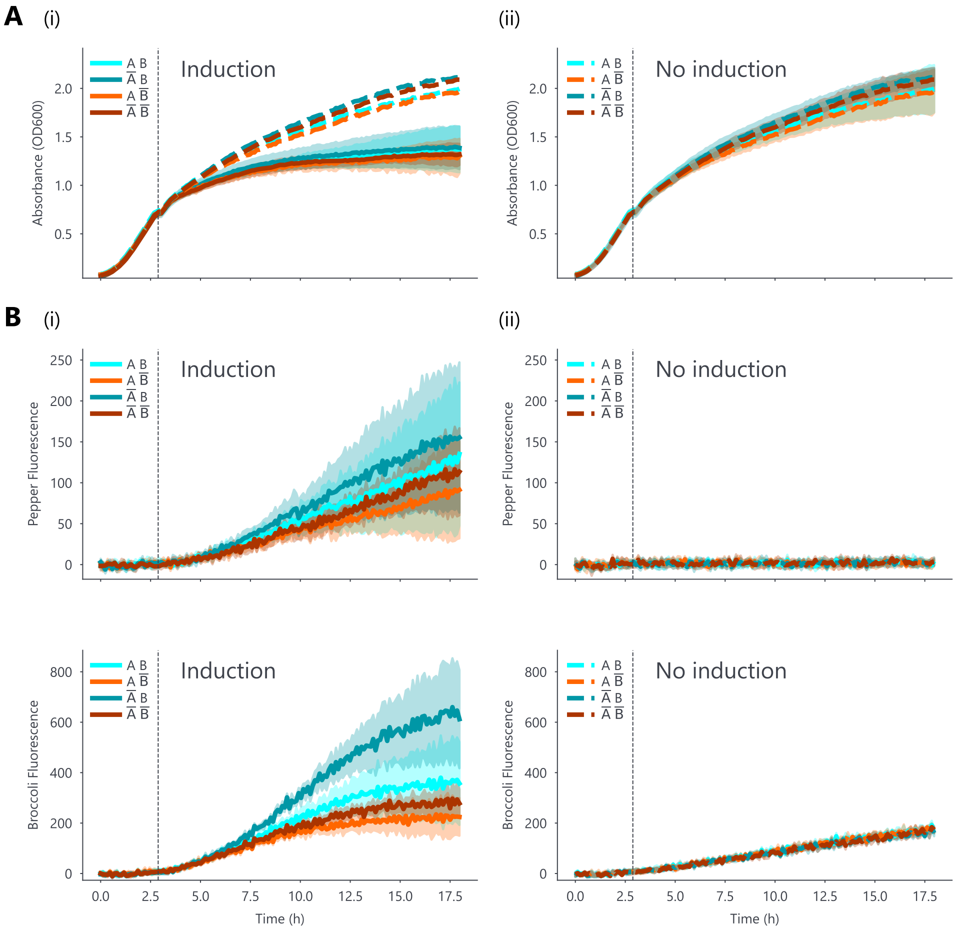


**Supplementary Figure 30: Growth and fluorescence curves of induced *E. coli* samples expressing two NSs.** (**A**) Optical Density (OD) curves for samples with (i) and without (ii) IPTG induction. In (i), data for uninduced samples are plotted as dashed lines for comparison. (**B**) Fluorescence curves collected in the Broccoli (NSs B and $\bar{B}$) and Pepper (NSs A and $\bar{A}$) for induced (i) and uninduced (ii) samples. For each condition, measurements were performed for three biological replicates from separate single colonies, with three technical replicates studied per biological replicate. Solid lines represent the mean values across all replicates, while the shaded regions indicate the standard deviation across all replicates. Data for induced samples (i) are also shown in Extended Data Fig. 7.

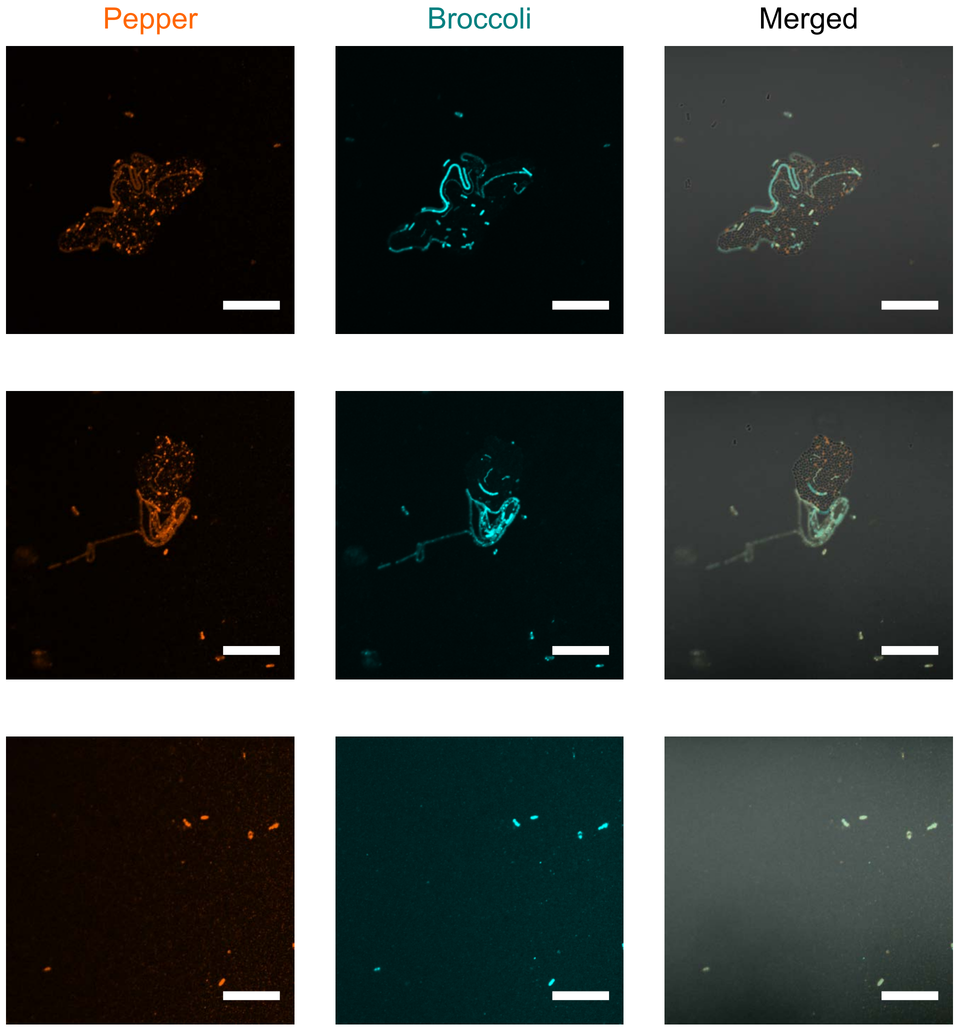


**Supplementary Figure 31**: **Micrographs of *E. coli* expressing NSs A+B after 7 days.** Confocal micrographs recorded in the Pepper channel (nanostar A, left), the Broccoli channel (nanostar B, centre) and merging both fluorescence channels with bright-field images (right) of *E. coli* co-expressing nanostars A and B. Images were collected 7 days after IPTG induction, during which cells were kept at room temperature. Many cells retain signal from both MLOs. Scale bars, 20 µm.

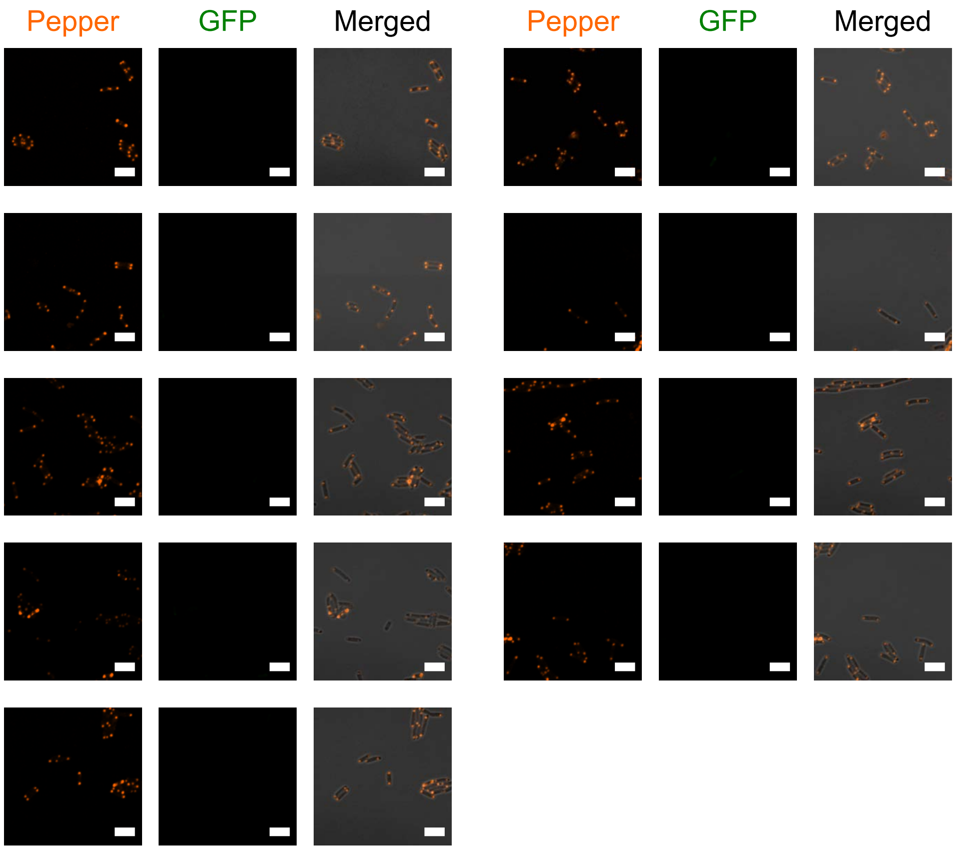


**Supplementary Figure 32**: **Additional micrographs of *E. coli* expressing NS $\mathbf{A_{AP3}}$.** Confocal micrographs recorded in the Pepper channel (NS $A_{AP3}$, left), the GFP channel, and merging both fluorescence channels with bright-field images (right) of *E. coli* expressing NS $A_{AP3}$ only (no GFP), 1 h after IPTG induction. Scale bars, 5 µm. Micrographs were obtained from three separate cultures, each starting from an individual single colony.

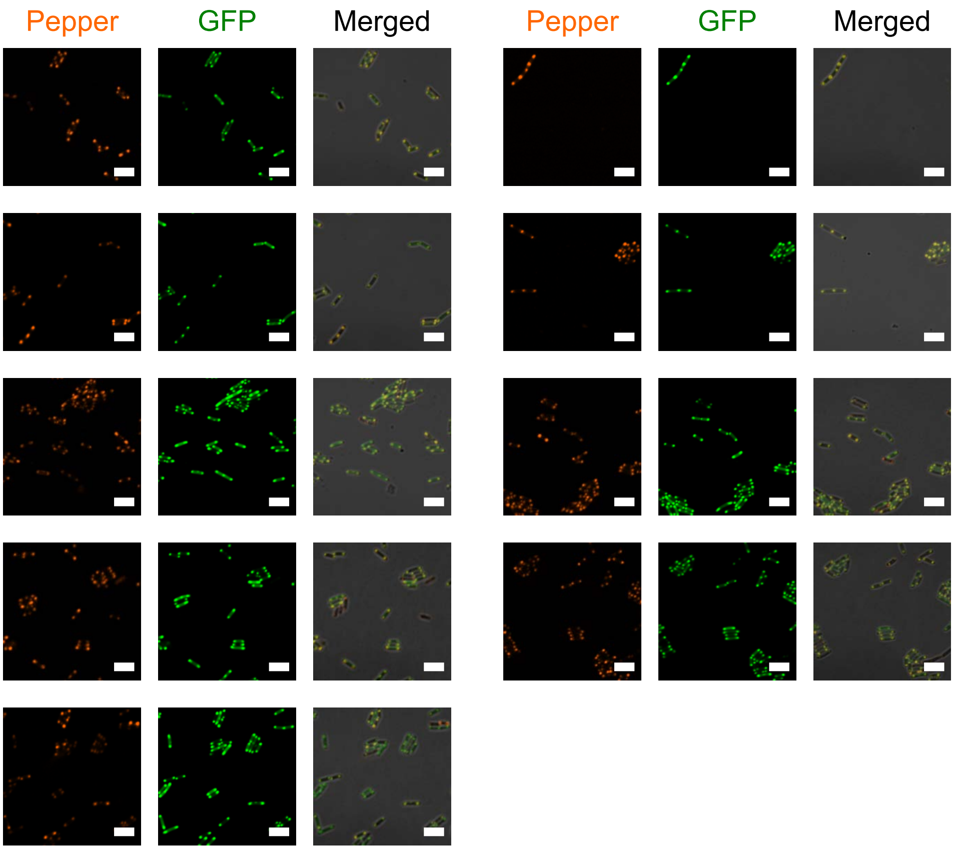


**Supplementary Figure 33**: **Additional micrographs of *E. coli* co-expressing NS $A_{AP3}$ and GFP.** Confocal micrographs recorded in the Pepper channel (NS $A_{AP3}$, left), the GFP channel, and merging both fluorescence channels with bright-field images (right) of *E. coli* co-expressing NS $A_{AP3}$ and GFP, 1 h after IPTG induction. Scale bars, 5 µm. Micrographs were obtained from three separate cultures, each starting from an individual single colony.

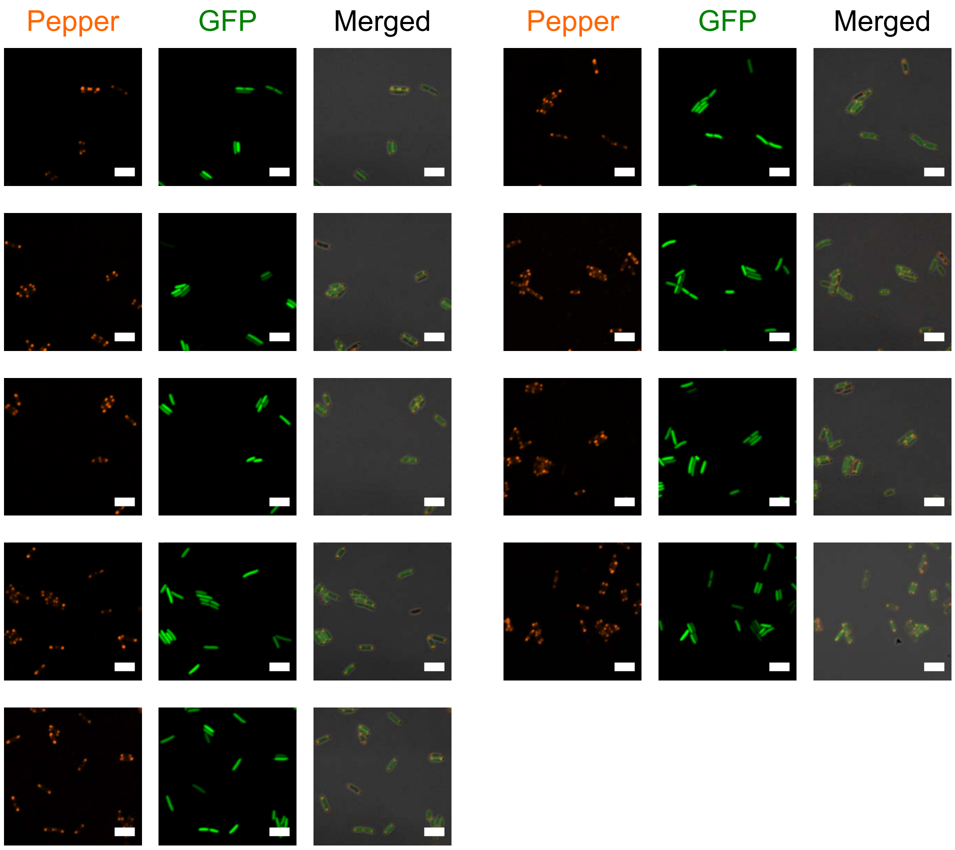


**Supplementary Figure 34**: **Additional micrographs of *E. coli* co-expressing NS A and GFP.** Confocal micrographs recorded in the Pepper channel (NS A, left), the GFP channel, and merging both fluorescence channels with bright-field images (right) of *E. coli* co-expressing NS A and GFP, 1 h after IPTG induction. Scale bars, 5 µm. Micrographs were obtained from three separate cultures, each starting from an individual single colony.

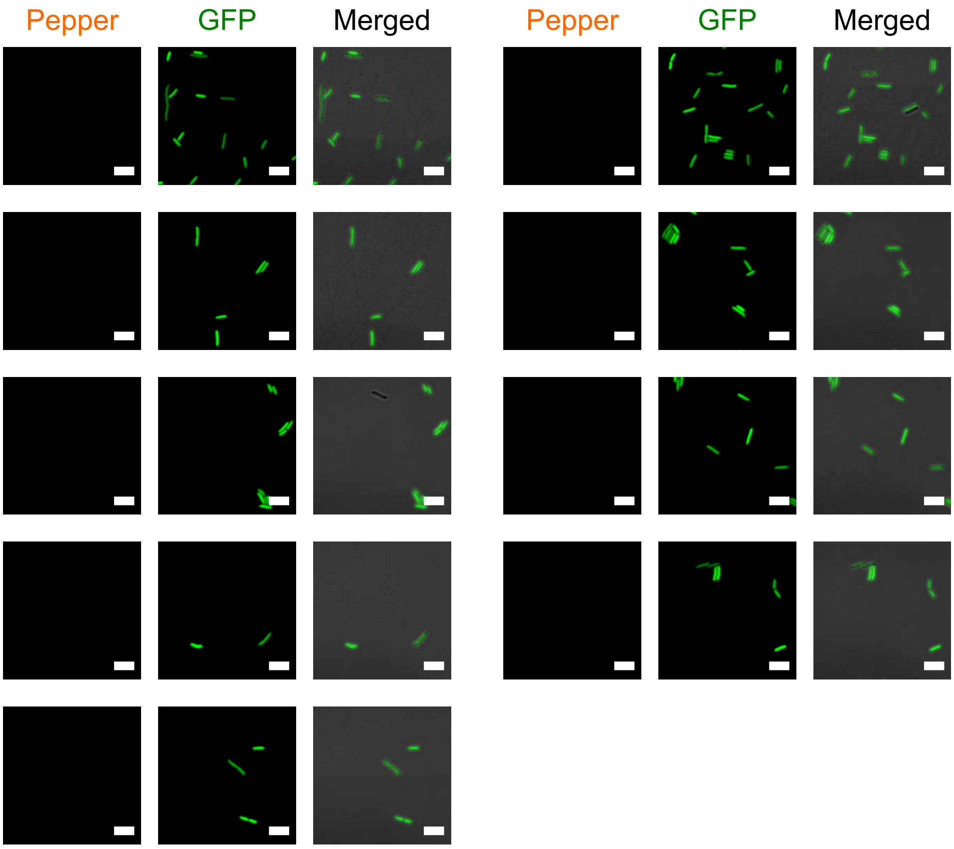


**Supplementary Figure 35**: **Additional micrographs of *E. coli* expressing GFP.** Confocal micrographs recorded in the Pepper channel (left), the GFP channel, and merging both fluorescence channels with bright-field images (right) of *E. coli* expressing GFP only (no NSs), 1 h after IPTG induction. Scale bars, 5 µm. Micrographs were obtained from three separate cultures, each starting from an individual single colony.

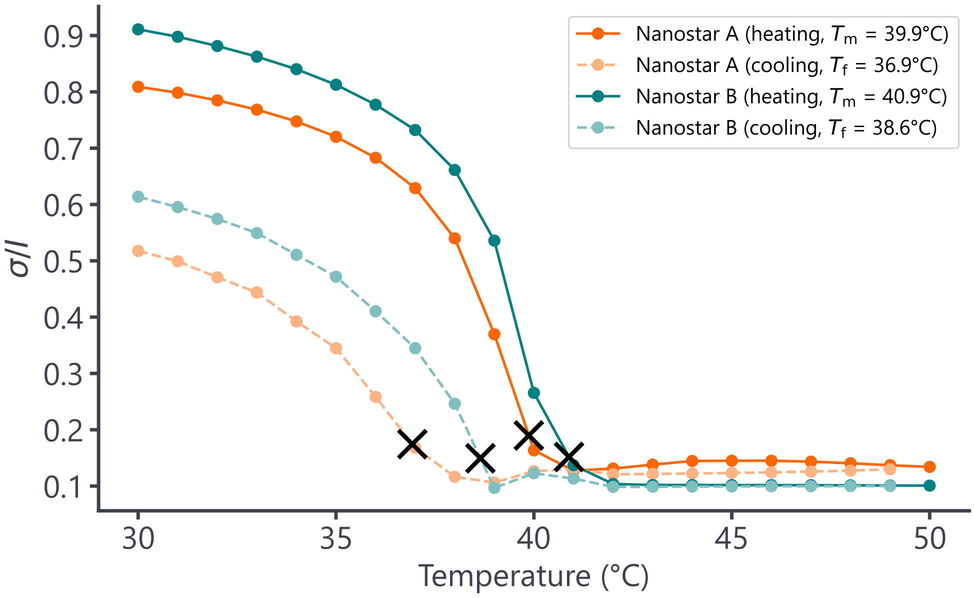


**Supplementary Figure 36**: **Thermal transition curves of A and B condensates expressed in vitro.** The thermal transition curves of NS A condensates are shown in orange, and those of NS B condensates in cyan. Solid curves corresponding to the heating ramp were used to estimate the melting temperatures ($T_m$), while dashed curves corresponding to the cooling ramp were used to estimate the condensate-formation temperatures ($T_f$). Data show the ratio between the standard deviation of the pixel values ($\sigma$) and the mean pixel intensity (I) of the epifluorescence micrographs. The quantity $\sigma/I$, plotted against $T$ displays a sharp transition when condensates melt or form, which can be used to estimate $T_m$ and $T_f$. These were computed from the intersections of the relevant curve with a threshold value in $\sigma/I$, defined as the high-temperature plateau of the data (mean computed for $T > 41\,^\circ C$) augmented by 0.05 units (Methods). For both A and B, $T_m$ values (39.9 °C and 40.9 °C, respectively) are slightly higher than their corresponding $T_f$ values (36.9 °C and 38.6 °C, respectively), reflecting the slower timescales of condensate assembly compared to condensate melting. Notably, the calculated $T_m$ and $T_f$ values are substantially lower than those observed in vivo, likely due to a combination of different ionic conditions and molecular crowding. Samples were initially incubated at 30 °C for 15 hours, after which the temperature was gradually changed from 30 °C to 50 °C and back to 30 °C, with 1 °C steps introduced every 5 minutes.

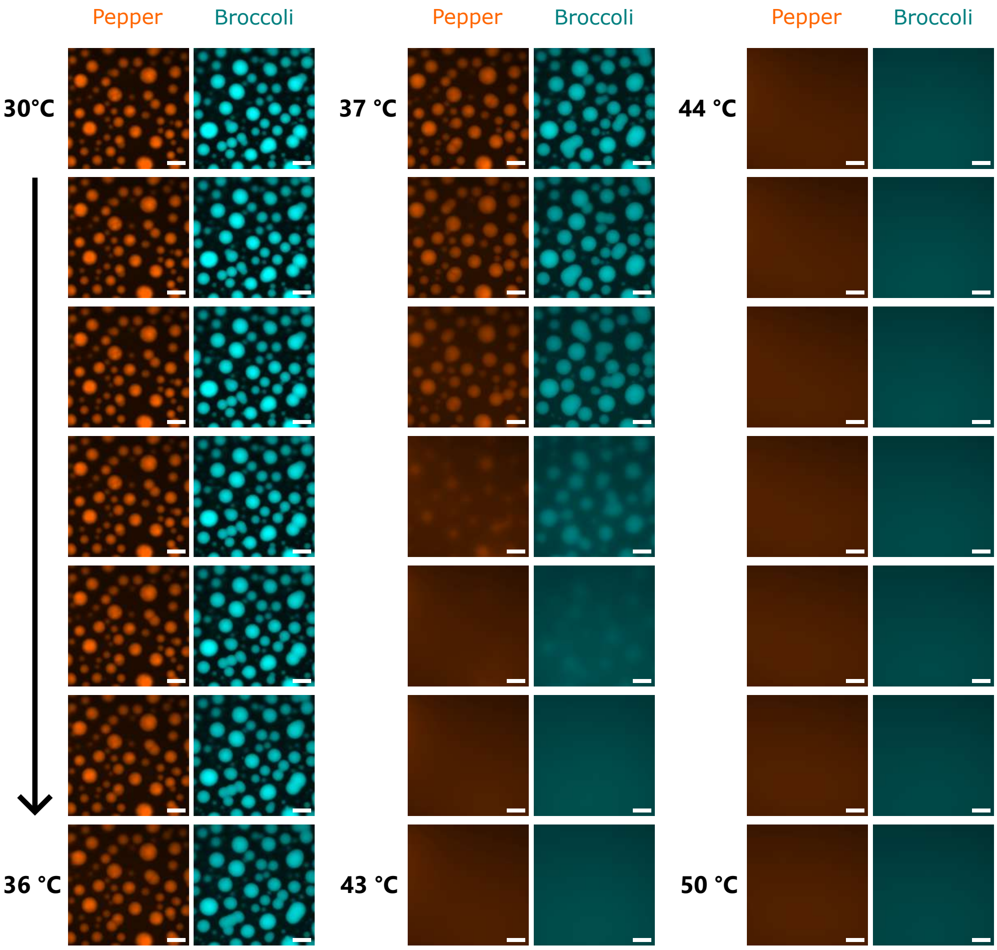


**Supplementary Figure 37**: **Epifluorescence micrographs of NS A (left) and NS B (right) condensate melting upon increasing temperature.** After the initial 15 hour incubation at 30 °C, the temperature was gradually increased from 30 °C to 50 °C in 1 °C steps introduced every 5 minutes, leading to NS condensate melting. Scale bars, 100 µm.

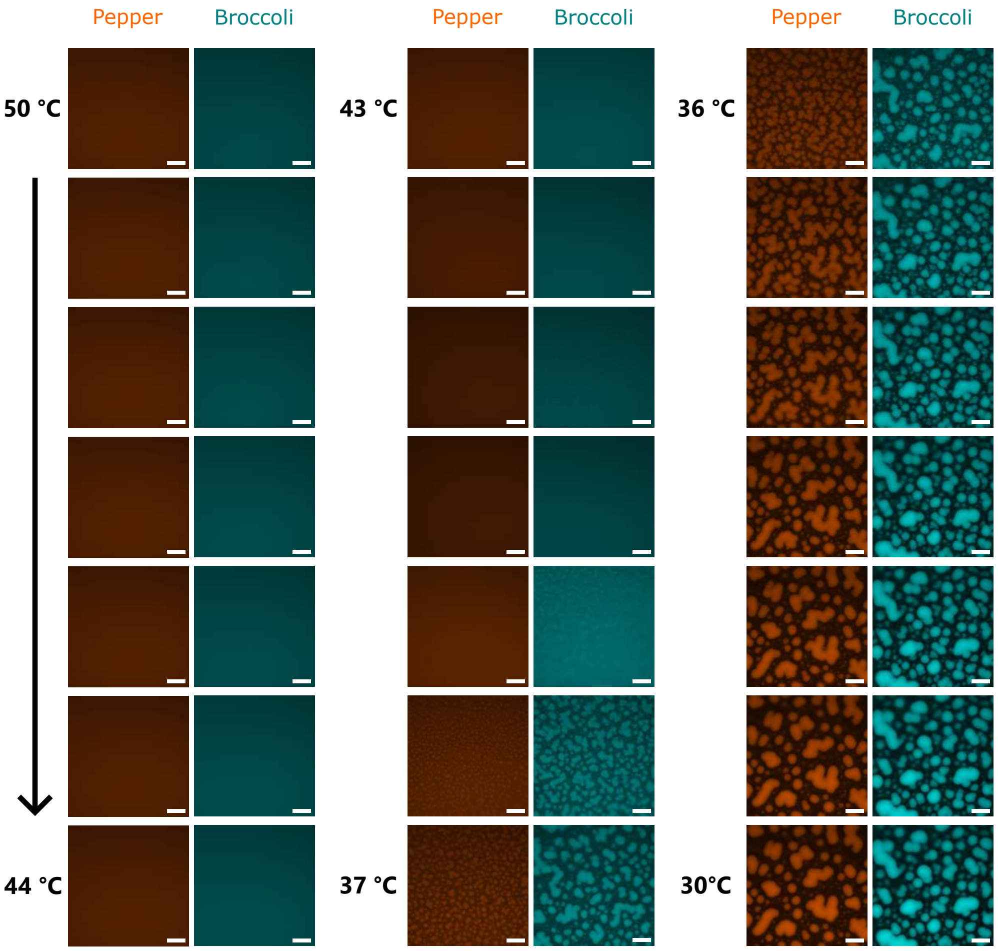


**Supplementary Figure 38**: **Epifluorescence micrographs of A (left) and B (right) condensate formation upon cooling.** The temperature was gradually decreased from 50 °C to 30 °C in 1 °C steps introduced every 5 minutes, leading to NS condensate-formation. Scale bars, 100 µm.

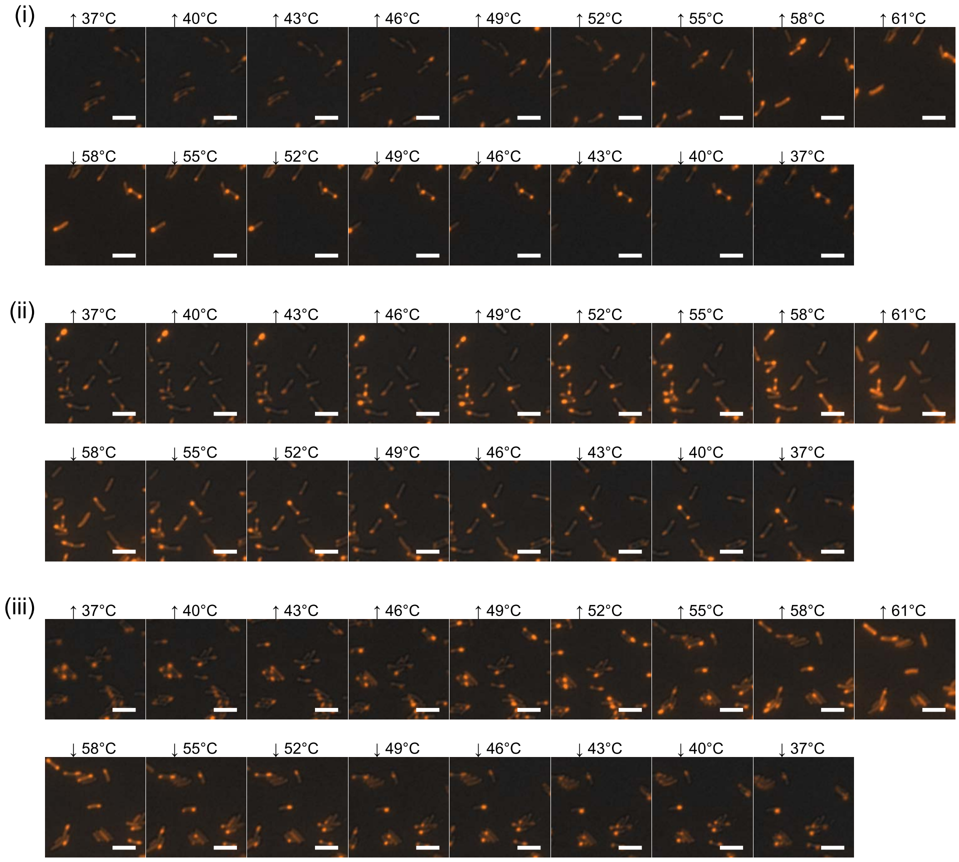


**Supplementary Figure 39**: **Additional micrographs of *E. coli* expressing NS A at different temperatures.** Merged epifluorescence (Pepper) and bright-field micrographs of *E. coli* expressing NS A at different temperatures. (i)-(iii) are micrographs from separate experiments, each starting from a single colony. Scale bars, 5 µm. Upward arrows indicate samples being heated, while downward arrows indicate samples being cooled. Micrographs were obtained from three separate cultures, each starting from an individual single colony.

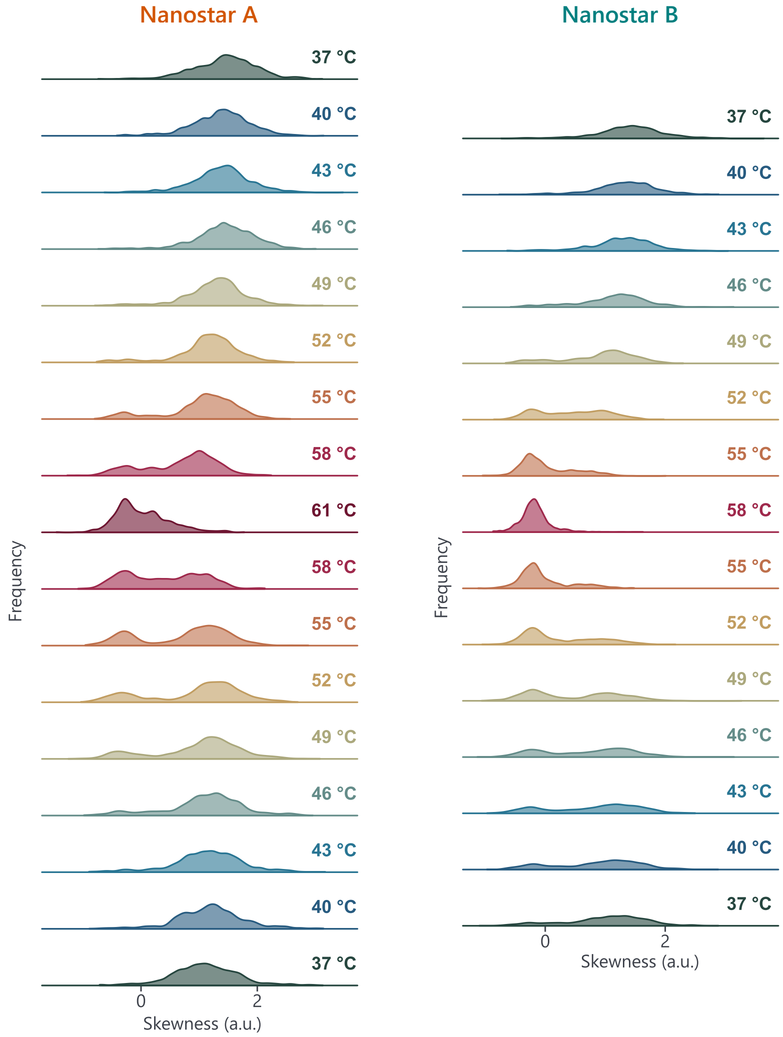


**Supplementary Figure 40**: **Cell population distribution of the skewness of the pixel-intensity distribution recorded within individual cells as a function of temperature.** The data is from three biological replicates conducted on *E. coli* expressing NS A (left) or NS B (right). The sample was heated from 37 °C ($n_A$ = 820, $n_B$ = 854) through 40 °C ($n_A$ = 819, $n_B$ = 866), 43 °C ($n_A$ = 858, $n_B$ = 862), 46 °C ($n_A$ = 872, $n_B$ = 886), 49 °C ($n_A$ = 854, $n_B$ = 827), 52 °C ($n_A$ = 801, $n_B$ = 849), 55 °C ($n_A$ = 840, $n_B$ = 846), 58 °C ($n_A$ = 814, $n_B$ = 873) up to 61 °C ($n_A$ = 795) for NS A and then cooled back down through 58 °C ($n_A$ = 878), 55 °C ($n_A$ = 903, $n_B$ = 889), 52 °C ($n_A$ = 896, $n_B$ = 877), 49 °C ($n_A$ = 885, $n_B$ = 914), 46 °C ($n_A$ = 874, $n_B$ = 877), 43 °C ($n_A$ = 896, $n_B$ = 895), 40 °C ($n_A$ = 879, $n_B$ = 907) to 37 °C ($n_A$ = 901, $n_B$ = 871), where $n$ is the number of segmented bacteria. Changes in cell number during heating and cooling are due to temperature-induced shifts in the field of view and ongoing cell division.

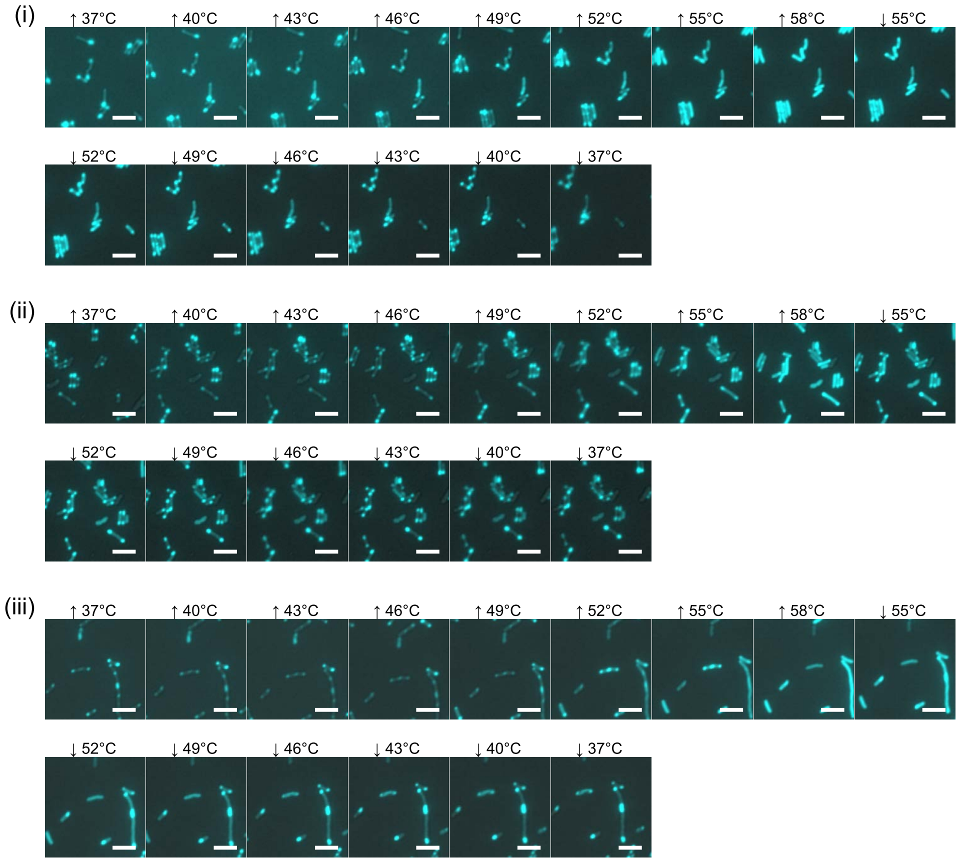


**Supplementary Figure 41**: **Additional micrographs of *E. coli* expressing NS B at different temperatures.** Merged epifluorescence (Broccoli) and bright-field micrographs of *E. coli* expressing NS B at different temperatures. (i)-(iii) are micrographs from separate experiments, each starting from a single colony. Upward arrows indicate samples being heated, while downward arrows indicate samples being cooled. Scale bars, 5 µm.

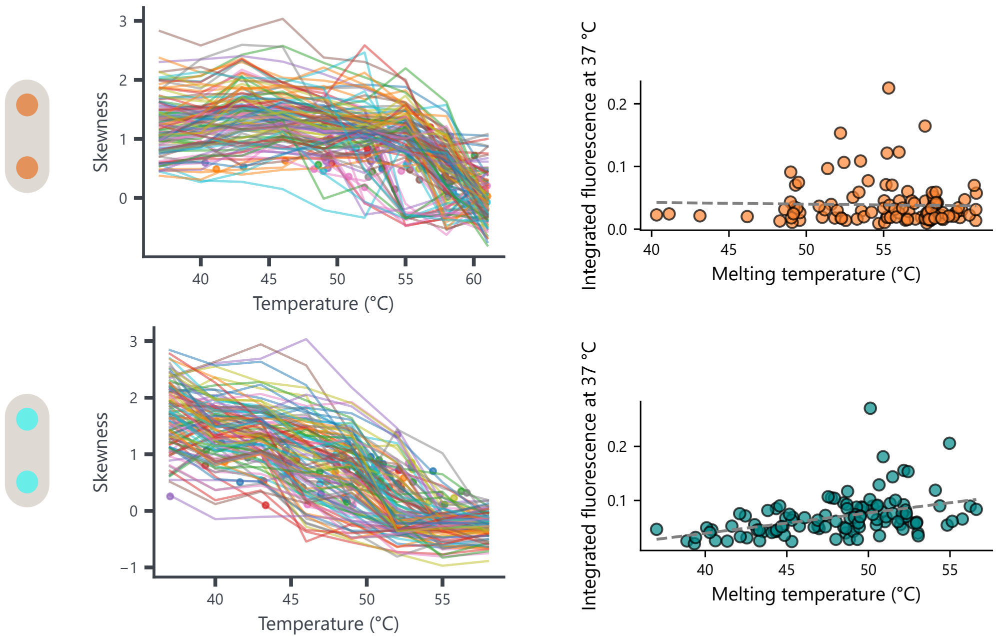


**Supplementary Figure 42**: **Analysis of the relationship between fluorescence intensity and melting temperature.** Each row contains data from one biological replicate of *E. coli* expressing A (top) or NS B (bottom). The left panels show single-cell skewness as a function of temperature, with the melting temperature determined by fitting a sigmoid curve (indicated by dots). The right panels display the relationship between the melting temperature and the integrated cellular fluorescence at the starting temperature of 37 °C. Linear fits were used to calculate the coefficients of determination ($R^2$) for NS A ($R^2 = 0.00$) and NS B ($R^2 = 0.18$). We note that, for NS A, skewness-temperature curves did not reach their high-temperature plateau at the maximum temperature we probed (61 °C), making the sigmoid fit imprecise. Furthermore, for both NS types, the reliability of the sigmoid fit is limited by the number of temperatures probed and the intrinsic noise of the measurements.

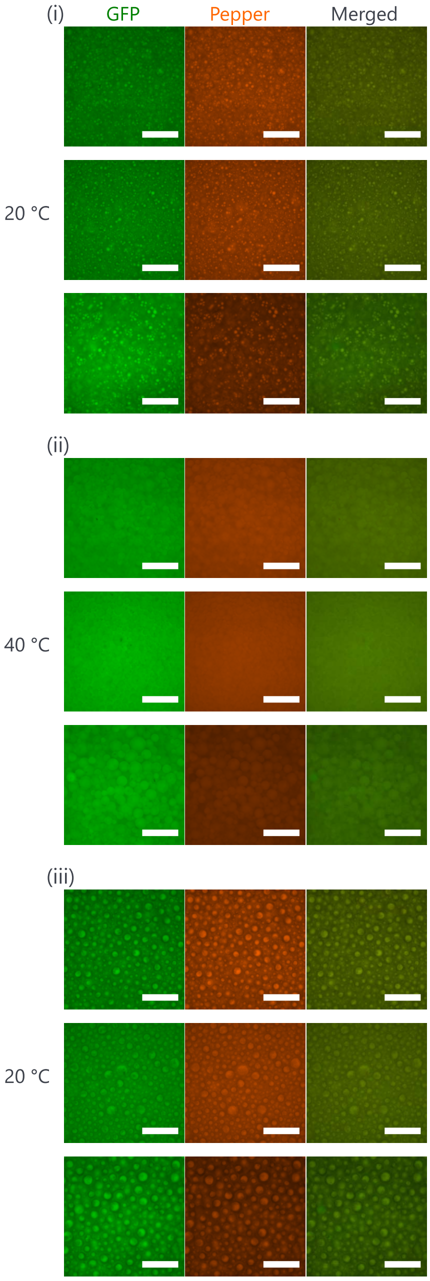


**Supplementary Figure 43**: **Epifluorescence micrographs of in vitro expression of NS $\mathbf{A_{AP3}}$, incubated with GFP in water-in-oil droplets.** Images were taken at (i) 20 °C, (ii) 40 °C, and (iii) back to 20 °C. Scale bars, 200 µm.

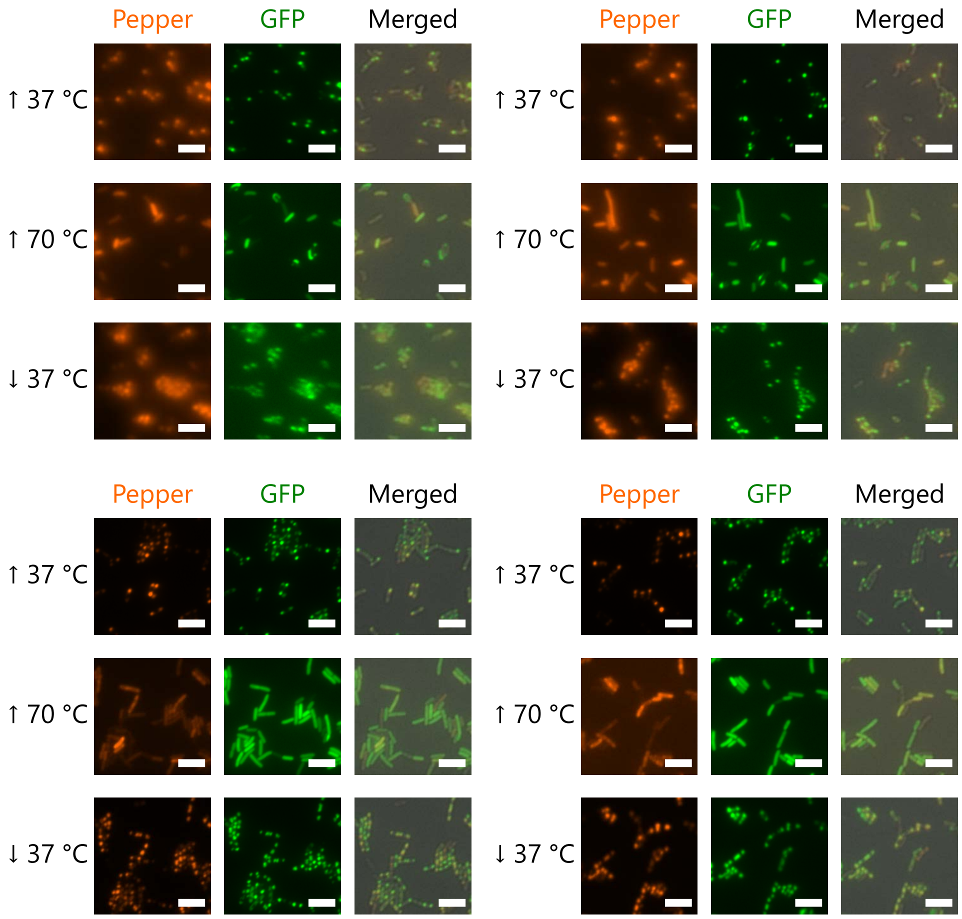


**Supplementary Figure 44**: **Additional micrographs of *E. coli* expressing NS $A_{AP3}$ and GFP at different temperatures.** Additional fluorescence micrographs, recorded in the Pepper channel (NS $A_{AP3}$, left), GFP channel (centre) and merging both fluorescent channels with bright-field images (right) of *E. coli* co-expressing NS $A_{AP3}$ and GFP. Images are shown at 37 °C, 70 °C, and after returning to 37 °C. Scale bars, 20 µm. Upward arrows indicate samples being heated, while downward arrows indicate samples being cooled. The micrographs are obtained from two separate cultures, each starting from a single colony.

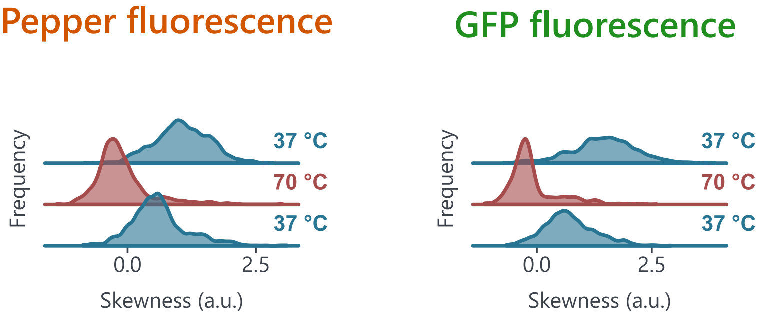


**Supplementary Figure 45**: **Cell population distribution of the skewness of the pixel-intensity distribution recorded within individual cells as a function of temperature.** Data are shown for bacteria co-expressing NS $A_{AP3}$ and GFP as a function of temperature. The samples were heated from 37 °C ($n = 831$) to 70 °C ($n = 774$) and then cooled back down to 37 °C ($n = 804$), where $n$ is the number of segmented bacteria.

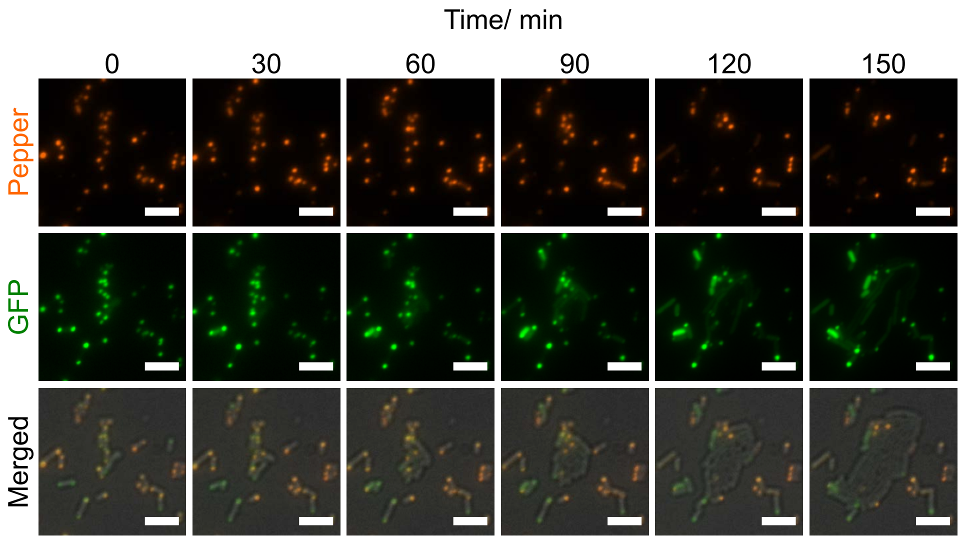


**Supplementary Figure 46**: **Timelapse micrographs of *E. coli* expressing NS $A_{AP3}$ and GFP after heating to 70 °C.** Timelapse confocal micrographs in the Pepper channel (NS $A_{AP3}$, top), the GFP channel (middle) and merging both fluorescent channels and bright-field images (bottom) of *E. coli* co-expressing NS $A_{AP3}$ and GFP, after heating to 70 °C. 10% of the cells retained the ability to grow and divide after heating to 70 °C for 2 min. Scale bars, 5 µm.

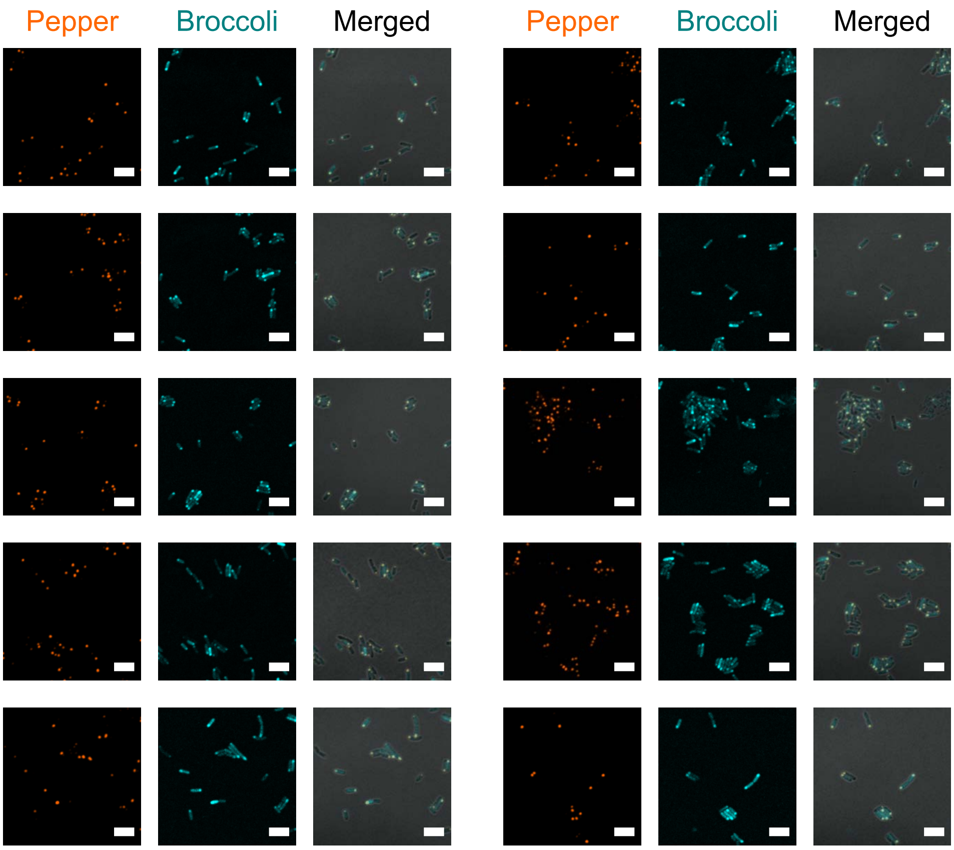


**Supplementary Figure 47**: **Additional micrographs of *E. coli* expressing NSs A+$\bar{\text{B}}$ before optimisation of imaging settings.** Additional confocal micrographs of *E. coli* expressing NSs A and $\bar{\text{B}}$, before optimizing imaging settings to minimise Pepper fluorescence in the Broccoli channel, 1 h after IPTG induction. Compare with optimised imaging settings in Supplementary Fig. 26. Scale bars, 5 µm. Micrographs were obtained from three separate cultures, each starting from an individual single colony.

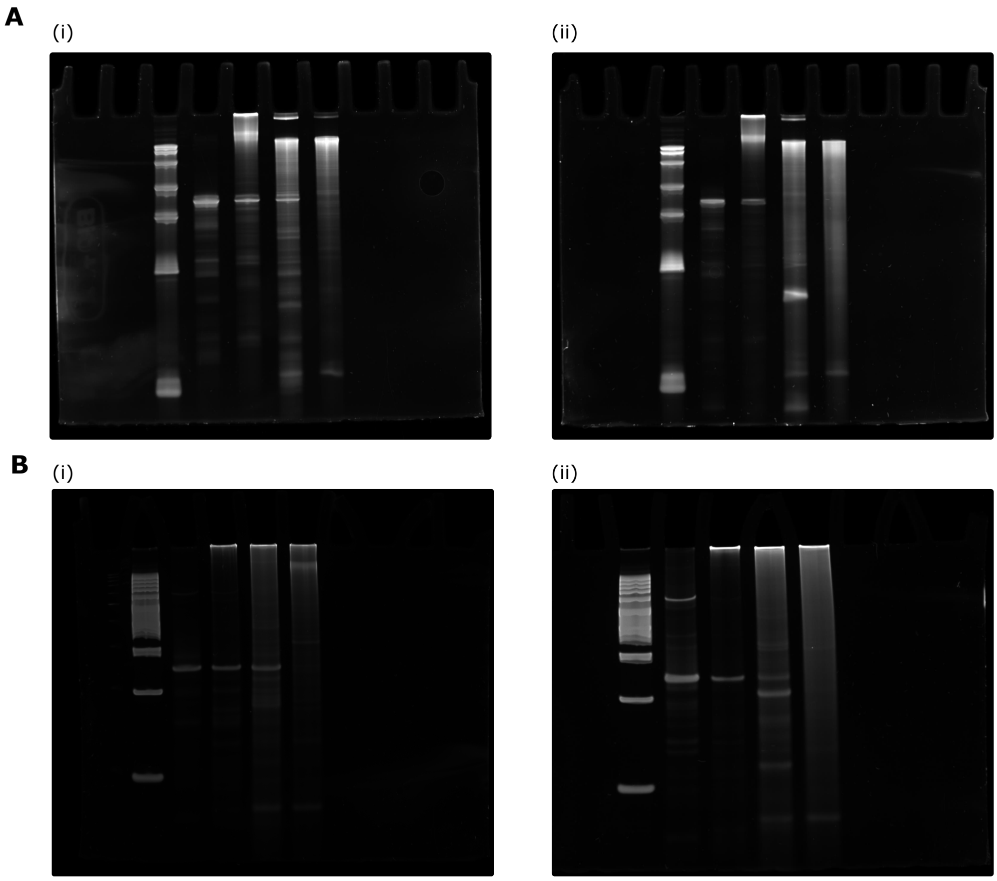


**Supplementary Figure 48**: **Full and uncropped PAGE gels from Extended Figs. 8 and 9.** (**A**) Denaturing PAGE of (i) NS A samples and (ii) NS B samples. (**B**) Native PAGE of (i) NS A samples and (ii) NS B samples.

| Nanostars | RNA Sequences (5'-3') |
|---|---|
| A | GCACAGUGCUAUGAGUGUGCACGGGAUCCGAUCCCCAAUCGUGGCGUGUCGGCCUGCAUCGC<br>GAAAGCAGGCACUGGCGCCGGGAUCGGAUCCUGUGCUGCACAUUAGAGUCGCUGUAUGACCC<br>AUCGCGAAAGGGUCGUACAGCGGCUCUAGUGUGCUCGCGUGCCUCAGAGGACCUGUCACCAU<br>CGCGAAAGGUGAUAGGUCCUUUGAGGUACGCGUCACUCGUAGCAUUGUGCCUGUCUCCAUCG<br>CGAAAGGAGAUAG |
| $\bar{A}$ | GCACAGUGCUAUGAGUGUGCACGGGAUCCGAUCCCCAAUCGUGGCGUGUCGGCCUGCCGCCA<br>ACGAGCAGGCACUGGCGCCGGGAUCGGAUCCUGUGCUGCACAUUAGAGUCGCUGUAUGACCC<br>AUAGCACAAGGGUCGUACAGCGGCUCUAGUGUGCUCGCGUGCCUCAGAGGACCUGUCACCAU<br>GGCGAAAGGUGAUAGGUCCUUUGAGGUACGCGUCACUCGUAGCAUUGUGCCUGUCUCCAACG<br>GCAAAGGAGAUAG |
| B | GCACAGUGCUAUGAGUGUCGCGACGGAGACGGUCGGGUCCAGAUAGGCCAGUCGACAAGGUC<br>UAUCUGUCGAGUAGAGUGUGGGCUCCGUCGCGUGCACAUUAGAGUCGCUGUAUGCCACAGUC<br>GACAAGUGGCGUACAGCGGCUCUAGUGUGCUGCACAGUGUCUGUGCGACUGCACCCAGUCGA<br>CAAGGGUGUAGUCGCAUAGACAUUGUGCUCACUCGUAGCAUUGUGCCUGUCUCCAGUCGACA<br>AGGAGAUAG |
| $\bar{B}$ | GCACAGUGCUAUGAGUGUCGCGACGGAGACGGUCGGGUCCAGAUAGGCCAUGGCGAAAGGUC<br>UAUCUGUCGAGUAGAGUGUGGGCUCCGUCGCGUGCACAUUAGAGUCGCUGUAUGACCCAUAG<br>CACAAGGGUCGUACAGCGGCUCUAGUGUGCUGCACAGUGUCUGUGCGACUGCACCCAACGAA<br>CAAGGGUGUAGUCGCAUAGACAUUGUGCUCACUCGUAGCAUUGUGCCUGUCUCCAACGGCAA<br>AGGAGAUAG |
| $A_{AP3}$ | GCACAGUGCUAUGAGUGUGCACGGGAUCCGAUCCCCAAUCGUGGCGUGUCGGCCUGCAUCGC<br>GAAAGCAGGCACUGGCGCCGGGAUCGGAUCCUGUGCUGCACAUUAGAGUCGCUGUAUGACCC<br>AUCGCGAAAGGGUCGUACAGCGGCUCUAGUGUGCUAGCUUCUGGACUGCGAUGGGAGCACGA<br>AACGUCGUGGCGCACGCGUGCCUCAGAAUCGCGAAAUCUGAGGCACGCGAUUGGGUGGGGAA<br>AGUCCUUAAAAGAGGGCCACCACAGAAGCUUCACUCGUAGCAUUGUGCCUGUCUCCAUCGCG<br>AAAGGAGAUAG |

**Supplementary Table 1: RNA sequences of nanostars used in this study.**

| Plasmid | DNA Sequences (5'-3') |
|---|---|
| A | CGAACAGAAAGTAATCGTATTGTACACGGCCGCATAATCGAAATTAATACGACTCACTATAG<br>CACAGTGCTATGAGTGTGCACGGGATCCGATCCCCAATCGTGGCGTGTCGGCCTGCATCGCG<br>AAAGCAGGCACTGGCGCCGGGATCGGATCCTGTGCTGCACATTAGAGTCGCTGTATGACCCA<br>TCGCGAAAGGGTCGTACAGCGGCTCTAGTGTGCTCGCGTGCCTCAGAGGACCTGTCACCATC<br>GCGAAAGGTGATAGGTCCTTTGAGGTACGCGTCACTCGTAGCATTGTGCCTGTCTCCATCGC<br>GAAAGGAGATAGCTAGCATAACCCCTTGGGGCCTCTAAACGGGTCTTGAGGGGTTTTTTG-P<br>1 |
| $\bar{A}$ | CGAACAGAAAGTAATCGTATTGTACACGGCCGCATAATCGAAATTAATACGACTCACTATAG<br>CACAGTGCTATGAGTGTGCACGGGATCCGATCCCCAATCGTGGCGTGTCGGCCTGCCGCCAA<br>CGAGCAGGCACTGGCGCCGGGATCGGATCCTGTGCTGCACATTAGAGTCGCTGTATGACCCA<br>TAGCACAAGGGTCGTACAGCGGCTCTAGTGTGCTCGCGTGCCTCAGAGGACCTGTCACCATG<br>GCGAAAGGTGATAGGTCCTTTGAGGTACGCGTCACTCGTAGCATTGTGCCTGTCTCCAACGG<br>CAAAGGAGATAGCTAGCATAACCCCTTGGGGCCTCTAAACGGGTCTTGAGGGGTTTTTTG-P<br>1 |
| B | CGAACAGAAAGTAATCGTATTGTACACGGCCGCATAATCGAAATTAATACGACTCACTATAG<br>CACAGTGCTATGAGTGTCGCGACGGAGACGGTCGGGTCCAGATAGGCCAGTCGACAAGGTCT<br>ATCTGTCGAGTAGAGTGTGGGCTCCGTCGCGTGCACATTAGAGTCGCTGTATGCCACAGTCG<br>ACAAGTGGCGTACAGCGGCTCTAGTGTGCTGCACAGTGTCTGTGCGACTGCACCCAGTCGAC<br>AAGGGTGTAGTCGCATAGACATTGTGCTCACTCGTAGCATTGTGCCTGTCTCCAGTCGACAA<br>GGAGATAGCTAGCATAACCCCTTGGGGCCTCTAAACGGGTCTTGAGGGGTTTTTTG-P1 |
| $\bar{B}$ | CGAACAGAAAGTAATCGTATTGTACACGGCCGCATAATCGAAATTAATACGACTCACTATAG<br>CACAGTGCTATGAGTGTCGCGACGGAGACGGTCGGGTCCAGATAGGCCATGGCGAAAGGTCT<br>ATCTGTCGAGTAGAGTGTGGGCTCCGTCGCGTGCACATTAGAGTCGCTGTATGACCCATAGC<br>ACAAGGGTCGTACAGCGGCTCTAGTGTGCTGCACAGTGTCTGTGCGACTGCACCCAACGAAC<br>AAGGGTGTAGTCGCATAGACATTGTGCTCACTCGTAGCATTGTGCCTGTCTCCAACGGCAAA<br>GGAGATAGCTAGCATAACCCCTTGGGGCCTCTAAACGGGTCTTGAGGGGTTTTTTG-P1 |
| AB | TAATACGACTCACTATAGCACAGTGCTATGAGTGTCGCGACGGAGACGGTCGGGTCCAGATA<br>GGCCAGTCGACAAGGTCTATCTGTCGAGTAGAGTGTGGGCTCCGTCGCGTGCACATTAGAGT<br>CGCTGTATGCCACAGTCGACAAGTGGCGTACAGCGGCTCTAGTGTGCTGCACAGTGTCTGTG<br>CGACTGCACCCAGTCGACAAGGGTGTAGTCGCATAGACATTGTGCTCACTCGTAGCATTGTG<br>CCTGTCTCCAGTCGACAAGGAGATAGAGATAACAGATACTTCGGTATCTGTTATCTGTTTTT<br>TTTCAACAGATAGCCGCGTTCGCGCGGCTATCTGTTTTTTTTCGAACAGAAAGTAATCGTAT<br>TGTACACGGCCGCATAATCGAAATTAATACGACTCACTATAGCACAGTGCTATGAGTGTGCA<br>CGGGATCCGATCCCCAATCGTGGCGTGTCGGCCTGCATCGCGAAAGCAGGCACTGGCGCCGG<br>GATCGGATCCTGTGCTGCACATTAGAGTCGCTGTATGACCCATCGCGAAAGGGTCGTACAGC<br>GGCTCTAGTGTGCTCGCGTGCCTCAGAGGACCTGTCACCATCGCGAAAGGTGATAGGTCCTT<br>TGAGGTACGCGTCACTCGTAGCATTGTGCCTGTCTCCATCGCGAAAGGAGATAGCTAGCATA<br>ACCCCTTGGGGCCTCTAAACGGGTCTTGAGGGGTTTTTTG-P1 |

| | |
|---|---|
| $A\bar{B}$ | TAATACGACTCACTATAGCACAGTGCTATGAGTGTCGCGACGGAGACGGTCGGGTCCAGATA<br>GGCCATGGCGAAAGGTCTATCTGTCGAGTAGAGTGTGGGCTCCGTCGCGTGCACATTAGAGT<br>CGCTGTATGACCCATAGCACAAGGGTCGTACAGCGGCTCTAGTGTGCTGCACAGTGTCTGTG<br>CGACTGCACCCAACGAACAAGGGTGTAGTCGCATAGACATTGTGCTCACTCGTAGCATTGTG<br>CCTGTCTCCAACGGCAAAGGAGATAGAGATAACAGATACTTCGGTATCTGTTATCTGTTTTT<br>TTTCAACAGATAGCCGCGTTCGCGCGGCTATCTGTTTTTTTTCGAACAGAAAGTAATCGTAT<br>TGTACACGGCCGCATAATCGAAATTAATACGACTCACTATAGCACAGTGCTATGAGTGTGCA<br>CGGGATCCGATCCCCAATCGTGGCGTGTCGGCCTGCATCGCGAAAGCAGGCACTGGCGCCGG<br>GATCGGATCCTGTGCTGCACATTAGAGTCGCTGTATGACCCATCGCGAAAGGGTCGTACAGC<br>GGCTCTAGTGTGCTCGCGTGCCTCAGAGGACCTGTCACCATCGCGAAAGGTGATAGGTCCTT<br>TGAGGTACGCGTCACTCGTAGCATTGTGCCTGTCTCCATCGCGAAAGGAGATAGCTAGCATA<br>ACCCCTTGGGGCCTCTAAACGGGTCTTGAGGGGTTTTTTG-P1 |
| $\bar{A}B$ | TAATACGACTCACTATAGCACAGTGCTATGAGTGTCGCGACGGAGACGGTCGGGTCCAGATA<br>GGCCAGTCGACAAGGTCTATCTGTCGAGTAGAGTGTGGGCTCCGTCGCGTGCACATTAGAGT<br>CGCTGTATGCCACAGTCGACAAGTGGCGTACAGCGGCTCTAGTGTGCTGCACAGTGTCTGTG<br>CGACTGCACCCAGTCGACAAGGGTGTAGTCGCATAGACATTGTGCTCACTCGTAGCATTGTG<br>CCTGTCTCCAGTCGACAAGGAGATAGAGATAACAGATACTTCGGTATCTGTTATCTGTTTTT<br>TTTCAACAGATAGCCGCGTTCGCGCGGCTATCTGTTTTTTTTCGAACAGAAAGTAATCGTAT<br>TGTACACGGCCGCATAATCGAAATTAATACGACTCACTATAGCACAGTGCTATGAGTGTGCA<br>CGGGATCCGATCCCCAATCGTGGCGTGTCGGCCTGCCGCCAACGAGCAGGCACTGGCGCCGG<br>GATCGGATCCTGTGCTGCACATTAGAGTCGCTGTATGACCCATAGCACAAGGGTCGTACAGC<br>GGCTCTAGTGTGCTCGCGTGCCTCAGAGGACCTGTCACCATGGCGAAAGGTGATAGGTCCTT<br>TGAGGTACGCGTCACTCGTAGCATTGTGCCTGTCTCCAACGGCAAAGGAGATAGCTAGCATA<br>ACCCCTTGGGGCCTCTAAACGGGTCTTGAGGGGTTTTTTG-P1 |
| $\bar{A}\bar{B}$ | TAATACGACTCACTATAGCACAGTGCTATGAGTGTCGCGACGGAGACGGTCGGGTCCAGATA<br>GGCCATGGCGAAAGGTCTATCTGTCGAGTAGAGTGTGGGCTCCGTCGCGTGCACATTAGAGT<br>CGCTGTATGACCCATAGCACAAGGGTCGTACAGCGGCTCTAGTGTGCTGCACAGTGTCTGTG<br>CGACTGCACCCAACGAACAAGGGTGTAGTCGCATAGACATTGTGCTCACTCGTAGCATTGTG<br>CCTGTCTCCAACGGCAAAGGAGATAGAGATAACAGATACTTCGGTATCTGTTATCTGTTTTT<br>TTTCAACAGATAGCCGCGTTCGCGCGGCTATCTGTTTTTTTTCGAACAGAAAGTAATCGTAT<br>TGTACACGGCCGCATAATCGAAATTAATACGACTCACTATAGCACAGTGCTATGAGTGTGCA<br>CGGGATCCGATCCCCAATCGTGGCGTGTCGGCCTGCCGCCAACGAGCAGGCACTGGCGCCGG<br>GATCGGATCCTGTGCTGCACATTAGAGTCGCTGTATGACCCATAGCACAAGGGTCGTACAGC<br>GGCTCTAGTGTGCTCGCGTGCCTCAGAGGACCTGTCACCATGGCGAAAGGTGATAGGTCCTT<br>TGAGGTACGCGTCACTCGTAGCATTGTGCCTGTCTCCAACGGCAAAGGAGATAGCTAGCATA<br>ACCCCTTGGGGCCTCTAAACGGGTCTTGAGGGGTTTTTTG-P1 |
| $A_{AP3}$ | TAATACGACTCACTATAGCACAGTGCTATGAGTGTGCACGGGATCCGATCCCCAATCGTGGC<br>GTGTCGGCCTGCATCGCGAAAGCAGGCACTGGCGCCGGGATCGGATCCTGTGCTGCACATTA<br>GAGTCGCTGTATGACCCATCGCGAAAGGGTCGTACAGCGGCTCTAGTGTGCTAGCTTCTGGA<br>CTGCGATGGGAGCACGAAACGTCGTGGCGCACGCGTGCCTCAGAATCGCGAAATCTGAGGCA<br>CGCGATTGGGTGGGGAAAGTCCTTAAAAGAGGGCCACCACAGAAGCTTCACTCGTAGCATTG<br>TGCCTGTCTCCATCGCGAAAGGAGATAGCTAGCATAACCCCTTGGGGCCTCTAAACGGGTCT<br>TGAGGGGTTTTTTG-P1 |

| | |
|---|---|
| GFP | TAATACGACTCACTATAGGGTCTAGAAATAATTTTGTTTAACTTTAAGAAGGAGGTATACAT<br>ATGGTGAGCAAGGGCGAGGAGCTGTTCACCGGGGTGGTGCCCATCCTGGTCGAGCTGGACGG<br>CGACGTAAACGGCCACAAGTTCAGCGTGTCCGGCGAGGGCGAGGGCGATGCCACCTACGGCA<br>AGCTGACCCTGAAGTTCATCTGCACCACCGGCAAGCTGCCCGTGCCCTGGCCCACCCTCGTG<br>ACCACCCTGACCTACGGCGTGCAGTGCTTCAGCCGCTACCCCGACCACATGAAGCAGCACGA<br>CTTCTTCAAGTCCGCCATGCCCGAAGGCTACGTCCAGGAGCGCACCATCTTCTTCAAGGACG<br>ACGGCAACTACAAGACCCGCGCCGAGGTGAAGTTCGAGGGCGACACCCTGGTGAACCGCATC<br>GAGCTGAAGGGCATCGACTTCAAGGAGGACGGCAACATCCTGGGGCACAAGCTGGAGTACAA<br>CTACAACAGCCACAACGTCTATATCATGGCCGACAAGCAGAAGAACGGCATCAAGGTGAACT<br>TCAAGATCCGCCACAACATCGAGGACGGCAGCGTGCAGCTCGCCGACCACTACCAGCAGAAC<br>ACCCCCATCGGCGACGGCCCCGTGCTGCTGCCCGACAACCACTACCTGAGCACCCAGTCCGC<br>CCTGAGCAAAGACCCCAACGAGAAGCGCGATCACATGGTCCTGCTGGAGTTCGTGACCGCCG<br>CCGGGATCACTCTCGGCATGGACGAGCTGTACAAGTAATGAGGATCCCGGGAATTCTCGAGT<br>AAGGTTAACCTGCAGGAGGCCTTTAATTAAGGTGGTGCGGCCGCGCTAGCGGTCCCGGGGGA<br>TCGATCCGGCTGCTAACAAAGCCCGAAAGGAAGCTGAGTTGGCTGCTGCCACCGCTGAGCAA<br>TAACTAGCATAACCCCTTGGGGCCTCTAAACGGGTCTTGAGGGGTTTTTTG-P2 |

**Supplementary Table 2**: **DNA sequences of nanostars used in this study.** P1: rest of the plasmid backbone with ampicillin resistance gene (Addgene #53137). P2: rest of the chloramphenicol resistance plasmid backbone (from pACYCDuet-1, Novagen).

| Dataset | Shapiro-Wilk | | Levene | | Kruskal-Wallis | | Nemenyi | | | | | | | |
|---|---|---|---|---|---|---|---|---|---|---|---|---|---|---|
| | W | p | F | p | H | p | AB | $\mathbf{A\bar{B}}$ | $\mathbf{\bar{A}B}$ | $\mathbf{\bar{A}\bar{B}}$ | $\mathbf{A_{AP3}}$+GFP | A+GFP | $\mathbf{A_{AP3}}$ | GFP |
| **AB** | 0.81 | **** | 75 | **** | 1500 | **** | - | **** | **** | **** | **** | 1.0 | 0.0030 | **** |
| $\mathbf{A\bar{B}}$ | 0.99 | 0.074 | | | | | **** | - | **** | **** | **** | **** | **** | **** |
| $\mathbf{\bar{A}B}$ | 0.77 | **** | | | | | **** | **** | - | **** | **** | 0.046 | 0.99 | 1.0 |
| $\mathbf{\bar{A}\bar{B}}$ | 0.99 | 0.0069 | | | | | **** | **** | **** | - | 0.0050 | **** | **** | **** |
| $\mathbf{A_{AP3}}$**+GFP** | 0.88 | **** | | | | | **** | **** | **** | 0.0050 | - | **** | **** | **** |
| **A+GFP** | 0.94 | **** | | | | | 1.0 | **** | 0.046 | **** | **** | - | 0.063 | 0.024 |
| $\mathbf{A_{AP3}}$ | 0.40 | **** | | | | | 0.0030 | **** | 0.99 | **** | **** | 0.063 | - | 1.0 |
| **GFP** | 0.87 | **** | | | | | **** | **** | 1.0 | **** | **** | 0.024 | 1.0 | - |

**Supplementary Table 3**: **Statistical analysis of $R^2$ across different experimental groups.** Normality was tested using the Shapiro–Wilk test, homogeneity of variances with Levene's test, and overall group differences with the Kruskal–Wallis test. Pairwise comparisons were conducted using the Nemenyi post-hoc test. $****$ indicates p-value lower than 0.0001.

| Dataset | Shapiro-Wilk | | Levene | | Kruskal-Wallis | | Nemenyi | | | |
|---|---|---|---|---|---|---|---|---|---|---|
| | **W** | **p** | **P** | **p** | **H** | **p** | **A** | **Ā** | **B** | **B̄** |
| **A** | 0.97 | **** | 250 | **** | 1400 | **** | - | **** | **** | **** |
| **Ā** | 0.98 | **** | | | | | **** | - | **** | **** |
| **B** | 0.88 | **** | | | | | **** | **** | - | 0.012 |
| **B̄** | 0.90 | **** | | | | | **** | **** | 0.012 | - |

**Supplementary Table 4: Statistical analysis of kurtosis across different experimental groups of single nanostar dataset in Pepper fluorescent channel.** Normality was tested using the Shapiro–Wilk test, homogeneity of variances with Levene's test, and overall group differences with the Kruskal–Wallis test. Pairwise comparisons were conducted using the Nemenyi post-hoc test. **** indicates p-value lower than 0.0001.

| Dataset | Shapiro-Wilk | | Levene | | Kruskal-Wallis | | Nemenyi | | | |
|---|---|---|---|---|---|---|---|---|---|---|
| | **W** | **p** | **P** | **p** | **H** | **p** | **A** | **Ā** | **B** | **B̄** |
| **A** | 0.97 | **** | 560 | **** | 1300 | **** | - | 0.29 | **** | **** |
| **Ā** | 0.97 | **** | | | | | 0.29 | - | **** | **** |
| **B** | 0.98 | **** | | | | | **** | **** | - | **** |
| **B̄** | 0.45 | **** | | | | | **** | **** | **** | - |

**Supplementary Table 5: Statistical analysis of kurtosis across different experimental groups of single nanostar dataset in Broccoli fluorescent channel.** Normality was tested using the Shapiro–Wilk test, homogeneity of variances with Levene's test, and overall group differences with the Kruskal–Wallis test. Pairwise comparisons were conducted using the Nemenyi post-hoc test. **** indicates p-value lower than 0.0001.

| Dataset | Shapiro-Wilk | | Levene | | Kruskal-Wallis | | Nemenyi | | | |
|---|---|---|---|---|---|---|---|---|---|---|
| | **W** | **p** | **P** | **p** | **H** | **p** | **AB** | **A$\bar{\text{B}}$** | **$\bar{\text{A}}$B** | **$\bar{\text{A}}\bar{\text{B}}$** |
| **AB** | 1.0 | 0.93 | 350 | **** | 1100 | **** | - | 0.00046 | **** | **** |
| **A$\bar{\text{B}}$** | 0.99 | 0.0076 | | | | | 0.00046 | - | **** | **** |
| **$\bar{\text{A}}$B** | 0.83 | **** | | | | | **** | **** | - | 0.71 |
| **$\bar{\text{A}}\bar{\text{B}}$** | 0.92 | **** | | | | | **** | **** | 0.71 | - |

**Supplementary Table 6**: **Statistical analysis of kurtosis across different experimental groups of cells expressing two nanostars in Pepper fluorescent channel.** Normality was tested using the Shapiro–Wilk test, homogeneity of variances with Levene's test, and overall group differences with the Kruskal–Wallis test. Pairwise comparisons were conducted using the Nemenyi post-hoc test. $****$ indicates p-value lower than 0.0001.

| Dataset | Shapiro-Wilk | | Levene | | Kruskal-Wallis | | Nemenyi | | | |
|---|---|---|---|---|---|---|---|---|---|---|
| | **W** | **p** | **P** | **p** | **H** | **p** | **AB** | **A$\bar{\mathbf{B}}$** | **$\bar{\mathbf{A}}$B** | **$\bar{\mathbf{A}}\bar{\mathbf{B}}$** |
| **AB** | 0.91 | **** | 430 | **** | 1200 | **** | - | **** | **** | **** |
| **A$\bar{\mathbf{B}}$** | 0.99 | 0.00065 | | | | | **** | - | **** | **** |
| **$\bar{\mathbf{A}}$B** | 0.97 | **** | | | | | **** | **** | - | **** |
| **$\bar{\mathbf{A}}\bar{\mathbf{B}}$** | 0.99 | 0.018 | | | | | **** | **** | **** | - |

**Supplementary Table 7: Statistical analysis of kurtosis across different experimental groups of cells expressing two nanostars in Broccoli fluorescent channel.** Normality was tested using the Shapiro–Wilk test, homogeneity of variances with Levene's test, and overall group differences with the Kruskal–Wallis test. Pairwise comparisons were conducted using the Nemenyi post-hoc test. **** indicates p-value lower than 0.0001.

| Dataset | Shapiro-Wilk | | Levene | | Kruskal-Wallis | | Nemenyi | | | |
|---|---|---|---|---|---|---|---|---|---|---|
| | **W** | **p** | **P** | **p** | **H** | **p** | **$A_{AP3}$+GFP** | **A+GFP** | **$A_{AP3}$** | **GFP** |
| **$A_{AP3}$+GFP** | 0.92 | **** | 66 | **** | 380 | **** | - | 0.79 | **** | **** |
| **A+GFP** | 0.95 | **** | | | | | 0.79 | - | **** | **** |
| **$A_{AP3}$** | 0.98 | 0.36 | | | | | **** | **** | - | **** |
| **GFP** | 0.83 | **** | | | | | **** | **** | **** | - |

**Supplementary Table 8: Statistical analysis of kurtosis across different experimental groups of protein capture in Pepper fluorescent channel.** Normality was tested using the Shapiro–Wilk test, homogeneity of variances with Levene's test, and overall group differences with the Kruskal–Wallis test. Pairwise comparisons were conducted using the Nemenyi post-hoc test. **** indicates p-value lower than 0.0001.

| Dataset | Shapiro-Wilk | | Levene | | Kruskal-Wallis | | Nemenyi | | | |
|---|---|---|---|---|---|---|---|---|---|---|
| | **W** | **p** | **P** | **p** | **H** | **p** | **$A_{AP3}$+GFP** | **A+GFP** | **$A_{AP3}$** | **GFP** |
| **$A_{AP3}$+GFP** | 0.93 | **** | 150 | **** | 440 | **** | - | **** | 0.0096 | **** |
| **A+GFP** | 0.73 | **** | | | | | **** | - | **** | 0.051 |
| **$A_{AP3}$** | 0.70 | **** | | | | | 0.0096 | **** | - | **** |
| **GFP** | 0.81 | **** | | | | | **** | 0.051 | **** | - |

**Supplementary Table 9**: **Statistical analysis of kurtosis across different experimental groups of protein capture in GFP fluorescent channel.** Normality was tested using the Shapiro–Wilk test, homogeneity of variances with Levene's test, and overall group differences with the Kruskal–Wallis test. Pairwise comparisons were conducted using the Nemenyi post-hoc test. **** indicates p-value lower than 0.0001.
